# Polysemanticity in human hippocampal neurons

**DOI:** 10.64898/2026.05.02.722435

**Authors:** Xinyuan Yan, Ji-An Li, Ana G. Chavez, Hanlin Zhu, Melissa Franch, Luca D. Kolibius, Rhiannon Cowan, James Belanger, Assia Chericoni, Taha Ismail, Kalman A. Katlowitz, Elizabeth A. Mickiewicz, Danika Paulo, Nisha Giridharan, Mohammed Hasen, Vaishnav Krishnan, Alica M. Goldman, Lu Lin, Atul Maheshwari, Eleonora Bartoli, Jay A. Hennig, Tomasz M. Frączek, Nicole R. Provenza, Shervin Rahimpour, Ben Shofty, Elliot Smith, Joshua Jacobs, Benjamin Y. Hayden, Sameer A. Sheth

## Abstract

As we perceive language, neurons in the brain represent the meanings of each word we hear. Inspired by recent advances in the mechanistic interpretability of large language models (LLMs), we hypothesized that hippocampal neurons utilize polysemanticity, a coding strategy wherein individual neurons represent multiple semantically distant features. Polysemanticity increases capacity, robustness, and scalability, and facilitates contextualization. We find that during podcast listening, hippocampal single neurons exhibit dense semantic codes characterized by multiple tuning peaks. These peaks, corresponding to the stimuli that most effectively drive the neurons, are not random; instead, they have overdispersed, isotropic geometry, consistent with interference-minimizing repulsive coding, although they lack grid-like hexagonal structure. Individual neuron responses are strongly modulated by context while population geometry remains stable and generalizable, allowing for pattern separation without sacrificing generalizability. Together, these results indicate that hippocampal neurons represent meaning using a distributed, dense, contextualized, polysemantic code consistent with the demands of superposition.

## INTRODUCTION

Just as neurons in the visual system are tuned for visual stimuli, neurons in the hippocampus are tuned for semantic features of the words we hear (Dijksterhuis et al., 2024; Franch et al., 2025; Goldstein et al., 2024; Katlowitz et al., 2025; Yan et al., 2026). Understanding the nature of that semantic tuning can help us understand how humans flexibly understand, learn, and deploy information. In large language models (LLMs), artificial neurons exhibit the property of **polysemanticity**, meaning that they respond to a range of seemingly unrelated stimuli (Bricken et al., 2023; Olah et al., 2020; Radford et al., 2019). For example, a single unit in a vision model may respond to either cats or cars; likewise, a unit in a language model may respond to either DNA code or Hebrew letters (Bricken et al., 2023; Olah et al., 2020). Polysemanticity has some similarity to mixed selectivity (Rigotti et al., 2013; Tye et al., 2024); however, polysemanticity collapses unrelated features onto a single axis (generally to increase coding capacity, known as superposition), whereas mixed selectivity creates non-linear conjunctions across distinct dimensions (generally to expand dimensionality, Fusi et al., 2016). (Here, features refer to fundamental, interpretable units of information, including the latent variables, semantic concepts, or sensory attributes, that the system needs to encode.)

Relative to monosemanticity, polysemantic tuning offers several advantages that are especially relevant for language representation. First, capacity: natural language relies on a vast, high-dimensional space of concepts, so it is critical to pack features into superposition to maximize representational efficiency with a limited number of units (Elhage et al., 2022; Olah et al., 2020). Second, polysemanticity provides a natural substrate for contextualization. Because a neuron’s firing can be conditioned upon specific co-occurring words, the local linguistic environment enables contingent differences in responding (Elhage et al., 2022). Polysemanticity, in other words, supports core hippocampal computations (Bakker et al., 2008; Cayco-Gajic & Silver, 2019; Leutgeb et al., 2007; Yassa & Stark, 2011): it facilitates pattern separation by projecting otherwise overlapping semantic inputs into distinct, nearly orthogonal population states, and it enables generalization by leveraging learned statistical dependencies between distributed features (Babadi & Sompolinsky, 2014; Bausch et al., 2026; Cayco-Gajic & Silver, 2019; Quiroga, 2020; Rey et al., 2025; Suthana et al., 2021). Moreover, because polysemantic codes distribute features across multiple shared units, they exhibit graceful degradation in the face of noise or network damage (Fusi et al., 2016; Hinton et al., 1984). This overlapping architecture also allows the system to flexibly scale to novel concepts without requiring dedicated, pre-allocated neurons (Bernardi et al., 2020; Rigotti et al., 2013).

The hypothesis that the human hippocampus utilizes polysemantic coding is consistent with its spatial codes. While hippocampal place cells are typically thought of as having single (i.e., monosemantic) response fields, recordings in large environments reveal multiple, unrelated firing fields that lack obvious topographical relationships (Chericoni et al., 2026; Eliav et al., 2021; Fenton et al., 2008; Mainali et al., 2025; Rich et al., 2014). And, while grid cells represent space with hexagonal multi-peaked fields (Hafting et al., 2005), this periodic structure breaks down into dense, non-hexagonal multi-field packing in higher-dimensional (3-D) or non-uniform environments (Ginosar et al., 2021; Grieves et al., 2021; Hayman et al., 2011). Given that the same coding principles used for place and grid cells apply to abstract concepts (Constantinescu et al., 2016; Knudsen & Wallis, 2021; Park et al., 2021), and given that semantic space is high-dimensional, we hypothesized that semantic tuning would show multiple, overdispersed, semantic modes. This hypothesis complements, but does not contradict, the theory of concept cells, which holds that neurons in medial temporal lobe (MTL) structures respond monosemantically to stimuli that evoke particular concepts (Quiroga, 2012), either in an all-or-none (Quiroga et al., 2005) or graded (Karkowski et al., 2025) manner. We further hypothesized that this code would be especially prominent in natural language listening, where contextualization is the rule, rather than the exception (Caucheteux & King, 2022; Goldstein et al., 2022; Katlowitz et al., 2026; Kumar et al., 2024).

## RESULTS

We collected responses of isolated single neurons in the hippocampus as patients listened to 47 minutes of speech (n=498 neurons, 15 patients; **Methods**, **Figure 1A**; Franch et al., 2026). All analyses were performed only on well-isolated single units, identified manually by trained lab members (e.g., **Figure 1B**/**C**). We defined word-evoked firing rates as beginning 80 ms after the onset of each word and lasting the duration of the word plus 40 ms. (All results shown below are robust to specific analysis windows used, see **Methods**). We independently replicated our major findings in a complementary dataset consisting of 46 minutes of dialogue (n = 217 neurons in the hippocampus, 6 patients, **Methods**, see **Supplementary Results**).

**Figure 1.**
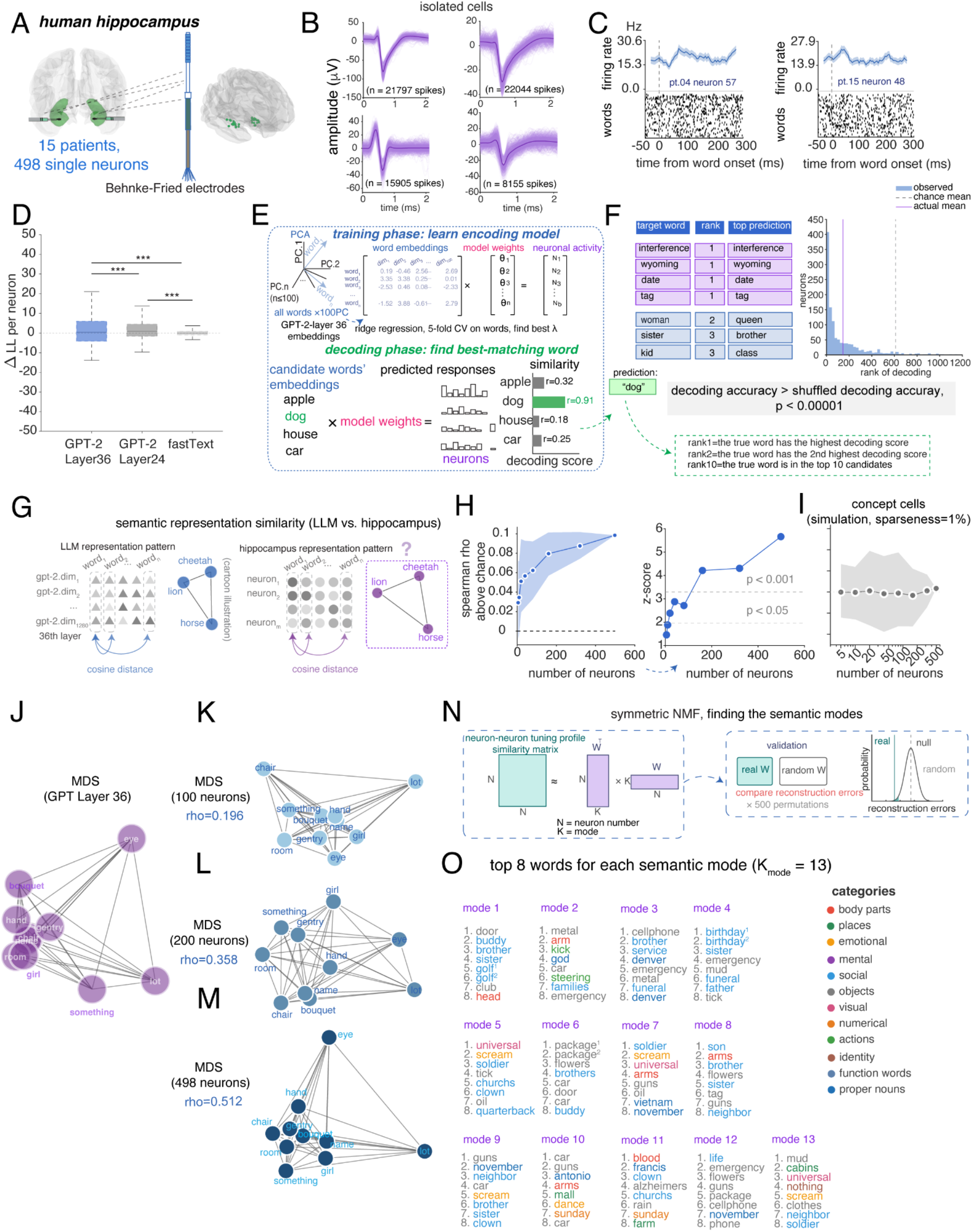
Hippocampal population recapitulates semantic embedding geometry. (A) Recording setup. Single-neuron recordings were obtained with Behnke-Fried microelectrodes in the hippocampus. (B) All analyses used well-isolated single units; four example cells shown. (C) Example PSTHs from two neurons showing firing rate modulation across words. (D) Encoding model performance. Cross-validated ΔLL per neuron (n = 498, Method). GPT-2 Layer 36 outperformed Layer 24 and fastText (paired t-tests, FDR-corrected; ***p < 0.001). Control analyses in **Figure S1A-C**. Boxplots show the distribution across neurons (n = 498) of cross-validated ΔLL per neuron (Methods); center line, median; box, 25th-75th percentiles. (E) Encoding-decoding framework. Top: ridge regression maps GPT-2 Layer-36 embeddings (100 PCs) to firing rates. Bottom: for each held-out word, the model ranks candidates by correlation with the observed population response (Methods). (F) Decoding performance. Left: examples of correct (purple) and semantically close (blue) recoveries. Right: recovery rank histogram is highly left-biased. (G) Representational similarity analysis (RSA) cartoon illustration. (H) RSA correlation scales monotonically with population size and does not saturate at N = 498. Shaded band: 95% CI of neuron-resampling distribution (Methods). Control analysis in **Figure S1D**. right panel: effect size of RSA at each population size. (I) Simulated concept cells showed no monotonically increasing RSA with population size and produced near-zero correlations with GPT-2 semantic geometry at all sparseness levels (here we show one example, all panels see **Figure S4**). (J-M) Multidimensional Scaling (MDS) visualizations of semantic geometry. Multidimensional scaling plots are shown for (J) the GPT-2 Layer-36 reference embedding, and neural populations of (K) 100, (L) 200, and (M) 498 neurons. Spearman rho between neural and LLM distance matrices increases. (N) Symmetric NMF (Non-negative matrix factorization) schematic (Methods). (O) Neural modules span multiple word categories. The top 8 words for each of the 13 neural modules are shown (Methods). We colored the top 8 words in each module by their semantic categories (Methods). Related to Table S1, Table S2, Figure S1, Figure S2, Figure S3, Figure S4.

### Hippocampal semantic coding during language listening is distributed

We quantified semantic tuning with ridge regression (Caucheteux & King, 2022; Franch et al., 2026; Jamali et al., 2024; Zada et al., 2025); we defined performance as cross-validated log-likelihood improvement over an intercept-only baseline (ΔLL; **Methods**). As predictors, we used the first 100 principal components of GPT-2 embeddings (Radford et al., 2019; Franch et al., 2026). Due to our interest in the encoding of word concepts, we analyzed only nouns; However, to confirm that the effects we report are robust, we then replicated our analyses using verbs and adjectives, all results support the same conclusions; see **Table S2**, **Figure S6** and **Supplementary Note 1**). Layer 36 (the final layer of GPT-2-large) yielded stronger encoding performance than Layers 24 or fastText (static word embeddings; Joulin et al., 2016, L36 vs L24: t(497) = 3.173, p<0.005; L36 vs fastText: t(497) = 5.336, p<0.001; L24 vs fastText: t(497) = 7.376, p<0.001; all comparisons remained significant after FDR-BH correction; also see **Figure S1A-C** for control analyses, using the first 50 PCs instead of the first 100 did not change the results). Layer 36 was therefore used for all subsequent analyses (**Figure 1D**). We next asked whether word identity could be recovered from population activity. Using an encoding-based reconstruction procedure (Tang et al., 2023; **Figure 1E**), many words were recovered exactly and nearly all others were semantically close (**Figure 1F**, right).

We asked whether the distances between words in neural population activity mirror the relational codes in LLM embedding space using representational similarity analysis (RSA; **Figure 1G**, Kriegeskorte et al., 2008; Nili et al., 2014). Indeed, the Spearman rank correlation between neural and LLM distance matrices is significantly above chance across the population (N = 498 neurons: rho = 0.092 vs. shuffle rho = -0.006; z = 5.66, p <0.001; **Figure 1H**). Moreover, this correlation increased monotonically with population size (Spearman’s rank correlation coefficient = 1 across eight population sizes, p < 0.001), reaching significance at ≥40 neurons. We repeated the RSA decimation analysis on simulated concept cells, in which encoding is known to be localized (Quiroga, 2012; see below and **Figure S3-S4**, **Methods**). Unlike real neurons, we find that simulated concept cells showed no monotonically increasing RSA with population size and produced near-zero correlations with GPT-2 semantic geometry at all sparseness levels (**Figure S4**; here we presented one representative example in **Figure 1I**). To rule out the possibility that the scaling merely reflects noise averaging across neurons with identical tuning, we constructed a control population by copying the single best-encoding neuron 498 times and adding independent noise matched to the empirical residuals (**Methods**). This copy-plus-noise population plateaued at moderate N, whereas the real population showed continued representational gains at large N, confirming that additional real neurons indeed contribute to semantic geometry (**Figure S1D**).

This finding confirms our previously published results by showing that the neural and semantic distances correspond monotonically, and extend them by showing the correlation increases with population size (**Figure 1J-M**; Franch et al., 2026). Note that the correspondence between neural and semantic representations also confirms earlier published results (e.g., Goldstein et al., 2024). This scaling behavior suggests the semantic encoding is distributed because it implies that individual neurons carry partial, overlapping information about many words that accumulate across the population (Kafashan et al., 2021).

### Lack of discrete functional categories of neurons with similar tuning

Response patterns of monosemantic neurons should roughly recapitulate the structure of semantic space, meaning that the set of stimuli that drive a given neuron (which we call its ***semantic modes***) should be semantically adjacent. This is true even for neurons that embody more abstract categories that cross basic animal/place/thing category lines, such as “Christmastime” (which would include reindeer, the North Pole, and candy canes). In either case a neuron’s semantic modes will necessarily have greater proximity than chance in some dimensions of embedding space. A polysemantic code, by contrast, predicts no such categorical grouping.

To quantify the internal coding structure within the population, we applied symmetric non-negative matrix factorization (NMF; Lee & Seung, 1999) to the neuron-by-neuron tuning similarity matrix (**Figure 1N**). NMF decomposes this similarity matrix into discrete components, each reflecting a distinct linear reweighting of the entire neural population. It produces neuronal “modules,” each representing a functionally coherent structure within the space of possible population tuning functions. We then ask whether these neurally-derived modules align with the structure of semantic space as determined by word models (fastText; **Methods**). This analysis revealed thirteen modules; this number was selected by computing reconstruction error and identifying the point of greatest curvature change in the error curve. We confirmed module validity by comparing its reconstruction error against a null distribution (permutation, p < 0.05). We defined semantic categories by performing k-means clustering on the fastText word embeddings to define discrete semantic categories (**Methods**). This method will find any categories that are latent in the training corpus, including both natural categories (e.g., animals) or abstract, even ad hoc, ones (e.g., “things you might encounter at the beach”). We found that NMF-driven modules do not map onto single fastText-derived semantic categories. Instead, each module recruits words from multiple semantic categories (**Figure 1O**; e.g., Modules 7 and 11 each span six different semantic categories).

To quantify this observation, we compared the mean pairwise semantic distance among each module’s top-scoring words (top 100 words) against a null distribution constructed by randomly sampling the same number of words (500 permutations). Only one of thirteen modules showed significantly clustered semantics (module 3, z=-2.03, p=0.020); eight modules were indistinguishable from chance, and four were significantly more dispersed than chance (modules 2, 4, 9, and 11; z = 1.83-7.10, all p ≤ 0.036), meaning their top words were drawn from more distant regions of semantic space than expected by chance. This overdispersion is critical because it cannot be explained by noise or by sparse responding. The number of dispersed modules exceeded that expected at α = 0.05 (4 of 13 vs. 0.65 expected; p = 0.003, one-sided binomial test), whereas the single clustered module was consistent with chance (1 of 13; p = 0.49). This bias toward overdispersion is a pattern we explore in more detail below.

Overall, the NMF analysis demonstrates that individual neurons have semantic modes that span a greater space than would be expected by chance even if their tuning surfaces were random samples of the larger semantic manifold. This pattern is replicated at the individual patient level (**Table S1** for nouns) and extends to verbs and adjectives (**Table S2**). As a result, this finding cannot be explained by the noise properties of the neurons, which would push the data towards the nulls.

### Non-sparse semantic coding during language listening

We next characterize the coding density of individual neurons in four different ways.

***Population sparseness*** quantifies how selectively a given word activates the neural population (**Figure 2A**; Rolls & Treves, 1997). Values near 1 indicate that only a single neuron responds, while values near 0 indicate uniform activation. We find population sparseness is moderate (mean±std=0.581±0.031). We compared this result to data generated from simulated concept cells that have metrics matching those of real concept cells (Waydo et al., 2006; **Figure S3A-E**; see **Methods**). Neurons in our dataset use a much denser code than the simulated concept cells (**Figure 2B**). Because Waydo et al. argue for a range of plausible sparseness parameters (0.20% to 1.0%), we confirmed this result using ten sparseness levels in this range (Wilcoxon rank-sum test, |z| ≥ 27.5, all p<0.001, all comparisons remained significant after FDR-BH correction).

**Figure 2.**
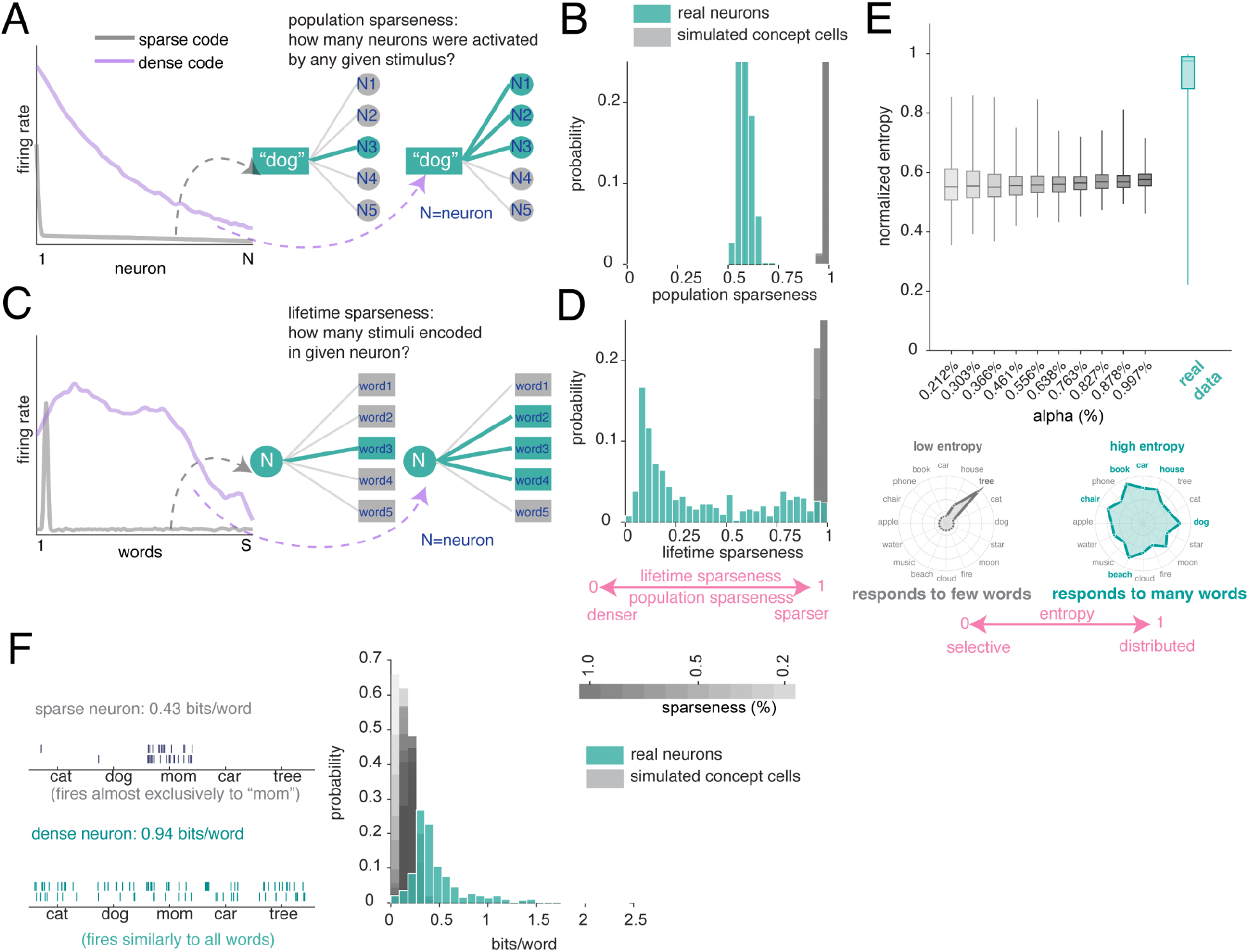
Hippocampal neurons use dense code to encode concepts. (A) Cartoon illustration of population sparseness: under a sparse code (gray), a stimulus activates few neurons; under a dense code (purple), it activates many. (B) Real neurons (teal) exhibit substantially lower population sparseness than simulated concept cells (gray). Simulation details in **Figure S3**. (C) Cartoon illustration of lifetime sparseness: under a sparse code, a neuron responds to few stimuli; under a dense code, it responds to many. (D) Real neurons exhibit substantially lower lifetime sparseness than simulated concept cells. (E) Normalized Shannon entropy (0 = maximally selective; 1 = maximally uniform) across simulated sparseness levels, with real neurons (teal, far right) clustering near the high-entropy end. Bottom: radar plots contrasting a low-entropy neuron (left), which concentrates responses on a few words, with a high-entropy neuron (right), which distributes responses broadly. (F) Information per word (bits/word; Skaggs et al., 1992, Methods). Left: cartoon contrasting a sparse neuron firing almost exclusively to one word (0.43 bits/word) with a dense neuron firing broadly (0.94 bits/word), illustrating that broadly tuned neurons can carry more information. Right: real neurons (teal) carry substantially more information per word than simulated concept cells (gray). Related to Figure S3, Figure S4, Figure S5, Figure S15, Supplementary Note 1 and Note 2.

***Lifetime sparseness*** quantifies how selectively a given neuron responds across words (**Figure 2C**): values near 1 indicate firing to a single word, while values near 0 indicate uniform responding (Rolls & Treves, 1997). Lifetime sparseness of neurons in the hippocampus (mean±std=0.369 ± 0.305) is considerably denser than that of the simulated concept cells (sparseness range: 0.20% to 1.0%, Wilcoxon rank-sum test, |z| ≥ 26.2, all p<0.001, all comparisons remained significant after FDR-BH correction; **Figure 2D**). We found 10 neurons with extreme high lifetime sparseness (i.e., responding to very few words) which met the criteria for defining concept cells, but such neurons responded to several semantically distant words (**Figure S5**), suggesting that their sparse firing reflects polysemantic tuning rather than narrow concept selectivity.

***Normalized Shannon entropy*** measures how uniformly a neuron distributes its firing across words (0 = maximally sparse, responding to a single word; 1 = maximally dense, responding uniformly to all words, **Methods,** Lehky et al., 2005). Neurons in our sample show relatively high entropy, that is, closer to responding to all words uniformly (mean±std = 0.910±0.130). Simulated concept cells produced much lower entropy values (sparseness range: 0.20% to 1.0%, Wilcoxon rank-sum test, |z| ≥ 25.1, all p<0.001, all comparisons remained significant after FDR-BH correction; **Figure 2E**).

A neuron could have high entropy (responds to many words) but carry very little information if its rate variations do not reliably distinguish between words. In other words, apparent density could be a by-product of high noise or variability. To test for this possibility, we computed the ***information per word*** (**Figure 2F**; **Methods**; Skaggs, Mcnaughton, et al., 1992). Neurons in our dataset convey substantially more information per word (0.464 ± 0.289) than simulated concept cells (sparseness range: 0.20% to 1.0%, Wilcoxon rank-sum test, |z| ≥ 25.1, all p<0.001, all comparisons remained significant after FDR-BH correction; **Figure 2F**). This result confirms that the broad tuning (high entropy) we observe is functionally meaningful and supports fine-grained discrimination among words. We replicated these results using narrower time windows, to control for the possibility that early responses to the next word may cause spurious coding density (**Figure S15A-D**). All four metrics yielded similar results for verbs, adjectives (**Supplementary Note 1**), and proper nouns (**Supplementary Note 2**).

We use simulated concept cells here to give an indication of what we would expect in a regime in which only concept cells were used. Note that this simulation matches our dataset in neuron count, word count, and word durations, while taking firing-rate and trial-to-trial variability parameters from the concept-cell literature (Quiroga et al., 2005; Waydo et al., 2006), so it indicates what a concept-cell regime would look like under our stimulus conditions. These results argue that a sparse concept cell-like code is not sufficient to describe neural coding for semantics in story listening. However, they do not argue that concept cells do not exist; instead, they argue for the existence of a dense code that may operate in parallel to a sparse one. Moreover, dense coding does not necessarily imply polysemanticity: a neuron could respond broadly within a single semantic category. We next asked whether the words that drive each neuron are semantically clustered.

### Hippocampal neurons are polysemantic

One way to visualize a neuron’s tuning function is to look at its predicted responses to a large corpus; we used a standard 50,000 noun corpus (Joulin et al., 2016). For illustration, we show the predicted response of three example neurons to these words (**Methods**). These show complex patterns of activation, with responses to a wide range of stimuli along with local clusters of particularly strong activation (i.e., semantic modes, **Figure 3A**). For comparison, we calculated estimated responses of simulated concept cells (see above and **Methods**), which have the expected narrow peaks corresponding to their preferred concepts (**Figure 3B**).

**Figure 3.**
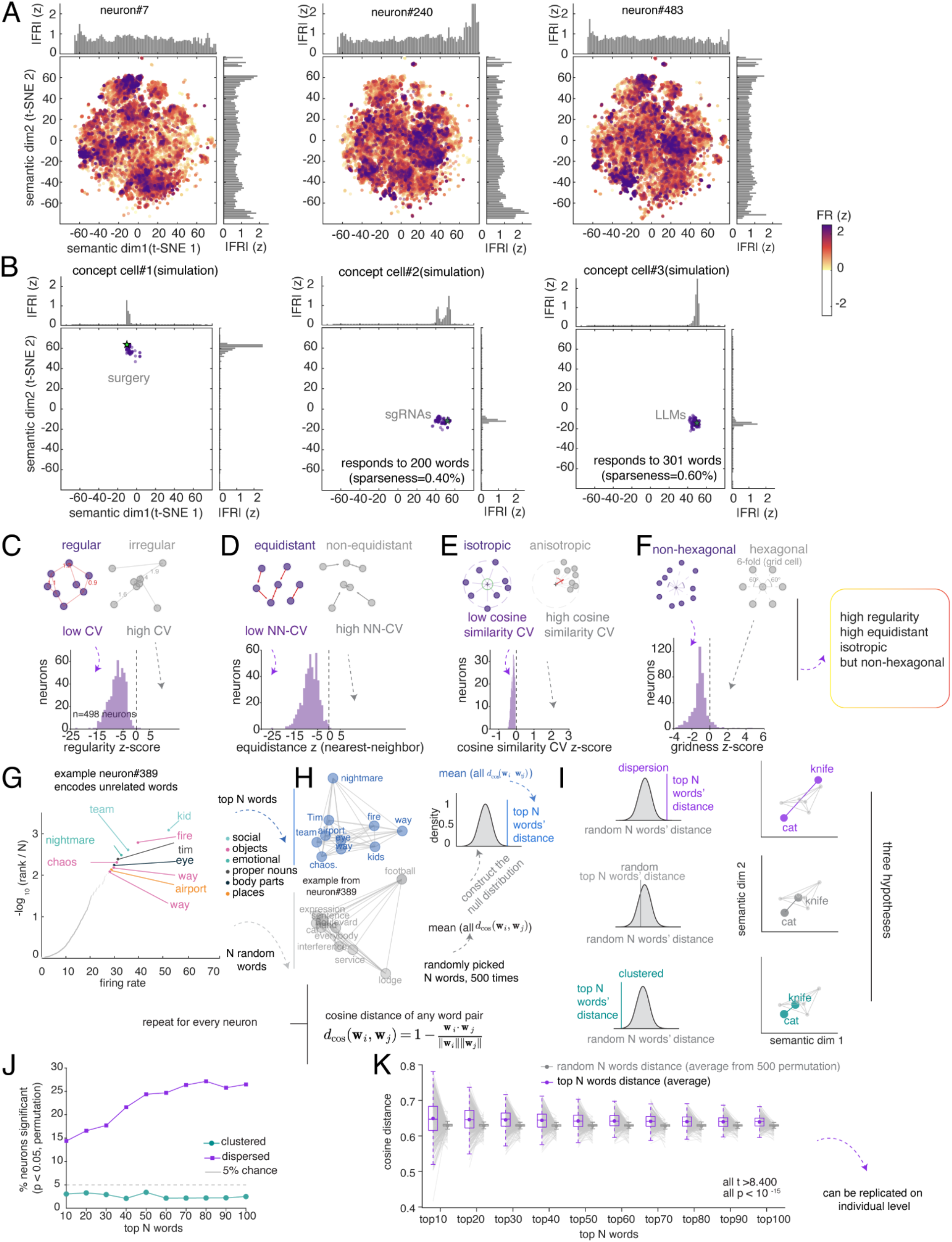
Single-neuron tuning landscapes are multi-moded and semantically overdispersed. (A) Single-neuron tuning landscapes. Predicted firing rates (z-scored) plotted on a t-SNE embedding of 50,000 nouns from an independent corpus (Methods). Three example neurons shown; dots colored by predicted firing rate (warm = high, white = low/zero); marginal histograms show mean |FR| per bin. (B) Simulated concept cells for comparison. Three concept cells (see Methods and Figure S3 for simulation details) at varying sparseness levels, displayed in the same t-SNE space. (C) Regularity. Top: schematic of regular vs. irregular arrangements. Bottom: z-score distribution shows tuning fields are more regularly spaced than chance. (D) Equidistance. Top: schematic. Bottom: nearest-neighbor distances are more uniform than chance. (E) Isotropy. Top: schematic. Bottom: tuning fields are approximately isotropically distributed. (F) Hexagonal symmetry. Top: schematic. Bottom: no evidence of six-fold symmetry, ruling out grid-like periodic tiling. (G) Example neuron (#389) encodes semantically distant words. Words ranked by firing rate with top 10 labeled by semantic category, spanning multiple unrelated categories. (H) Dispersion analysis schematic. Mean pairwise cosine distance among each neuron’s top N words compared against a null from randomly sampled words (Methods). (I) Schematic of possible outcomes: dispersed, random, or clustered. (J) Proportion of significantly dispersed neurons exceeds both chance (5%) and clustered neurons at all N values. (K) Top-N words are significantly more dispersed than random words at every N (all t > 8.4, p < 0.001). Control analyses see **Figure S7-S10**. Related to Figure S6, Figure S7, Figure S8, Figure S9, Figure S10, Figure S15 and Supplementary Note 3.

We identified individual modes using DBSCAN clustering of high-firing words in t-SNE 2D semantic plane (cf. Cao et al., 2025; **Methods**), where density-based clustering is more effective than in high-dimensional spaces due to distance concentration (Aggarwal et al., 2001). For each mode, we then computed its centroid as the firing-rate-weighted mean of all the words within that mode in the original high-dimensional space (**Methods**). All subsequent geometric analyses (except the gridness score, see below) are based on the mode centroids in the original embedding space, rather than the reduced t-SNE space.

We assessed the regularity of inter-mode spacing by computing the coefficient of variation (CV) of all pairwise distances between mode centroids (**Figure 3C**). The population showed significantly more ***regular*** spacing than chance (mean z ± SD=-7.274 ± 3.063, one-sample t-test vs. 0: p<0.001). (Note that here and in subsequent analyses, “chance” refers to the spacing obtained when the same high-firing words are randomly reshuffled into modes of the same sizes). We then tested whether nearest-neighbor distances between modes are uniform (**Figure 3D**). The population showed significantly more ***equidistant*** nearest neighbor spacing than chance (mean z±SD=-8.130 ±3.582, p<0.001; same reshuffling null, with nearest-neighbor distances recomputed on each reshuffle). We then examined the directional arrangement of modes by computing the CV of pairwise cosine similarities among direction vectors from the grand centroid to each mode centroid in the high-dimensional embedding space (**Figure 3E**). The population was significantly more ***isotropic*** than chance (mean z±SD= -0.200± 0.100, p<0.001; chance is the CV obtained for the same number of mode directions drawn uniformly at random in embedding space), indicating that modes were more likely distributed uniformly across all directions in semantic space rather than concentrated along a few preferred directions, as grid cells would. Indeed, we found no evidence of hexagonal symmetry in the hippocampal population’s semantic tuning maps (**Figure 3F**). Specifically, ***gridness*** scores (Hafting et al., 2005; Langston et al., 2010) were negative (mean z=-1.127±0.874, one-sample t-test vs. 0: p<0.001; chance is the gridness obtained after randomly reassigning the neuron’s predicted firing rates across words). In summary, hippocampal semantic representations are organized into regularly spaced, approximately equidistant, and isotropically distributed modes, without the periodic hexagonal symmetry characteristic of grid cells in two dimensions (equidistant and isotropic arrangements in high-dimensional spaces do not necessarily suggest hexagonal symmetry in two dimensions). This structure is reminiscent of non-hexagonal grid cell geometry in high dimensional and non-uniform environments (Ginosar et al., 2021, 2023; Grieves et al., 2021; Krupic et al., 2015). Note that the above results are robustly observed with a range of parameters for both t-SNE and DBSCAN (p<0.001 in each of 20 combinations of t-SNE and DBSCAN parameters, see **Methods**).

We also note that PCA produces dense components, while ridge regression penalizes large weights without eliminating them. It is possible therefore that this pipeline may artifactually favor distributed codes by construction. To test for this possibility, we conducted a control analysis using a sparse-dominated alternative: non-negative matrix factorization (NMF, Lee & Seung, 1999) for embedding decomposition paired with elastic net regression (L1 ratio=0.9, see **Methods**), which promotes sparsity by forcing the coefficients of less relevant features to exactly zero. None of these results were affected by the choice of encoding model.

The fact that geometry is isotropic suggests that neuronal responses may be more semantically dispersed (***overdispersion***) than if their optimal driving stimuli were random (example neuron, **Figure 3G**). To test for overdispersion, for each neuron we identified the top N preferred nouns (by firing rate), computed their mean pairwise cosine distance in GPT-2 embedding space, and compared this distribution against a null distribution constructed from random nouns (**Figure 3H-I**). Across the population, the proportion of neurons whose preferred nouns are significantly more dispersed than chance (specifically, across a range of Ns, 14.4%-26.5%) substantially exceeds both the 5% false positive rate and the proportion showing significant clustering (i.e., across a range of Ns, 3.0%-2.5%). This pattern is not an artifact of the N we chose, as it holds across a range of Ns (specifically, all values of N from 10 to 100, **Figure 3J**). Quantitatively, the mean pairwise distance among each neuron’s top N words is greater than the null expectation at every top N tested (paired-t test, all t>8.400, all p<0.001, all comparisons remained significant after FDR-BH correction, **Figure 3K**; df ranged from 441 to 493, reflecting the number of valid neurons at each N, see **Methods**; similar results held for verbs and adjectives, see **Figure S6**). This result can be replicated at the patient level (**Figure S7**). Note that these results are unlikely to be explained by factors like noisy and/or sparse firing (**Supplementary Note 3**). Most critically, noisy/sparse firing would push individual neuron semantic maps in the direction of randomness, but would not produce the overdispersed pattern we observe. Nor would they produce the isotropy and equidistance patterns we observe in the relative positioning of semantic modes.

This finding of overdispersed semantic modes provides a robust unsupervised method for demonstrating polysemanticity; in other words, the neurons’ tuning surfaces can be described as a collection of modes that tile the semantic space more than would be expected if they were unimodal, and, moreover, more than would be expected if their tuning surfaces were random. Overdispersion does not depend on the choice of embedding space: the same pattern holds in GPT-2-large Layer 24, GPT-2-XL (Layers 32 and 48), and Gemma (Layers 29 and 42) (**Figure S8**), indicating that it reflects a property of the neural code rather than of any single language model.

One possible alternative explanation for these findings is that semantic overdispersion among each neuron’s preferred words might be driven by noise in the tail of the firing rate distribution rather than by the neuron’s strongest responses. To test for this possibility, we next performed a firing-rate-weighted control analysis (**Methods**). Each word pair is weighted by the product of their firing rates to ensure that the highest-firing words dominate the metric. This control analysis showed nearly identical results (paired-T test, all t>7.870, all p<0.001, all comparisons remained significant after FDR-BH correction; **Figure S9**). Consistent with this, firing declined gradually along each neuron’s preference ranking (**Figure S10**), so the top-N sets comprise nouns that drive each neuron at a substantial fraction of its peak rate rather than a sparse peak padded with noise. We replicated these results using narrower time windows to control for the possibility that early responses to the next word may cause spurious polysemanticity (**Figure S15E-F**). Of note, we performed a model-free split-half analysis of measured firing rates. Each neuron’s semantic tuning replicated across independent data halves, and its validated top-driving semantic regions were significantly more dispersed than random sets. Polysemantic overdispersion is therefore a property of the observed responses, not of the encoding model (see **Table S3** and **Methods**).

### Context reshapes single-neuron tuning while preserving population geometry

We next asked whether hippocampal semantic tuning functions are modulated by speaker-identity context (i.e., story context; **Figure 4A**). For each neuron, we computed split-half tuning vectors from random halves (repeated 50 times with different random splits) of the same story and from different stories. Across-story cosine distances were greater than within-story distances (paired t-test, t(497) = 19.813, p<0.001; **Figure 4B**), indicating that individual neurons adopt different semantic tuning depending on story. This change was individually significant in all fifteen patients (**Figure 4C**). This effect is not explainable by representational drift, as across-story tuning distance did not correlate with temporal separation between stories (Spearman rho=-0.074, p=0.794; **Figure S11**). Note that speaker identity is confounded here with a range of context-driving factors, such as volume, timbre, pitch, and so on. However, our central claim is that context, regardless of its form, drives changes in semantic tuning. As such, the specific details of which features drive the change are irrelevant.

**Figure 4.**
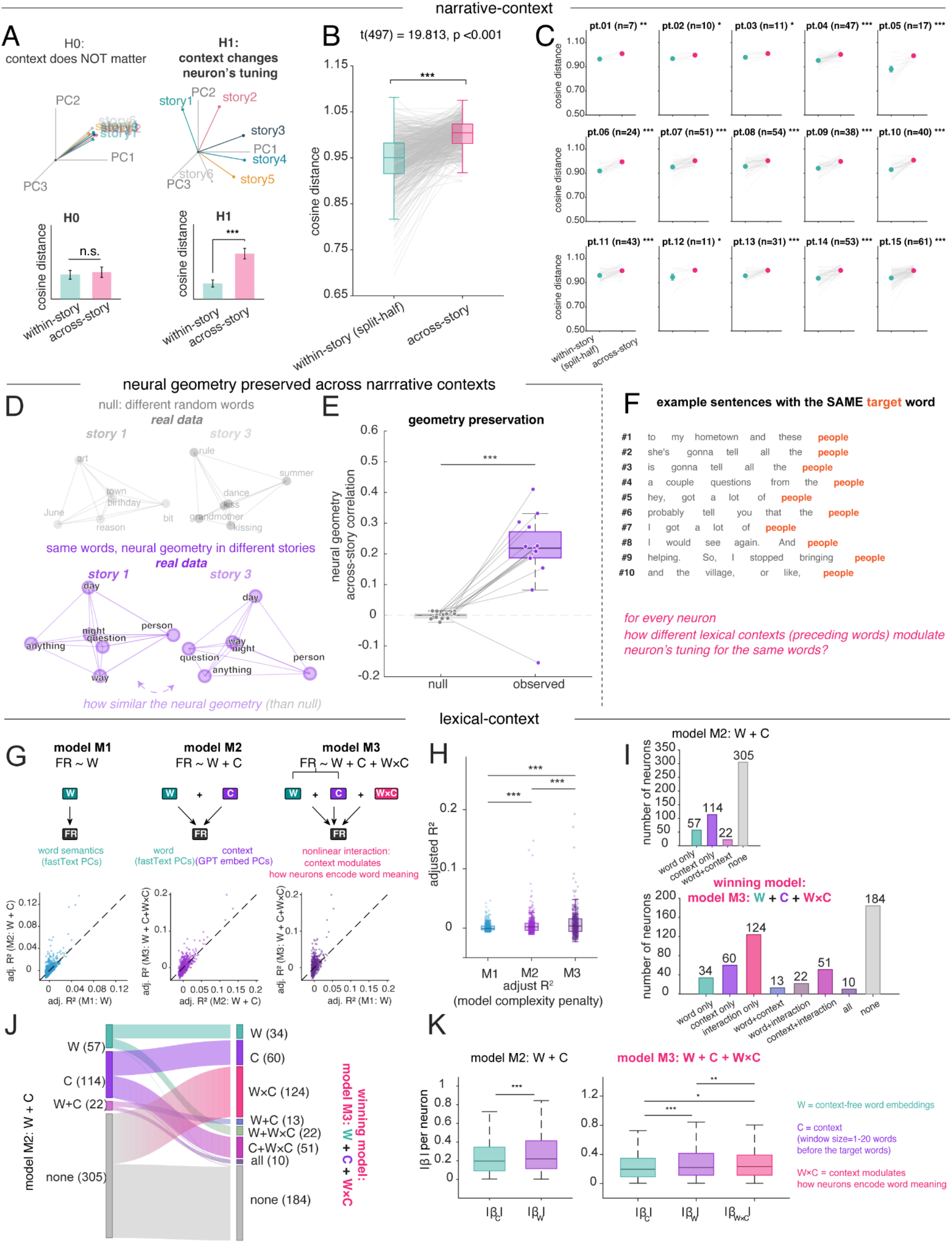
Story context and lexical context modulate semantic representations. (A) Static vs. contextualized hypotheses for semantic tuning across stories. Control analysis in **Figure S11**. (B) Semantic tuning is less stable across stories than within stories (story context), indicating context-dependent tuning profiles. (C) The story context modulating semantic tuning effect replicates across all 15 patients individually. (D) Neural geometry of shared nouns preserved across stories. For each story pair, RDMs of shared nouns are computed separately and compared via Spearman correlation. Null uses non-matching words (Methods). (E) RSA correlation for shared words (purple) exceeds the null (gray) across all 15 story pairs, demonstrating preserved geometric relationships regardless of story context. (F) The same target word (“people”) appears in 10 different lexical contexts across the podcast. (G) Model comparison framework. M1: firing rate ∼ lexical embedding (W). M2: ∼ W + context embeddings (C). M3: ∼ W + C + W×C interaction. Scatter plots show neuron-by-neuron adjusted R² comparisons, where W is the lexical embedding. (H) M3 is the winning model, confirming that both context and the word-by-context interaction contribute explanatory variance. (I) Neuron classification by significant predictors. M2: 57 neurons with significant word effects only, 114 with significant context effects only, 22 with both significant, 305 with neither. M3: 124 interaction-specific neurons emerge from the previously unclassified group; 207 total (41.6%) show significant interaction effects. (J) Sankey plot showing that interaction-only neurons in M3 derive entirely from unclassified neurons in M2. (K) Magnitude hierarchy: |β_C| > |β_W×C| ≥ |β_W|, stable across context window sizes (**Figure S12**). *Notes:* Panels G, H, K show results for context window size = 20 words. Other window sizes in **Figure S12**. Control analyses see **Figure S13**. In panel E and K, boxes show the distribution across story pairs (E) or neurons (K); center line, median; box, 25th-75th percentiles. Related to Figure S11, Figure S12, Figure S13

Despite these context-dependent shifts in individual neurons’ tuning, the population-level geometric arrangement of word representations was preserved across stories. For each pair of stories 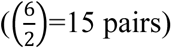, we identified nouns that appeared in both stories and computed the pairwise distance (representational dissimilarity matrix, RDM) of these shared words separately within each story (**Figure 4D**; **Methods**). The correlation between story-specific RDMs was significantly higher than null distributions constructed from mismatched random words across all 15 story pairs (observed r=0.206 ±0.126, null r =-0.001 ± 0.010; Wilcoxon signed-rank, p<0.001) (**Figure 4E**). In other words, sets of nouns maintain similar geometric relationships to each other regardless of their story context, even as neurons shift their tuning across stories.

We next examined how local lexical context affects semantic tuning (**Figure 4F**). We compared three nested linear models (**Figure 4G**; **Methods**): M1 predicted firing rates from static fastText word embeddings alone (lexical embedding, no context); M2 added contextual GPT-2 embeddings of the preceding 1-20 words (i.e., context embeddings), allowing context to independently shift firing rates without changing how the neurons responds to word meaning; M3 further included the word-by-context interaction, allowing the context to directly modulate the way that neurons encode word meaning. To avoid arbitrarily selecting a single context window, we varied the number of preceding words from 1 to 20 and tested each window size independently. Model comparison using adjusted R^2^ (which penalizes model complexity) revealed that M2 significantly outperformed M1 at all 20 context window sizes (paired t-tests across 498 neurons, t(497) = 5.74∼8.11, all p < 0.001 after FDR-BH correction; **Figure S12A**), and M3 further improved over M2 at 19 of 20 window sizes (t(497) = 2.74∼4.90, all p < 0.007 after FDR-BH correction, except window 1; **Figure S12A**), confirming that both context and its nonlinear interaction with lexical embedding contribute explanatory variance beyond lexical embedding alone (**Figure 4H**).

We further classified neurons by which predictors reached significance (**Figure 4I**). Under the additive model (M2), neurons are divided into word-only (n = 57), context-only (n = 114), and word-plus-context (n = 22) groups, with 305 neurons unclassified. When the interaction term was included (M3), a substantial population of interaction-specific neurons emerged (n = 124/498, 24.9%). These interaction-only (WxC) neurons are all from the unclassified neurons in M2, meaning that linear regression would define these as unmodified by either semantics or context. In total, 207 neurons (41.6%) involved the interaction term in M3, drawn from every category in M2 (**Figure 4J**). Together, these results demonstrate that apparent main effects of lexical embedding and context in M2 are better explained as a word-by-context interaction in M3. In other words, local lexical context does not simply shift firing rates up or down in a gain-like process, but changes *how* neurons encode word meaning.

We next examined the relative magnitude of each predictor. We found that context embeddings (the GPT-2 representation of the preceding words) carried significantly larger standardized regression coefficients than lexical embedding in both M2 and M3 at all window sizes (all p<0.002 after FDR-BH correction; **Figure S12B-C**). In M3, the interaction term was comparable to or larger than lexical embedding (**Figure S12D**) and consistently smaller than context main effects (**Figure S12E**, all p<0.05 after FDR-BH correction, except window2), yielding a magnitude hierarchy of |β_C_| > |β_W×C_| ≥ |β_W_| (**Figure 4K**). Thus, local lexical context is the dominant driver of hippocampal firing rate variance, while its nonlinear modulation constitutes a secondary but substantial influence that exceeds the contribution of lexical embedding alone.

To assess the prevalence of nonlinear interaction effects, we tested all two-way (W_i_ × C_j_; 100 terms) and three-way (W_i_ × W_j_ × C_k_ and W_i_ × C_j_ × C_k_; 900 terms) interaction terms constructed from the first 10 principal components of the word (explaining 34.84% variance) and context embedding spaces (explaining 20.97% variance), yielding 1000 interaction models per neuron. For each neuron, we fit a simple regression of firing rate onto each interaction term independently and assessed significance at α = 0.05. Under the null hypothesis of no interaction, the number of significant terms per neuron follows a binomial (1000, 0.05) distribution, with an expected value of 50 and a one-tailed significance threshold of 63. Across context window sizes 1-20, interactions per neuron ranged from 59.1 to 65.0, consistently exceeding chance. Overall, 39.8%-47.8% of neurons (198-238 out of 498 neurons) exhibited more interaction than chance (per-neuron binomial test, p < 0.05; versus 5% of neurons expected, population-level binomial test, all p < 0.001 after FDR-BH correction across the 20 window sizes; 35.5%-45.2% of neurons remained above chance when the per-neuron test was FDR-corrected across neurons). These results therefore indicate that hippocampal neurons do not make use of a stable amodal semantic code; instead, the code exhibits a high degree of mixed selectivity (cf. Posani et al., 2026).

### Polysemantic codes enable pattern separation without sacrificing generalizability

Pattern separation is the process by which the hippocampus transforms overlapping inputs into distinct, non-interfering neural representations (Yassa & Stark, 2011). We next asked whether and how hippocampus maps *identical* lexical inputs onto *distinct* neural representations when they occur in different semantic contexts (**Figure 5A-B**). Identical word inputs should evoke distinct population responses in different contexts, and the magnitude of this shift should scale with the distance between those contexts (Ethayarajh, 2019). For each noun appearing at least five times across the podcast, we computed pairwise distances among its occurrences in lexical context space (defined as the mean population firing rate vector across the 1-20 words immediately preceding each occurrence) and target word space (population response to the noun itself; **Figure 5C**). Across the population, observed context-repeated word geometry coupling (averaged across all repeated words with occurrence >=5) significantly exceeded a random-context null baseline across all context window sizes (1 to 20 preceding words used to define the context vector; mean r = 0.110-0.183; permutation test, all p < 0.05 after FDR-BH correction across window sizes; **Figure 5D**; **Methods;** see **Figure S14** for control analysis). This finding goes beyond showing that context shifts responses (**Figure 4D-E**): it demonstrates that the population maps contextual dissimilarities onto representational dissimilarity in a graded manner. This graded coupling between contextual distance and representational distance shows how vectorial semantic representations can implement semantic pattern separation.

**Figure 5.**
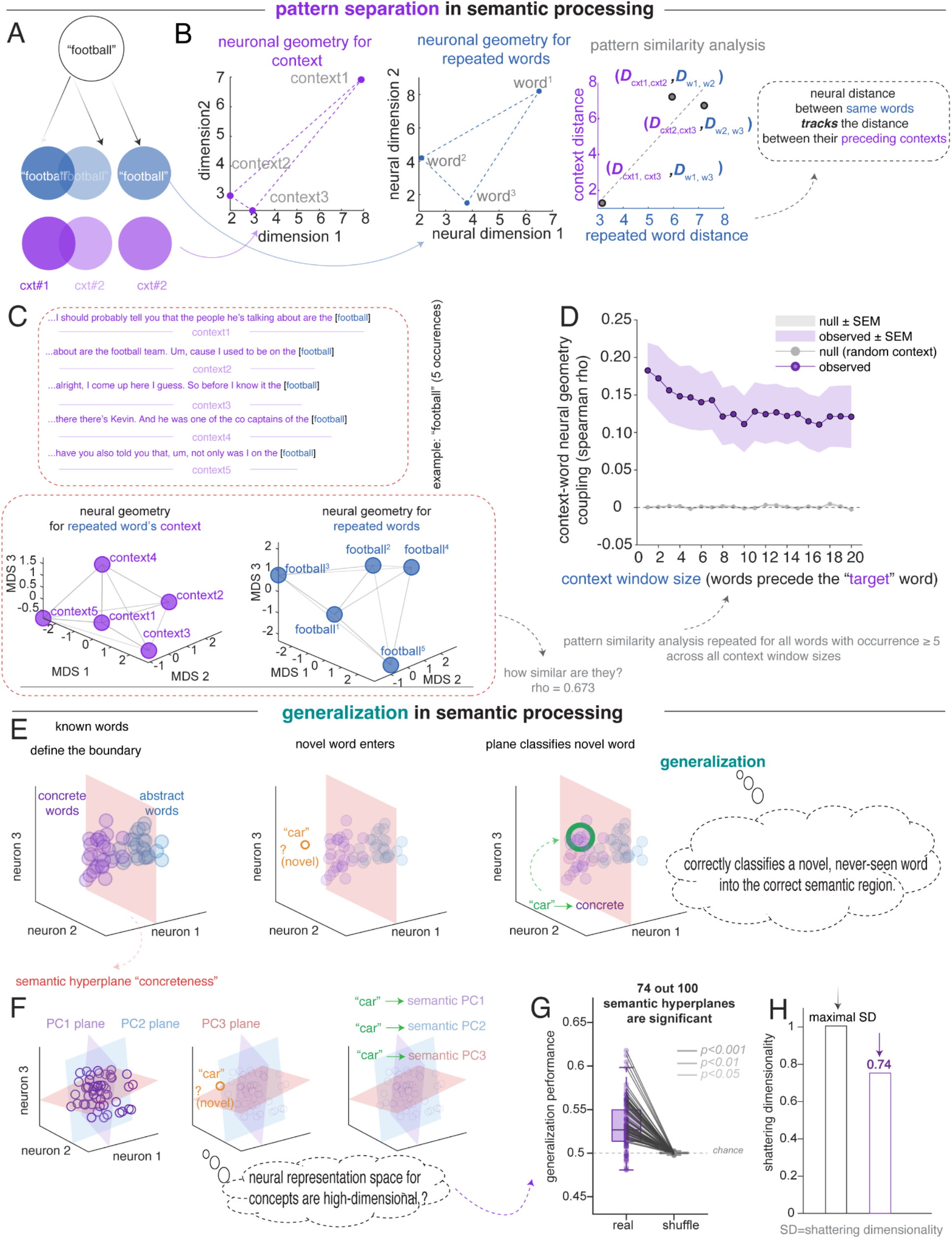
Hippocampal population supports context-driven pattern separation and preserves generalizability. (A) Schematic of context-driven pattern separation. The same word (“football”) appears in multiple lexical contexts; occurrences preceded by dissimilar contexts should elicit dissimilar population responses. (B) Context-word geometry coupling schematic. Pairwise distances among occurrences of a repeated word are computed in lexical context space (left) and target word space (middle). Pattern separation predicts that representational distance tracks contextual distance (right). (C) Example: “football” appears in 5 contexts. Cosine distances in lexical context space and target word space show correlated geometry (Spearman rho = 0.673). (D) Context-word coupling across windows of 1-20 preceding words. Observed coupling (purple) significantly exceeds random-context null (gray) at all window sizes (Methods). (E) Generalization schematic. Population responses to known words define hyperplanes partitioning neural space. A novel word is classified by which side its response falls, without prior exposure. (F) Multiple semantic hyperplanes partition neural space along different semantic contrasts, enabling multi-attribute generalization. (G) Generalization performance across 100 semantic dimensions. Real performance (purple) significantly exceeds shuffle baseline (gray); 74 of 100 dimensions reach significance (p < 0.05, permutation test, FDR-corrected; Methods). (H) Shattering dimensionality, computed as the proportion of semantic dimensions classifiable above chance. The observed value of 0.74 indicates that the population supports linear readout along three-quarters of tested dimensions, confirming a high-dimensional distributed code. Related to Figure S14.

To be maximally usable, pattern separation needs to preserve some degree of generalizability, even in the face of varying context. In language, the brain can map novel or incomplete inputs onto the appropriate semantic region defined by previously learned word relationships (**Figure 5E-F**). Concept cells are excellent at generalization in the sense that they can represent the same stimulus regardless of modality or context (Quiroga, 2012; Rey et al., 2025). Polysemantic neurons can also achieve generalizability, albeit in a different manner. By leveraging the superposition of unrelated features, a single polysemantic neuron can participate in multiple attractor states, allowing the broader network to collapse onto the correct semantic manifold as contextual cues accumulate (Elhage et al., 2022; Whittington et al., 2020).

We reasoned that if the hippocampal population maintains a high-dimensional, structured semantic geometry that positions words along many semantic dimensions simultaneously, then this geometric structure itself should support generalization. That is, a semantic boundary defined by one set of words should correctly classify words never used to define that boundary. We borrowed the framework of cross-condition generalization performance (CCGP, Bernardi et al., 2020; Courellis et al., 2024). For each of the first 100 principal components of GPT-2 Layer-36 embedding space, we split words into high versus low groups, trained a linear SVM on 70% of the words, and tested on the remaining 30% (Bernardi et al., 2020). Because the classifier never sees the test words during training, above-chance accuracy requires that the population geometry encodes the semantic dimension in an abstract format (**Figure 5E-F**, **Methods**). Generalization performance exceeded both chance and the shuffled baseline; 74 out of 100 semantic dimensions reached significance (all p<0.05 after FDR-BH correction, **Figure 5G**), corresponding to a shattering dimensionality (SD) of 0.74 (**Figure 5H**).

This high-dimensional geometry simultaneously supports pattern separation without sacrificing generalizability. That is, when a novel word is encountered, the neural ensemble can use many different semantic hyperplanes learned from the population responses of other words to correctly position the novel word. For example, when we hear “*dog*”, this word can be classified into the animacy plane representing “*animate”* or *“inanimate*”, and the affective plane representing “*warm*” or “*cool*” (exemplifying generalizability). At the same time, the abundance of distinguishing dimensions ensures that words sharing one semantic feature can nonetheless be differentiated along others, allowing the same word to be interpreted differently depending on the context (exemplifying separation).

## DISCUSSION

When processing natural speech, hippocampal neurons robustly encode semantic meaning (Franch et al., 2026); here we show that the set of stimuli that most effectively drive a given neuron form an overdispersed, isotropic, and equidistant structure within the neural embedding space. This polysemanticity closely parallels the properties of neurons in LLMs, where networks utilize superposition to maximize information capacity within a finite-dimensional space, forcing individual units to represent multiple disparate features (Bricken et al., 2023; Elhage et al., 2022). Human language is an ideal use case for polysemanticity because the set of concepts available to language is both very high dimensional and sparse, so the hippocampus can project two words into the same neural dimensions with low risk. By maintaining an isotropic and equidistant structure, the network further ensures that these overlapping representations remain as close as possible to orthogonal, reducing the chance of catastrophic interference and allowing the broader neural population to unambiguously decode meaning (Olah et al., 2017, 2020). These results do not, however, imply that the brain, generally speaking, uses the same computational principles as LLMs. Indeed, even if representations are similar in both brains and LLMs, this fact does not imply that they develop through similar principles.

Polysemantic tuning in hippocampal representations of meaning does not invalidate the theory of concept cells; instead, it suggests they may be part of a larger continuum of semantic coding. While we found very little evidence for concept cells in our data, we did not perform the types of careful searching that have proven critical for finding concept cells (Quiroga et al., 2005); we may therefore have simply missed them. Alternatively, story listening may push the hippocampus from a localized to a distributed code. Indeed, superposition theory predicts a phase transition governed by feature sparsity (Elhage et al., 2022). Classic concept cell protocols (experimental paradigms), which involve discrete concepts presented without context, naturally favor a dedicated-neuron regime (Quiroga, 2012), whereas naturalistic speech, with thousands of rapidly and transiently appearing contextualized concepts, favors polysemanticity. So both regimes may coexist within the same population, depending on the statistical structure of the input.

Most physiological studies report that conjunctive coding of stimulus and context is rare in human MTL, and that as a result pattern separation is limited (Bausch et al., 2026) or absent (Quiroga, 2020; Rey et al., 2025). In contrast, our results, obtained during language listening, revealed robust conjunctive coding (i.e., nonlinear mixing of context and semantics). This non-linear mixed selectivity is the key ingredient that allows for contextualization and efficient neural computation (Johnston et al., 2020) and its cousin, pattern separation (Barak et al., 2013; Bernardi et al., 2020; Fusi et al., 2016). It also increases coding capacity and fidelity (Johnston et al., 2020). Moreover, these mixed-selective patterns directly recapitulate those observed in hippocampal neural populations in a range of contexts (Bernardi et al., 2020; Boyle et al., 2024; Chericoni et al., 2026; Courellis et al., 2024; Tang et al., 2023). Given the strong need for contextualization in language, such conjunctive coding is invaluable. This discrepancy, then, may reflect that hippocampal pattern separation is latent under discrete, static paradigms and actively recruited by the computational demands of continuous language.

Overall, our observations provide a bridge between semantic representation and classical models of hippocampal pattern separation (Bakker et al., 2008; Leutgeb et al., 2007; McClelland et al., 1995; Yassa & Stark, 2011). Traditional frameworks emphasizing sparse, amodal concept cells pose a computational paradox in this regard: their monosemantic rigidity lacks the high-dimensional capacity required to decorrelate closely overlapping lexical inputs (Quiroga, 2020). This limitation seemingly contradicts the amply demonstrated role of the hippocampus in contextualization and pattern separation (Suthana et al., 2021). Our results suggest that such sparse identity-coding is complemented by a polysemantic architecture that supports pattern separation (Cayco-Gajic & Silver, 2019; Rigotti et al., 2013). Indeed, our findings in the hippocampus suggest a conceptual unification between the psychology of pattern separation, which has typically been applied to relational memory, and language. It is not clear how these results would differ, if at all, if we analyzed data from classic language network regions (Fedorenko et al., 2024). However, reports of contextual modulation in single unit recordings in language regions suggest some of these principles may be universal (Cai et al., 2026; Jamali et al., 2024).

An important consideration is whether the observed polysemanticity reflects intrinsic feature superposition or simply the episodic binding of disparate narrative elements as an episode unfolds. However, we propose that these two interpretations are not mutually exclusive alternatives, but rather two sides of the same computational coin; representing rich, compositional, and relational stories with a finite population of neurons necessitates distributed, non-linear mixed selectivity. While presenting words in isolation has been suggested as a control to disentangle lexical tuning from contextual linking, such an approach alters the operational regime of the circuit by stripping away narrative structure.

The polysemantic neuronal tuning we observe recalls the dispersed sampling characteristic of grid cells (Moser et al., 2008). Grid cells are a canonical example of neurons whose firing fields tile an environment with multiple, widely separated fields. Crucially, the canonical hexagonal periodicity of grid cells breaks down in three-dimensional or topologically complex environments, resulting in fragmented, locally ordered firing fields (Ginosar et al., 2021, 2023; Grieves et al., 2021; Krupic et al., 2015; Mainali et al., 2025). This fact suggests that navigational coding neurons, at least in spaces of more than two dimensions, exhibit a form of spatial polysemanticity. The neurons in our dataset, which code locations in a high dimensional semantic embedding space, show a similar pattern: isotropic overdispersion without hexagonality. We hypothesize that these overdispersed response sets may reflect the same computational principle that may motivate 3-D grid coding, maximizing coverage of a representational space while minimizing redundancy and interference. In spatial navigation, distributed firing fields support efficient generalization across locations (Bellmund et al., 2018; Whittington et al., 2020). More speculatively, an overdispersed sampling of conceptual space could allow hippocampal neurons to link distant experiences while preserving separability among neighboring memories, thereby facilitating relational binding, and flexible inference, and, potentially, creativity.

Our results bear on longstanding debates concerning localist versus distributed coding schemes in the medial temporal lobe (Bowers, 2009; Plaut & McClelland, 2010; Haxby et al., 2001). These researchers, among others, argued about the relative merits of localist versus distributed codes, on both normative and empirical grounds. In tandem with the established concept cell literature, our results provide empirical support for the distributed side, without excluding a hybrid combined scheme. More broadly, our results underscore the value of adopting the population doctrine perspective (Ebitz & Hayden, 2021; Saxena & Cunningham, 2019; Urai et al., 2022; Vyas et al., 2020). Such an approach has proven fruitful in domains ranging from motor control to decision making (Cao et al., 2025; Churchland et al., 2012; Cunningham & Yu, 2014; Elsayed et al., 2016; Jazayeri & Ostojic, 2021; Tang et al., 2023; Yan et al., 2026). Specifically, conceptualizing language representations as geometric structures provides a natural bridge between neural data and computational models of semantics, which similarly represent meaning in continuous vector spaces (Franch et al., 2025; Kriegeskorte & Douglas, 2018; Piantadosi et al., 2024).

## METHODS

### Ethics approval

The experiment’s procedures were conducted in accordance with the standards set by the Declaration of Helsinki and approved by the Institutional Review Board for Baylor College of Medicine, Houston, TX (H-50885 and H-18112). Participants provided written informed consent after the experimental procedure had been fully explained and were reminded of their right to withdraw at any time during the study.

### Patient recruitment

Fifteen English-speaking epilepsy patients participated in the study and were aware that participation was voluntary and would not affect their clinical course. Patients’ age ranged from 18-48 years old (mean ± s.e.m = 30.07 ± 9.15 years), with 9 female patients and 6 male patients.

### Experimental stimuli

Participants listened to a set of naturalistic spoken narratives selected to be engaging and linguistically diverse (Franch et al., 2025). Specifically, the stimulus set comprised six episodes from The Moth Radio Hour (each approximately 5-13 minutes long), totaling 47 minutes and 25 seconds of audio (7346 words). The stories were: “Life Flight,” “The One Club,” “The Tiniest Bouquet,” “My Father’s Hands,” “Wild Women and Dancing Queens,” and “Juggling and Jesus.” Each excerpt featured a single speaker delivering an autobiographical story to a live audience. Audio was presented continuously through the built-in speakers of the participant’s hospital television. To synchronize stimulus timing with neural recordings, the audio output from the playback computer was routed as an analog input directly into the Neural Signal Processor (sampled at 30 kHz), ensuring precise temporal alignment between the stimulus waveform and recorded neural activity.

### Microelectrodes recording in the hippocampus

None of the analyzed data included seizures. Single neuron data were recorded from stereo-electroencephalography (sEEG) electrodes with microwires extending from the tip (AdTech Medical Behnke-Fried). Each patient had an average of 3 probes terminating in the left and right hippocampus. Electrode locations were verified by co-registered pre-operative MRI and post-operative CT scans. Each probe includes 8 microwires, each with 1 contact, specifically designed for recording single-neuron activity. Single neuron data were recorded using a 512-channel *Blackrock Microsystems Neuroport* system sampled at 30 kHz. To identify single neuron action potentials, the raw traces were spike sorted using the *Wave_clus* sorting algorithm (Chaure et al., 2018) and then manually evaluated. Noise was removed and each signal was classified as multi or single unit using several criteria: consistent spike waveforms, waveform shape (slope, amplitude, trough-to-peak), and exponentially decaying ISI histogram with no ISI shorter than the refractory period (1 ms).

The initial number of neurons was simply the total number of neurons that we collected. All preprocessing was done by a lab member blind to the goals of the study. In preprocessing (spike sorting), we only kept the isolated neurons. We analyzed all collected neurons and did not restrict analyses to task-responsive neurons.

### Firing rate responses to words

We computed firing rates during a time window starting 80 ms after the onset of each word (to account for the approximate response latency) and lasting the duration of the word plus 40 ms, accounting for transmission delays to hippocampus (Charest et al., 2009; Chavez et al., 2025; Zhu et al., 2026). Spike counts were divided by window duration and multiplied by 1000 to yield spikes per second.

### Control analysis: firing rate responses to words in narrower time windows (Figure S15)

To control for the possibility that early responses to the next word may cause spurious coding density and polysemanticity, we computed firing rates during a narrower time window (starting 80 ms after the onset of each word and lasting the duration of the word). Spike counts were divided by window duration and multiplied by 1000 to yield spikes per second.

### Audio transcription

After experiments, the audio.wav file was automatically transcribed using Assembly AI, a state-of-the-art AI model trained to transcribe speech. Speaker diarization was achieved with Assembly AI’s internal speaker diarization model. The transcribed words, corresponding timestamps, and speaker labels output from these models were converted to a TextGrid and then loaded into Praat (Boersma, 2001), a well-established software for speech analysis (https://www.fon.hum.uva.nl/praat/). The original *.wav* file was also loaded into Praat. Trained lab members manually inspected the spectrograms, timestamps, and speaker labels, correcting each word to ensure that onset and offset times were precise and assigned to the correct speaker (this process took about 4-5 hours per minute of speech). The TextGrid output of corrected words and timestamps from Praat was converted to an Excel file and loaded into Python (version 3.11) for further analysis.

### Electrode visualization for Figure 1A

Electrode visualization was performed using MNE-Python (version 0.24.0) with FreeSurfer’s fsaverage template brain. The visualization pipeline created an ultra-transparent glass brain (3% opacity) with highlighted hippocampal structures to clearly display electrode positions within deep brain regions. For each patient, DICOM images of the preoperative T1 anatomical MRI and the postoperative Stealth CT scans were acquired and converted to NIfTI format (Li et al., 2016). The CT was aligned to MRI space using FSL (Jenkinson & Smith, 2001; Jenkinson et al., 2002). The resulting co-registered CT was loaded into BioImage Suite (version 3.5β1; Joshi et al., 2011), and the electrode contacts were manually localized. Electrodes’ coordinates were converted to the patient’s native space using iELVis MATLAB functions (Groppe et al., 2017) and plotted on the Freesurfer (version 7.4.1) reconstructed brain surface (Dale et al., 1999). Microelectrode coordinates are taken from the first (deepest) macro contact on the Ad-Tech Behnke Fried depth electrodes. RAVE (Magnotti et al., 2020) was used to transform each patient’s brain and electrode coordinates into MNI152 average space. The coordinates were plotted together on a glass brain with the hippocampus segmentation.

### Semantic embedding extraction from language models

#### fastText embeddings

We used the pre-trained *fastText* model in MATLAB to extract word embeddings for all words in our dataset (Joulin et al., 2016; Mikolov et al., 2013). This pre-trained model provides 300-dimensional word embedding vectors, trained on millions of words text, to capture semantic relationships between words. Of note, these embeddings are static, that is, a given word gets the same embedding no matter the context in which it occurs.

#### GPT-2 embeddings

In an excel file of our transcribed words, each row corresponded to an individual token, including punctuation and sentence terminators such as “.”, “?”, and “!”. We fed each stimulus sentence by iterating through the file and appending words until encountering a recognized sentence delimiter. This process yielded multi-word segments that more accurately captured natural linguistic boundaries. Any remaining words following the last delimiter were grouped into a final sentence. Notably, punctuation tokens were preserved to maintain contextual fidelity but were tracked carefully to avoid introducing sub-word alignment errors during tokenization. To extract GPT-2 embeddings, we employed the “gpt2-large” model (Radford et al., 2019) from the Hugging Face Transformers library (Wolf et al., 2020). This version of GPT-2 features a total of 37 layers (1 initial embedding layer plus 36 transformer layers), each producing 1280-dimensional hidden states, with a maximum context length of 1024 tokens. Although GPT-2 supports a 1024-token context window, all embeddings used in this study were computed within individual sentences only - that is, the model’s context was reset whenever a sentence-ending delimiter (e.g., “.”, “!”, “?”) was encountered. We extracted hidden states from all 37 layers using a progressive context approach designed to incorporate GPT-2’s autoregressive nature. Specifically, for each sentence, we began with an empty context and added words sequentially. After adding each new word, we tokenized the updated text string via GPT-2’s Byte Pair Encoding (BPE) tokenizer, ran the token sequence through the model in no-grad (inference-only) mode, and retrieved the hidden states. We then identified the newly appended word’s sub-word tokens and averaged their corresponding vectors to obtain a single word-level embedding. Because GPT-2 often splits words into multiple sub-word fragments (otherwise known as “tokens,” e.g., “computa” + “tion”), averaging those fragments preserves consistency with the transcript’s original word boundaries. The GPT-2 outputs resulted in 1280-dimensional embeddings for each layer.

#### Maximal-context embedding extraction (control for Figure 4)

To test whether restricting the GPT-2 context to a single sentence affected the context-dependent results, we re-extracted word embeddings with the model’s full context window. The procedure was identical to the main extraction (GPT-2 large; all 37 hidden states, 1280 dimensions; progressive left-to-right causal context; word-level vectors obtained by averaging sub-word tokens aligned by character offset) except that the context was not reset at sentence boundaries. All analyses in Figure 4 that depend on embeddings (Figure 4A-C and 4G-K) were then repeated with these maximal-context embeddings and are reported in Figure S13; analyses that use only neural data (Figure 4D-E, Figure 5D) are unaffected by this choice.

#### GPT-2-XL and Gemma-2 embeddings (Figure S8)

To test whether our results depend on the specific language model, we repeated the embedding extraction with two additional autoregressive models: GPT-2-XL (Radford et al., 2019; 48 transformer layers plus the input embedding layer, yielding 49 hidden states of 1600 dimensions) and Gemma-2-9B (Gemma Team, 2024; 42 transformer layers plus the input embedding layer, yielding 43 hidden states of 3584 dimensions). Both were loaded from the Hugging Face Transformers library (Wolf et al., 2020) and run in inference-only mode: GPT-2-XL in 32-bit floating point and Gemma-2 in bfloat16 (its native training precision) with the eager attention implementation, which correctly applies Gemma-2’s attention logit soft-capping. Hidden states were upcast to 32-bit precision before averaging. Sentence segmentation was identical to the GPT-2-large pipeline: the transcript was divided into sentences at terminal punctuation (“.”, “?”, “!”), yielding 556 sentences from the 7346-word corpus, and the model’s context was reset at each sentence boundary. Because both models are causal decoders, each token’s hidden state depends only on preceding tokens; a single forward pass over the full sentence therefore produces hidden states mathematically identical to the word-by-word incremental procedure described above, and we used this computationally equivalent single-pass implementation. Sub-word tokens were assigned to words by character-offset alignment: each word received all tokens whose character span overlapped its span in the sentence string, and their hidden states were averaged to yield one word-level vector per layer. Special tokens with empty character spans (Gemma-2’s beginning-of-sequence token) were excluded from averaging.

### Ridge regression analysis for Figure 1D

We used ridge regression to learn a linear mapping from semantic embeddings to single-neuron firing rates. The predictor matrix consisted of the first 100 PCs (from PCA, explain 61.52% variance) of GPT-2 Layer-36 embeddings (Radford et al., 2019), augmented with an intercept column. A separate ridge regression model was fit for each neuron. The ridge penalty was applied to all predictor dimensions except the intercept term. We selected the regularization parameter λ via 5-fold cross-validation over words. For each candidate λ (25 values logarithmically spaced from 10^-3^ to 10^5^), the total cross-validated Gaussian log-likelihood was computed across all neurons and folds, then we selected the λ which maximizes this quantity. Specifically, for each fold, the model was trained on the training partition and evaluated on the held-out test partition. Encoding model quality was quantified as the improvement in cross-validated Gaussian log-likelihood relative to an intercept-only null model (ΔLL = LL_model_ - LL_null_). The null model predicted each neuron’s firing rate as the training-set mean, with its own variance estimated from training-set residuals. ΔLL was summed across neurons and test observations within each cross-validation fold and then summed across folds. The 5-fold partition was a random split of the 1221 noun words pooled across the six stories, with no temporal blocking by story. Random splits can inflate encoding performance when training and test samples are temporally autocorrelated (Hadidi, Feghhi et al., 2026); this concern is mitigated here because each sample is a single noun with its own embedding rather than a lagged feature stream, consecutive nouns were separated by a median of 4 words (1.1 s) and typically fell in different sentences, and Hadidi et al. report that word-level electrophysiology data of this kind are far less affected than continuous fMRI data.

### Control analyses for Figure 1D (results see Figure S1A-C)

To test whether the distributed scaling pattern reflects a genuine property of the neural code or an artifact of the encoding pipeline, we compared two encoding pipelines with opposing architectural biases. The original pipeline used PCA to reduce GPT-2 Layer-36 embeddings to 100 components, yielding a dense, signed basis, followed by ridge regression, which penalizes large weights and favors smooth, distributed solutions. The control pipeline replaced PCA with non-negative matrix factorization (NMF), which decomposes embeddings into non-negative, parts-based components that naturally permit sparse word loadings, and replaced ridge regression with elastic net (L1 ratio = 0.9, the estimator strongly favors sparse solutions), which permits exactly zero weights and can reveal sparse patterns if present in the data. Because NMF requires non-negative input, GPT-2 Layer 36 embeddings were shifted to non-negative values by subtracting the column-wise minimum from each feature dimension prior to decomposition. The shifted matrix was then factorized into 100 components using the multiplicative update algorithm (Lee & Seung, 1999) with 10 random initializations, retaining the factorization with lowest reconstruction error. The resulting word-by-component score matrix (W) was z-scored across words to place components on comparable scales for regression. Elastic net regularization paths were fit per neuron with 5-fold cross-validation for lambda selection. Both pipelines were evaluated using the same cross-validated ΔLL metric, and the analysis was repeated for three embedding models (fastText, GPT-2 Layer-24, and GPT-2 Layer-36) to ensure generality.

### Encoding-based decoding (Figure 1E-F)

To evaluate whether the word could be recovered from population activity, we implemented an encoding-based decoding procedure (Tang et al., 2023). Both neural firing rates and GPT-2 Layer-36 embeddings were z-scored across words. The embedding matrix was further reduced to 100 principal components via PCA (Yan et al., 2026; Franch et al., 2026). The first 100 PCs explain 88.00%, 58.97%, and 61.52% of the variance in fastText, GPT-2 Layer-24, and GPT-2 Layer-36 embeddings, respectively; the first 50 PCs explain 72.17%, 43.15%, and 46.57%; and the first 5 PCs explain 24.12%, 11.33%, and 13.66%. Repeating the encoding analysis with 50 PCs did not change any result, and GPT-2 Layer 36 remained the best-performing embedding, indicating that the first 100 PCs capture the embedding structure well.

#### Per-neuron lambda tuning

A separate ridge regression encoding model was fit for each neuron. The regularization parameter λ was tuned independently per neuron using 5-fold cross-validation over words, selecting the λ (from 25 values logarithmically spaced between 10^-3^ to 10^5^) that minimized mean squared prediction error.

#### Leave-one-out decoding

Decoding was performed using leave-one-out cross-validation (LOOCV) across words. On each iteration, one word was held out as the test item and the encoding model was trained on the remaining words using the per-neuron λ values determined above. The trained model was then used to generate predicted neural population vectors for all candidate words from their embeddings. The observed neural population vector for the held-out word was compared to each candidate’s predicted vector using Pearson correlation, and candidates were ranked by similarity. The decoding rank indicates the position of the true word among all candidates (rank 1 = the true word has the highest similarity score; rank 2 = the true word has the second highest similarity score; rank = (W+1)/2, where W is the number of candidate words, represents the chance level).

#### Statistical testing

Decoding significance was assessed using a permutation test (500 permutations). On each permutation, the pairing between embeddings and neural responses was shuffled within the training set only, preserving the true test observation. The full LOOCV decoding procedure was re-run under this null, and top-1 accuracy was recorded. The p-value was computed as the proportion of permutation accuracies equal to or exceeding the observed accuracy.

#### Construction of Representational Dissimilarity Matrices (RDM) and representation similarity analyses (RSA) (*Figure 1G*)

For both neural and semantic spaces, we computed *Representational Dissimilarity Matrices* (RDMs; Diedrichsen & Kriegeskorte, 2017; Nikolaus Kriegeskorte & Kievit, 2013; Nili et al., 2014) using cosine distance as the dissimilarity metric. The cosine distance between two vectors **x** and **y** is defined as:

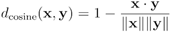

For a dataset with *n* words, this yields a condensed distance vector of length n*(n-1)/2 containing all unique pairwise distances. The neural RDM for language *l* was computed as:

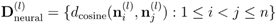

Where 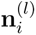 represents the neural population vector for word *i* in language *l*. Similarly, the semantic RDM was computed as:

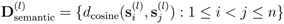

Where 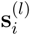 represents the GPT2-Layer-36 embedding vector for word *i* in language *l*.

### Neural-semantic correspondence quantification

The correspondence between neural and semantic representational geometries was quantified by computing Spearman correlation between the vectorized RDMs (i.e., 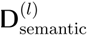 and 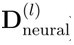). This correlation coefficient quantifies the degree to which neural dissimilarity patterns mirror semantic dissimilarity patterns. A high positive correlation indicates that words with similar meanings (small semantic distance) tend to evoke similar neural responses (small neural distance).

### Representation similarity analyses and decimation scaling analyses for Figure 1H-I

To assess how the correspondence between neural and LLM semantic geometry scales with population size, we performed a decimation analysis on the RSA. For each population size N ∈ {5, 10, 20, 40, 80, 160, 320, 498}, we randomly sampled N neurons from the full population and computed the Spearman rank correlation (rho) between pairwise cosine distance vectors in neural space and in GPT-2 Layer-36 embedding space (we also did analyses in GPT-2 Layer 24 and fastText embeddings). This procedure was repeated 200 times at each population size, each time with a different random draw of neurons. A shuffle control was performed by randomly permuting the word order of the neural firing rate matrix before computing pairwise distances, while the GPT embedding distances were held fixed (50 repetitions per population size). For visualization in Figure 1H, the plotted quantity is ρ above chance (real mean minus shuffle mean at each N). Shaded bands denote 95% confidence intervals (2.5th-97.5th percentiles) of the neuron-resampling distribution, shifted by the shuffle mean. That is, the lower and upper bounds correspond to the 2.5th and 97.5th percentiles of the real ρ distribution minus the shuffle mean at each N, treating the shuffle mean as a fixed baseline correction rather than propagating shuffle variance through error combination. At N = 498, the band collapses to zero width because all neurons are included in every iteration, leaving no variability across resampled subsets. Figure 1I reports the effect size (z-score) at each population size, computed as the mean ρ above chance divided by the standard error of the difference. Because z-scores are scalar values at each population size rather than distributions, they are plotted as discrete points with horizontal reference lines indicating significance thresholds (p < 0.05 and p < 0.001).

### Control analyses for Figure 1H-I (results see Figure S1D)

To rule out the possibility that scaling improvements reflect mere redundancy (i.e., averaging out noise across copies of the same signal), we implemented a copy-plus-noise control. We identified the single neuron with the highest cross-validated encoding performance (ΔLL relative to an intercept-only null), fit a ridge regression to decompose its activity into a predicted (signal) component and a residual (noise) component, then generated a synthetic population of up to 498 units by adding independent Gaussian noise (matched in variance to the empirical residuals) to copies of the signal. We then ran the identical RSA decimation procedure on this synthetic population. If the real data exhibits a qualitatively different scaling profile from the copy-plus-noise control, particularly continued gains at large N where redundant copies would plateau, this constitutes evidence that additional neurons carry non-redundant semantic information, consistent with a distributed code.

### Multidimensional Scaling (MDS) network visualization (Figure 1J-M, Figure 3H, Figure 4D and Figure 5C)

In Figure 1J-M, Figure 3H, Figure 4D and Figure 5C, we employed classical Multidimensional Scaling (MDS) (Mead, 1992) combined with network visualization techniques to visualize the geometric organization of neural representations for words. This approach reveals the topological structure of semantic representations in neural space by projecting high-dimensional neural patterns onto interpretable low-dimensional manifolds while preserving pairwise dissimilarities. The resulting 2D coordinates were visualized as network graphs where nodes represent words and edges encode distance between words.

### Symmetric non-negative matrix factorization (NMF) of neural semantic tuning (Figure 1N-O)

To identify shared semantic modes across the neural population, we decomposed the structure of neuron-level semantic tuning using symmetric NMF.

#### Tuning vectors

For each neuron, a tuning vector was estimated by fitting a ridge regression model (method see above).

#### Neuron similarity matrix

Tuning vectors were rectified (negative values set to zero) and L2-normalized. The neuron-by-neuron similarity matrix S was computed as the dot product of the normalized tuning vectors, yielding a non-negative symmetric matrix with diagonal entries set to 1.

#### Symmetric NMF

The similarity matrix was decomposed as S ≈ WWᵀ, where W (N × K) is a non-negative weight matrix and K is the number of semantic modes. The factorization was performed using multiplicative update rules: at each iteration, W was updated element-wise as W ← W ⊙ (SW) / (W(WᵀW) + ε), followed by column-sum normalization. The algorithm was run for 500 iterations per replicate, with 50 random initializations, and the solution with the lowest relative Frobenius reconstruction error was retained.

#### Selection of K

The optimal number of modes was selected using an elbow method. Reconstruction error was computed for K = 2 to 15, with 10 random restarts at each K. The second derivative of the reconstruction error curve was computed, and the K at which the second derivative was maximized (i.e., the point of greatest curvature change) was selected as optimal (K = 13).

#### Validation

We confirmed that each NMF solution captured meaningful structure by comparing its reconstruction error against a null distribution. For each of 500 permutations, we generated a random weight matrix of the same dimensions with entries drawn from a uniform distribution, column-normalized it, and scaled it to match the column sums of the real W. We computed the p-value as the proportion of random reconstruction errors less than or equal to the real reconstruction error.

#### Mode characterization

For each mode k, we computed a prototype vector in semantic embedding space as the weighted average of all neurons’ tuning vectors (prior to rectification), with weights given by the k-th column of W, normalized to sum to 1. We then L2-normalized each prototype and projected the embedding matrix onto each prototype to obtain word scores, which quantify how strongly each word aligns with the mode’s semantic direction. We reported the top 8 words per mode (in Figure 1O) after labeling repeated words with numeric suffixes (e.g., “golf ^1^”, “golf ^2^” in mode#1) to distinguish multiple occurrences.

#### Control analysis: sign-split tuning vectors

Because rectification discards negative weights, we repeated the full NMF pipeline using a sign-split representation of each tuning vector, β → [β⁺; β⁻], where β⁺ = max(β, 0) and β⁻ = max(-β, 0). The concatenated vectors (2 × 100 dimensions) were L2-normalized and the similarity matrix was computed as their dot product, which is non-negative by construction and preserves all weight information. K selection, validation, mode characterization and the semantic-distance test were then applied unchanged. Across the 15 patients this yielded 151 modules, of which 6 were significantly clustered, 59 were at chance and 86 were significantly dispersed (dispersed vs. clustered, binomial p < 0.001), reproducing the pattern reported in Table S1.

### Definitions of word categories

To identify the natural semantic categories present in our word data (Figure 1O), all unique words were clustered into groups based on related meanings using a word embedding approach (Joulin et al., 2016; Mikolov et al., 2013). We used the pre-trained *fastText Word2Vec* model in MATLAB to extract word embeddings for all words in our dataset (Joulin et al., 2016; Mikolov et al., 2013). This pre-trained model provides 300-dimensional word embedding vectors, trained on millions of words text, to capture semantic relationships between words. To compute and visualize semantic clusters, we used a t-distributed Stochastic Neighbor Embedding (t-SNE) algorithm (Maaten & Hinton, 2008) on word embedding values to reduce the dimensionality of each unique word based on their cosine distance to all other words, thus reflecting their semantic similarity. Words with similar meanings now have similar 2D coordinates. We then applied the K-means clustering algorithm to these 2D word representations and visualized clustered words on a 2D word map (k = 12 clusters, determined from the silhouette score; Rousseeuw, 1987). We then manually inspected and assigned a distinct semantic label to each cluster and adjusted categories for accuracy. For example, words bordering the edges of categories might get mis-grouped and were manually corrected. This method is comparable to that used in other studies of semantics (Huth et al., 2016; Jamali et al., 2024).

### Simulation of concept cells (sparse semantic coding) under the Waydo et al. theoretical model (Figure S3)

To enable a direct comparison between classic concept-cell results from image presentation paradigms and neural responses during continuous naturalistic language listening, we developed a simulation framework that generates concept-cell-like sparse coding under controlled target sparseness levels. The simulation follows the logic of the Waydo-style sparse coding model and published concept cell papers (Quiroga et al., 2008, 2007, 2005; Waydo et al., 2006): only a small fraction of neuron-stimulus pairs evoke detectable responses, and “responsive neurons” and “evocative stimuli” emerge from that sparse response matrix. The model captures sparse stimulus selectivity but does not explicitly simulate invariance across different exemplars of the same concept, a defining property examined in classic concept-cell experiments. The simulation was calibrated to reflect both (i) published concept-cell benchmarks and (ii) the timing constraints of naturalistic listening.

We simulated a population of N = 498 neurons and W = 516 stimuli (unique nouns), matching the scale of our empirical dataset used for comparison. The stimulus set included both common and proper nouns (e.g., “Applebee’s,” “Texas,” and “Montana”), without assigning different simulation rules to these categories. Each unique noun was treated as a distinct stimulus identity. Our naturalistic listening paradigm did not provide controlled exemplars for testing response invariance. Although some nouns appeared repeatedly, their occurrences were embedded in different linguistic contexts (see Figure 4). To preserve the temporal structure of the language stimulus set, each simulated stimulus was assigned a real word duration taken from our noun stimulus list. Word durations were converted to seconds and used as the stimulus presentation duration for the simulated “response window.” In addition, each trial included a fixed baseline window (0.1 s) to mimic the baseline period used in image-based concept-cell paradigms.

For each simulated sparseness condition, we specified a target probability that a neuron responds to a stimulus (the target “sparseness” level). For each neuron-stimulus pair, a latent binary variable determined whether the response window was drawn from a baseline firing regime or an elevated firing regime. Baseline and response firing rates were specified by parameters r_0_ (baseline rate) and r_1_ (response rate). We set the r_0_ to 0.02Hz and r_1_=5Hz, based on empirical results (Quiroga et al., 2005). Crucially, because stimulus durations vary across words, we generated expected spike counts by combining the firing rate regime with the word-specific duration.

To capture realistic trial-to-trial variability beyond Poisson spiking, spike counts were generated using a Gamma-Poisson (negative binomial) model governed by a Fano factor parameter (F = 2.66; Quiroga et al., 2005). This creates overdispersed spike counts consistent with observed neural variability. A lower bound on the Gamma shape parameter was imposed (kappa_floor_ = 10) to prevent unstable behavior for very small expected counts.

### Waydo-style detection criterion and thresholding

To classify a neuron-stimulus pair as a “response,” we implemented a detection rule analogous to concept-cell screening criteria: (1) For each neuron, baseline spike counts were used to compute a neuron-specific baseline mean and standard deviation. (2) A neuron-specific detection threshold was set as mean + kSD × SD, with kSD = 5, matching the commonly used stringent criterion in concept-cell studies. (3) A response was counted only if the response-window spike count exceeded the threshold and also exceeded a minimum spike-count criterion (nSpkMin = 2). This produced a binary response matrix of size W × N, indicating whether each neuron responded to each stimulus.

### Calibration of latent response probability

For each target sparseness level, the simulation calibrated the latent response probability (pi) using a bisection search so that the empirically observed response probability (the fraction of ones in the response matrix) matched the target sparseness (α) as closely as possible. Following Waydo et al. (2006), sparseness α is defined as the probability that a neuron responds to a randomly selected stimulus. The proportion of responsive neurons is therefore not independently specified but emerges from α and the number of stimuli W, with an expected fraction of 1 - (1 - α)^W^ under independent stimulus-response assignments. For example, α = 0.2% predicts approximately 16% responsive neurons for 88 stimuli (cf. Quiroga et al., 2005). This ensured that different sparseness conditions could be compared on a common scale. We simulated 10 target sparseness levels spanning 0.2% to 1.0% (inclusive). For each sparseness condition, we computed the standard Waydo-style population statistics from the binary response matrix: (1) Number of responsive neurons (Nr): neurons that respond to at least one stimulus; (2) Stimuli per responsive neuron: average number of stimuli eliciting a response among responsive neurons; (3) Neurons per evocative stimulus: average number of neurons activated among evocative stimuli. We additionally computed the fraction Nr/N as a “responsive neuron fraction.” These quantities were compared to published benchmarks (**Figure S3E**).

### Validation of the Waydo-style lifetime sparseness model

We repeat the simulation process above, but using the exact number of neurons and stimulus numbers to make sure the simulation results can be compared with the published results. Specifically, we set the time for baseline and response as 0.7s (Quiroga et al., 2008: “*The response to a picture at a specific duration was defined as the median number of spikes across trials between 300 and 1000 ms after stimulus onset.*” Also see Quiroga et al., 2007, 2005). For Quiroga et al., 2008, neuron number=440, Stimulus=98. For Quiroga et al., 2007, number=1547, Stimulus=88. For Quiroga et al., 2005, neuron number=993, Stimulus=88.

### Population sparseness (Figure 2A-B)

To quantify the fraction of the neural population activated by each word, we computed population sparseness (Rolls & Treves, 1997) for each word. Whereas lifetime sparseness characterizes the selectivity of a single neuron across stimuli, population sparseness characterizes the selectivity of a single stimulus across the population. For a given word with firing rates r = [r_1_, r_2_, …, r_N_] across N neurons:

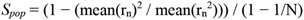

Values of *S_pop_* range from 0 (all neurons respond equally, maximally dense) to 1 (only a single neuron responds, maximally sparse). Words for which fewer than two neurons had valid (non-zero) firing rates, or for which all firing rates were zero, were excluded from the population sparseness computation. We compared the observed population sparseness distribution to simulated populations across 10 levels of target sparseness (alpha).

### Lifetime sparseness (Figure 2C-D)

To quantify the selectivity of individual neurons, we computed lifetime sparseness (Rolls & Treves, 1997) for each neuron. For a neuron with firing rates r = [r_1_, r_2_, …, r_W_] across W words, lifetime sparseness was defined as:

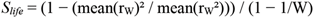

where mean(r_W_) is the mean firing rate and mean(r_W_^2^) is the mean squared firing rate across all words.

Values of *S_life_* range from 0 (uniform response across all words, maximally dense) to 1 (response to only a single word, maximally sparse). Neurons with zero total firing or fewer than two valid observations were excluded. To contextualize the observed sparseness distribution, we compared real neurons against simulated populations with known sparse coding properties. Simulated populations were generated across 10 levels of target sparseness (parameterized by alpha), spanning a range from highly sparse to moderately dense coding. To avoid inflating selectivity estimates through stimulus repetition, all sparseness metrics were computed on unique words only; repeated words of the same word were averaged prior to analysis.

### Normalized entropy (Figure 2E)

To quantify the uniformity of each neuron’s response distribution across words, we computed normalized entropy (Lehky et al., 2005). For each neuron with firing rates r = [r_1_, r_2_, …, r_W_] across W words, firing rates were converted to a probability distribution by normalizing to unit sum:

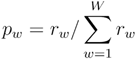

Shannon entropy was then computed as:

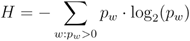

where the sum is restricted to words with *p_w_* > 0. Entropy was normalized by the maximum possible entropy for W words:

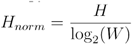

yielding values between 0 (neuron responds to only one word, maximally selective) and 1 (neuron responds uniformly to all words, maximally distributed). Neurons with zero total firing or fewer than two valid observations were excluded.

The distribution of normalized entropy for real neurons was compared against simulated populations across 10 levels of simulated sparseness, computed identically. Radar plots were additionally used to visualize the shape of the entropy distribution: for each condition, the histogram of normalized entropy values (20 bins spanning [0, 1]) was plotted in polar coordinates, with each bin mapped to an angular position and the bin probability mapped to radius. A uniform reference circle indicates the expected shape under a distributed code; deviations from circularity indicate sparse code. Computations were restricted to unique words, as described above.

### Information per word (Figure 2F)

To quantify how much information each neuron carries about word, we computed information per word (bits/word) using the framework (Skaggs, McNaughton, et al., 1992) adapted to the lexical domain. For each neuron with firing rates r = [r_1_, r_2_, …, r_W_] across W words, each with prior probability p(w) and duration T(w):

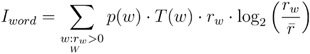

where 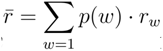 is the overall mean firing rate computed across all words. Because each unique noun was represented once, p(w) = 1/W for all words. T(w) is the duration of each word in the continuous speech stimulus, which varies across words. The information sum is restricted to words with non-zero firing rates (*r_w_* > 0), as the log_2_ term is undefined at zero. Intuitively, this metric accumulates information from every word for which a neuron’s firing rate deviates from its mean: even neurons with high entropy (broadly distributed firing) carry substantial information if their rate modulations, though small, are systematic across words.

### Decimation RSA for simulated concept cells (Figure S4)

To test whether the monotonic scaling of representational similarity with population size (Figure 1H) could arise under a sparse concept-cell code, we repeated the RSA decimation analysis on simulated concept cell populations. For each of the 10 sparseness levels (response probability ranging from 0.20% to 1.00% of words; see Figure S3 and *method* <u>Simulation of concept cells (sparse semantic coding) under</u> <u>the Waydo et al. theoretical model</u> above), we constructed a population of 498 simulated neurons and computed pairwise cosine distances between all words in neural firing rate space. We then correlated the resulting neural distance matrix with the GPT-2 Layer 36 embedding distance matrix using Spearman rank correlation, following the same procedure as the real-neuron analysis. At each population size (N = 5, 10, 20, 40, 80, 160, 320, 498), we randomly subsampled N neurons and computed the RSA correlation, repeating this procedure 200 times to obtain means and 95% confidence intervals. Unlike the real hippocampal population, which shows a monotonic increase in RSA correlation with population size, simulated concept cell populations produced near-zero correlations at all population sizes and sparseness levels (Spearman rho range across all simulated sparseness: -0.014 to 0.030 at N = 498), confirming that a sparse code in which each neuron responds to very few words cannot reconstruct the continuous semantic geometry captured by the language model.

### Single-neuron tuning landscapes in semantic space (Figure 3A-B)

To visualize the tuning profile of individual neurons across semantic space, we extracted GPT-2 embeddings (Layer 36) for 50,000 nouns from an independent large text corpus, z-scored them, and projected them into a two-dimensional semantic map using t-SNE (MATLAB’s tsne function, Barnes-Hut algorithm, perplexity = 30).

#### Predicted firing rates

For each neuron, we used the ridge regression encoding model fit on the experimental data (see Methods for Figure 1D) to predict firing rates across the 50,000 nouns. The z-scored GPT-2 embeddings were projected onto the first 100 principal components (computed from the experimental embedding matrix), and predicted firing rates were computed as FR = X · β + β₀, where X is the 50,000 × 100 PC score matrix, β are the regression weights, and β₀ is the intercept. Predicted rates were passed through a softplus nonlinearity (f(x) = log(1 + eˣ), implemented in numerically stable form) to enforce non-negativity, then z-scored across nouns.

#### Visualization

For each neuron, the predicted firing rate (z-scored) was plotted as a color-coded scatter on the t-SNE coordinates. To reduce file size, only nouns with raw predicted firing rate > 0 were plotted. Points were colored using a rectified sequential colormap (white for values ≤ 0, inferno gradient for positive values, saturating at a z-score of 2.5). Marginal distributions along each t-SNE axis were computed by binning coordinates into 100 equally spaced bins and computing the mean absolute z-scored firing rate within each bin. These marginals were displayed as gray bar plots along the top (t-SNE dimension 1) and right (t-SNE dimension 2) edges of the scatter plot, providing a summary of the distribution of the neuron’s tuning across semantic space. Axis limits were held constant across neurons using the full range of the 50,000-noun t-SNE coordinates.

### Grid-cell-like feature analysis of semantic modes

To assess whether neurons with multiple semantic modes exhibit spatial organization analogous to grid cells, we quantified four canonical grid-cell properties: regularity, equidistance, isotropy, and hexagonal symmetry. Statistical significance was established via permutation testing (500 permutations for regularity, equidistance, isotropy, and gridness score). Details of each analysis are described below.

#### mode identification and centroid computation

For each neuron, predicted firing rates across 50,000 nouns were softplus-transformed and z-scored. Words exceeding the 98th percentile of this distribution were designated as “high-firing words.” These high-firing words were then clustered using DBSCAN (ε = 5, minPts = 5) applied to their coordinates in the t-SNE semantic plane, where each resulting cluster defines one *semantic mode*, which is a spatially contiguous hotspot of elevated firing in semantic space. For each mode, we computed a centroid as the firing-rate-weighted average coordinate of its constituent words, such that words eliciting stronger responses exert greater influence on the centroid’s position. For example, if a mode contains the words “dog,” “cat,” “lion,” “fish,” and “bird,” and the neuron fires most strongly to “dog,” the centroid will be pulled toward the position of “dog” in the embedding space rather than lying at the unweighted geometric center of all five words. Centroids were computed in both the two-dimensional t-SNE plane (for visualization) and the original high-dimensional embedding space (for regularity, equidistance, and isotropy analysis). Neurons with fewer than two modes were excluded from regularity and equidistance analyses; neurons with fewer than three modes were excluded from isotropy analyses.

#### Rationale for CV-based metrics in high-dimensional embedding space

Classical grid cell analyses quantify regularity, equidistance, and isotropy using two-dimensional spatial autocorrelation maps (Hafting et al., 2005; Stensola et al., 2012). These methods are inherently tied to 2D geometry and may not be directly applied to semantic embeddings, which reside in a 1280-dimensional space. We therefore adopted the coefficient of variation (CV) of pairwise distances as our primary metric, for several reasons. First, pairwise distances (here, cosine distances) are well-defined in arbitrary dimensions and do not require dimensionality reduction, avoiding distortions introduced by nonlinear projections such as t-SNE. Second, CV is a scale-free measure of dispersion that directly captures the intuition behind regularity: if all inter-mode distances are similar, CV is low regardless of the absolute scale or dimensionality. Third, a critical concern in high-dimensional spaces is distance concentration, whereby pairwise distances between random points converge to similar values as dimensionality increases, potentially inflating apparent regularity. Our permutation null model controls for this by shuffling word assignments among the same high-firing words while preserving mode sizes, so that any distance concentration effect applies equally to real and null distributions. Significance is therefore assessed relative to the regularity expected from random partitions within the same high-dimensional geometry, not relative to an absolute standard.

### Regularity (Figure 3C)

For each neuron with ≥ 2 modes, we computed pairwise cosine distances in original embedding space, then measured the coefficient of variation (CV = σ/μ) of these distances. Lower CV indicates more regular inter-mode spacing. To construct a null distribution, we randomly partitioned the set of non-noise high-firing words into groups matching the original mode sizes, recomputed centroids from shuffled assignments, and recalculated the pairwise distance CV. This was repeated 500 times per neuron. The z-score was defined as z = (CV_real_ - μ_null_) / σ_null_, where negative values indicate greater regularity than expected by chance. The permutation p-value was computed as the fraction of null CV values less than or equal to the observed CV.

### Equidistance (Figure 3D)

Using the same mode centroids and null model as above, we assessed whether each mode’s nearest neighbor (k = 1) was at a consistent distance across modes. We extracted the nearest-neighbor distance for every mode and computed the CV across modes. The same random-partition permutation procedure provided null distributions. Negative z-scores indicate that nearest-neighbor distances are more uniform than expected by chance. The permutation p-value was computed as the fraction of null CV values less than or equal to the observed CV.

### Isotropy in embedding space (*Figure 3E*)

For each neuron with ≥ 3 modes, we tested whether mode centroids were distributed uniformly in direction around the grand centroid in the original embedding space. We computed the grand centroid as the arithmetic mean of all mode centroids in embedding space, then obtained direction vectors from the grand centroid to each mode centroid, normalized to unit length. We computed all pairwise cosine similarities among these unit direction vectors and measured the CV of the resulting similarity values. For centroids distributed uniformly on a high-dimensional hypersphere, pairwise cosine similarities concentrate near zero with low variance; thus, lower CV indicates more isotropic arrangement. To establish a null distribution, we generated 500 sets of random unit vectors on the hypersphere (by drawing from an isotropic Gaussian and normalizing), matched in number to the observed modes, and computed the pairwise cosine similarity CV for each. The z-score was defined as z = (CV_real_ − μ_null_) / σ_null_, and the p-value as the fraction of null CV values less than or equal to the observed CV.

### Hexagonal symmetry (*Figure 3F*)

To test whether the spatial organization of semantic modes exhibits six-fold rotational symmetry analogous to entorhinal grid cells, we computed a gridness score for each neuron following established procedures from the spatial navigation literature, adapted for the semantic domain (Hafting et al., 2005; Langston et al., 2010). For each neuron, we constructed a two-dimensional firing rate map by binning all 50,000 nouns onto the t-SNE semantic plane using a uniform grid (bin width = 1 t-SNE coordinate unit) and averaging the z-scored predicted firing rates (after softplus transformation) within each occupied bin. Rate maps were smoothed with a Gaussian kernel (σ = 3 bins). We then computed the two-dimensional spatial autocorrelation of the smoothed rate map. To avoid edge effects and unvisited locations, bins with zero occupancy were excluded prior to smoothing, and the rate map was mean-centered using only occupied bins before computing the autocorrelation.

The central peak of the autocorrelation (arising from the trivial zero-lag self-correlation) was excluded using a data-driven procedure following (Langston et al., 2010). We computed the radial autocorrelation profile by averaging autocorrelation values within concentric annular rings (width = 1 bin) at increasing distances from the center. The central exclusion radius was defined as the first local minimum or the first zero crossing (positive-to-negative transition) in this radial profile, whichever occurred at a shorter distance. This approach avoids arbitrary parameter choices and adapts to the autocorrelation structure of each individual neuron.

Outside the central exclusion zone, we identified the six nearest peaks (local maxima with positive autocorrelation values) sorted by distance from the autocorrelation center. A minimum of three detectable peaks was required for a neuron to be included in the analysis. The annular region for rotational symmetry analysis was defined with inner radius equal to the distance of the nearest peak and outer radius equal to the distance of the farthest peak among the six selected, with the central exclusion zone removed. This data-driven annulus definition ensures that the analysis targets the spatial frequency band containing the first-order periodic structure, without relying on arbitrary scaling factors.

To quantify hexagonal symmetry, we extracted autocorrelation values within the annulus and computed the Pearson correlation between these values and those obtained after rotating the autocorrelation map by 30°, 60°, 90°, 120°, and 150° (using bilinear interpolation). For a pattern with perfect six-fold symmetry, rotations by 60° and 120° map the pattern onto itself (yielding high correlations), whereas rotations by 30°, 90°, and 150° produce maximal mismatch (yielding low correlations). The gridness score was defined as the mean correlation at hexagonal-consistent angles (60°, 120°) minus the mean correlation at hexagonal-inconsistent angles (30°, 90°, 150°), such that positive values indicate hexagonal periodicity.

To assess statistical significance, we generated a null distribution for each neuron by randomly permuting predicted firing rates across the 50,000 nouns (500 iterations), recomputing the full rate map, autocorrelation, and gridness score for each permutation while holding the annulus geometry fixed to that of the real data. This word-identity shuffle is analogous to the spike-train time-shift procedure used in rodent grid cell studies (Langston et al., 2010), destroying any relationship between firing rate and position in semantic space while preserving the marginal distributions of both. The z-score was defined as z = (gridness_real_ − μ_null_) / σ_null_, and the permutation p-value as the fraction of null gridness scores greater than or equal to the observed value.

### Robustness to t-SNE and DBSCAN parameters (Figure 3)

To confirm that the geometric properties of semantic modes do not depend on the dimensionality-reduction or clustering parameters, we repeated the full pipeline (t-SNE projection, DBSCAN mode identification, and permutation testing for regularity, equidistance, isotropy, and gridness) across a grid of t-SNE perplexity values (30, 50, 100, 150) and DBSCAN minPts values (5, 10, 15, 20, 25), yielding 20 parameter combinations; ε was held at 5 throughout. All other steps, including the 98th-percentile threshold for high-firing words, centroid computation in the original embedding space, and the 500-permutation null models, were identical to those described above. All four effects remained statistically significant (p < 0.001) in every parameter combination.

### Validation of single-neuron overdispersion (Table S3)

To validate that polysemantic tuning is a property of directly observed neuronal responses rather than of the encoding model, we performed a model-free split-half analysis of measured firing rates. For each unique noun occurring at least twice in the stimulus (213 words), occurrences were randomly partitioned into two halves (A and B), and each neuron’s mean firing rate per word was computed separately within each half, yielding two statistically independent estimates of the neuron’s semantic tuning. Because most words occurred only a small number of times, word-level split-half estimates are dominated by sparse-spiking noise; we therefore aggregated words into 60 semantic clusters by applying k-means clustering (cosine distance, 5 replicates) to the word embeddings (GPT-2, layer 36), and averaged the word-level firing rates within clusters to obtain, for each neuron, a 60-dimensional cluster-level tuning profile in each half. Cluster assignments were derived from the embeddings alone and were therefore identical for, and independent of, both halves. The entire procedure was repeated over 100 random occurrence partitions, and all statistics were averaged across partitions within each neuron before population-level testing.

Tuning reliability was quantified as the Spearman correlation between each neuron’s half-A and half-B cluster profiles, compared against a null obtained by randomly permuting the half-B profile; neurons with fewer than 20 valid clusters or zero variance in either half were excluded. To test whether the top-driving semantic regions replicate in held-out data, we ranked clusters by half-A firing, selected the top K (K = 5, 7, 10 evaluated within identical partitions), and computed the mean percentile rank of these clusters in the independent half-B data (chance = 50; one-sample t-test versus 50 across neurons). To test whether the replicated preferred regions were polysemantic, we retained only the top-K clusters ranking above the 50th percentile in half B and computed the mean pairwise cosine distance between their centroids; this observed dispersion was compared, by paired t-test across neurons, against a size-matched null given by the expected dispersion of randomly drawn cluster sets (500 random draws per set size and partition). Neurons contributing fewer than three replicated clusters at a given K were excluded from the dispersion analysis at that K. Under a monosemantic organization, replicated preferred regions should be closer together in embedding space than randomly drawn regions; dispersion exceeding the random null therefore indicates that reliable tuning spans multiple semantically distant regions. No encoding model, regression, or extrapolation beyond the heard vocabulary was used at any stage of this analysis. All analyses used the same noun set as the main encoding analyses; random-number seeds were fixed for reproducibility.

### Semantic dispersion of preferred stimuli (Figure 3G-K)

To assess whether each neuron’s most-preferred words are semantically clustered (as expected under a concept-cell model) or dispersed across semantic space (as expected under a distributed code), we analyzed the pairwise semantic distances among each neuron’s top-ranked words. For each neuron, words with non-zero firing rates were ranked in descending order. Zero-firing-rate words were excluded because they carry no information about stimulus preference and their ties at zero make rank ordering arbitrary. The top N words (N ∈ {10, 20, 30, …, 100}) were identified, and the mean pairwise cosine distance in GPT-2-Layer-36 embedding space was computed among all unique pairs within this set. This observed distance was compared against a null distribution constructed by randomly sampling N words from the remaining words (i.e., words outside the top N) and computing the same mean pairwise distance. The null procedure was repeated 500 times per neuron. A neuron was included in the analysis for a given N only if it responded to at least 2N words with valid (non-NaN), non-zero firing rates, ensuring that both the top-N set and the null sampling pool (of at least N words) could be constructed. Because this inclusion criterion scales with N, the number of valid neurons, and hence the degrees of freedom of the population-level tests, decreases as N increases: neurons with sparse responses that qualify at small N (e.g., N = 10, requiring ≥20 responsive words) may fail to qualify at larger N (e.g., N = 100, requiring ≥200 responsive words). Then for each valid neuron, a z-score was computed as the difference between the observed mean distance and the null mean, divided by the null standard deviation. Two one-sided p-values were computed: p_cluster (proportion of null distances ≤ observed, testing for semantic clustering) and p_disperse (proportion of null distances ≥ observed, testing for semantic dispersion). This analysis was repeated across all values of N to assess the robustness of the effect across different thresholds of stimulus preference. We further used these two p-values to classify which neurons showed significant semantic clustering or semantic dispersion (p<0.05), among valid neurons at each N.

### Control analyses for top-N words dispersion (Figure S9)

To address the concern that semantic overdispersion among each neuron’s preferred words might be driven by noise in the tail of the firing rate distribution rather than by the neuron’s strongest responses, we performed a firing-rate-weighted control analysis. For each neuron, we identified the top N words by firing rate (N = 10 to 100) and computed a weighted mean pairwise cosine distance in GPT-2 Layer-36 embedding space, where each word pair was weighted by the product of their firing rates:

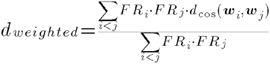

where *FR_i_* is the firing rate of the neuron to word *i* and *d*_cos_(***w****i,**w**j*) is the cosine distance between the GPT-2 Layer-36 embedding vectors of words *i* and *j*. This weighting ensures that the highest-firing words dominate the distance metric, effectively downweighting low-firing words near the selection threshold that might reflect noise rather than genuine tuning.

### Control analyses for top-N words dispersion: firing-rate drop-off along the noun-preference ranking (Figure S10)

To characterise how rapidly firing declines as one moves from a neuron’s most-preferred word to less-preferred words, we ranked, for each neuron, all words (1221 nouns in Figure 3) by firing rate and normalised the ranked rates by the rank-1 rate, FR(k)/FR(1), for k = 1-100. The population was summarised by the median and interquartile range at each rank, the ratios at ranks 2, 5, 10 and 50, and the rank at which FR(k)/FR(1) first fell below 0.5 (half-maximum rank; set to 101 if this did not occur within the top 100 nouns).

The same profile was computed for concept-cell populations simulated with the framework described above (Figure S3; N = 498 neurons, W = 1221 noun, Gamma-Poisson spike counts with Fano factor 2.66, 5-SD detection threshold, response probability calibrated by bisection to target sparseness α, all ten levels α = 0.2-1.0%; word durations resampled from the 516 unique-noun durations). In the Figure S3 simulation the nouns to which a neuron responds are assigned at random; here each simulated neuron’s evocative nouns were instead set, after calibration, to a randomly chosen seed noun together with its nearest neighbours in static fastText (word2vec) embedding space, keeping the number of evocative nouns per neuron (i.e., α, unchanged), so that each simulated cell responds to a semantically coherent concept. Because repeated tokens of a word share a fastText vector, a cell’s evocative set includes every occurrence of its concept words.

### Neuron tuning across different story-contexts (Figure 4A-C)

To assess whether semantic tuning is stable or context-dependent across different story contexts, we performed the following analyses.

#### Story-specific encoding

The podcast stimulus comprised 6 distinct stories. For each story, we independently fit a ridge regression encoding model (the same ridge regression fitting procedure as Figure 1D) for every neuron, yielding a story-specific tuning vector per neuron.

#### Split-half within-story reliability

To quantify the stability of semantic tuning within a single story, we performed a split-half analysis. For each story, PCA was computed once on all nouns, and the resulting PC scores were then randomly split into two halves. Then we fit separate ridge regression models (the same ridge regression fitting procedure as Figure 1D) on each half, yielding two tuning vectors per neuron per story. The cosine distance between the two tuning vectors was computed as 1 - cos(β_half1_, β_half2_). This split-half procedure was repeated 50 times with different random splits, and the mean cosine distance across repeats was taken as the within-story tuning distance for each neuron-story pair. The final within-story distance for each neuron was averaged across all 6 stories.

#### Across-story tuning distance

Across-story tuning variability was quantified by computing pairwise cosine distances between story-specific tuning vectors (averaged across folds) for each neuron across all pairs of stories.

This across-story distance was compared to the within-story split-half distance: if tuning is context-independent, the two should be comparable; if tuning is modulated by story context, across-story tuning vector distances should exceed within-story tuning vector distances.

*Cross-story representational similarity analysis (Figure 4D-E)*.

To test whether the population-level geometric representation of shared words is preserved across different story contexts, we performed cross-story RSA. The podcast stimulus comprised 6 stories, and many nouns appeared in multiple stories, allowing us to ask whether the same word elicits a similar pattern of population activity regardless of the story in which it occurs. We performed RSA separately for each of the 15 unique story pairs (stories 1-2, 1-3, 1-4, 1-5, 1-6, 2-3, 2-4, 2-5, 2-6, 3-4, 3-5, 3-6, 4-5, 4-6, 5-6) rather than restricting the analysis to words shared across three or more stories. This pairwise approach maximizes statistical power: while only 3 nouns appeared in all 6 stories and 8 appeared in 5 or more, the number of shared nouns per story pair ranged from 8 to 32 (mean ± SD: 19.7 ± 6.5), yielding substantially more pairwise distances for RSA computation. Moreover, restricting to words shared across many stories would bias the sample toward high-frequency, semantically general words (e.g., “thing,” “way,”), undermining the goal of testing whether semantic representations generalize across contexts for a diverse vocabulary.

For each story pair, we identified all nouns that appeared in both stories. For each shared word, we computed a population firing rate vector separately for each story by averaging firing rates across all occurrences of that word within that story for each neuron. We then computed representational dissimilarity matrices (RDMs) for each story using pairwise cosine distances among the shared words’ population vectors and quantified the correspondence between the two story-specific RDMs as the Spearman rank correlation (rho) between the upper-triangular distance vectors. We excluded story pairs with fewer than 3 valid shared words, as fewer than 3 words yield too few pairwise distances (≤ 2) for a meaningful correlation.

#### Null distribution

The null hypothesis was that population geometry reflects general neural state differences between stories rather than word-specific representational structure. To test this, we constructed a null distribution by randomly selecting the same number of (different) words from each story: we drew N words without replacement from one story and N different words from another story, where N equals the number of valid shared words for that pair. Because the two RDMs in each null iteration describe different sets of words, any correlation reflects only incidental similarity in population geometry rather than word-identity-driven structure. We computed RDMs for these mismatched word sets and recorded their Spearman correlation. We repeated this procedure 500 times per story pair.

#### Statistical testing

For each story pair, we computed a permutation p-value as the proportion of null correlations equal to or exceeding the observed correlation. At the group level, we tested whether observed RSA values systematically exceeded null means across all 15 story pairs using the Wilcoxon signed-rank test.

### Lexical context modulation (Figure 4F-K)

To test whether hippocampal semantic tuning is modulated by the local lexical context, we performed the following analyses.

#### Context embedding extraction

For each noun in the stimulus, we computed a context embedding as the mean of the GPT-2 Layer-36 embeddings of the preceding words within a context window, using the original word order in the 7346-word corpus. We defined context window sizes from 1 to 20 preceding words to avoid any bias from choosing a specific window. The context window excluded the target word itself. Words at the beginning of the stimulus with fewer than 20 preceding words used all available preceding context. For the model comparison analysis, we reduced context embeddings to 3 PCs via PCA computed on noun embeddings only; we then projected all words (including non-nouns in the context window) onto these noun-defined PCs to ensure a shared coordinate system.

#### Model comparison (word, context, and interaction effects)

To decompose the contributions of lexical embedding, local context, and their interaction to neural firing rates, three nested linear models were compared for each neuron using:

● M1 (word only): FR ∼ W, where W is the lexical embedding (reduced to low-dimensional PCs, we used the first three PCs)
● M2 (word + context): FR ∼ W + C, where C is the contextual GPT-2 embedding (3 noun-defined PCs of the preceding context window)
● M3 (word + context + interaction): FR ∼ W + C + W×C Where FR is the firing rate of each neuron.

We z-scored and mean-centered all predictors prior to model fitting. We orthogonalized interaction terms (W×C, all pairwise products of W and C dimensions) with respect to the main effects by regressing each interaction column on [1, W, C] and retaining the residuals, ensuring that interaction variance was not confounded with main effects. We quantified model quality using adjusted R², which penalizes model complexity.

#### Standardized beta magnitudes

To compare the relative contribution of word, context, and interaction terms, the mean absolute standardized regression coefficient was computed for each predictor group (|β_W_|, |β_C_|, |β_W×C_|) in M3, as well as (|β_W_|, |β_C_|) in M2 for each neuron. Pairwise comparisons were performed using paired t-tests across neurons.

#### Phonetic regression

We built a fixed-length feature space over the 43 IPA phonemes observed in our corpus. Here, IPA refers to the International Phonetic Alphabet, a standardized set of symbols that denote speech sounds independent of spelling. The podcast transcript was processed word by word: tokens were lower-cased and converted from graphemes to phonemes using the Python phonemizer package (version 3.0.1) with the eSpeak backend as the grapheme-to-phoneme (G2P) model (https://pypi.org/project/phonemizer/3.0.1/). For each word, we took the canonical IPA transcription returned by phonemizer. Phonemes were then manually inspected and corrected by linguists (JB) in Praat.

Each word was then represented as a 43-dimensional count vector whose entries reflect the number of occurrences of each IPA phoneme in that word (unigram counts; no duration or stress information). This approach is similar to Khanna et al., in that words are represented by the presence of their constituent phonemes; however, we did not group phonemes into broader classes, in order to preserve the full identity of the 43 phones observed in our data and maintain the embedding’s informational richness and dimensionality (Khanna et al., 2024). To better differentiate words that contain the same phoneme set but in different orders (e.g., dog vs god), we double-counted the first phoneme in every word while counting all subsequent phonemes once. The resulting word-by-phoneme matrix (N words × 43 features) was used in subsequent analyses relating phonetic composition to neural responses.

These results were unchanged when we repeated the model comparison with phonetic embeddings, word frequency, GPT-2 surprisal and entropy, pitch, loudness, and story identity added as covariates: M2 outperformed M1 and M3 outperformed M2 at all 20 context window sizes (all p < 0.001, BH-FDR-corrected). Word duration was not included as a covariate because firing rates are already normalized by word duration.

### Context-driven pattern separation (Figure 5A-D)

We tested whether the hippocampus performs context-driven *pattern separation* by asking whether repeated occurrences of the same word evoke neural population patterns that reflect differences in their preceding lexical context. If the hippocampus performs context-driven pattern separation, then occurrences of the same word (e.g., “people” appearing five times) should produce more dissimilar population responses when their preceding contexts are more dissimilar. In contrast, if word representations are strictly context-invariant (as predicted by a concept-cell model), repeated occurrences should evoke similar neural patterns regardless of context.

#### Context firing rate extraction

For each word at position *t* in the original 7346-word corpus, we computed a context firing rate vector as the mean population firing rate across the preceding *w* words

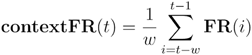

where FR(i) ∈ ℝ^498^ is the population firing rate vector at position *i* and *w* is the context window size. The target word itself was excluded. For words near the beginning of the corpus with fewer than win-size preceding words, all available preceding words were used. Context extraction operated on the original corpus ordering (prior to noun selection), ensuring that non-noun words contributed to the context representation. Words with no preceding context (position = 1) were assigned a zero vector. The context firing rate vectors were stored separately for each occurrence of each repeated word type, preserving the word-level variability needed for the context-driven pattern separation analysis. This procedure was repeated independently for context window sizes *w* = 1 to 20.

#### Repeated words and their context geometry coupling

For each noun type with ≥ 5 occurrences, we extracted two matrices across the *k* occurrences of that word: a context matrix **C** (*k* × 498 neurons), where each row is the mean population firing rate vector of the preceding context window (see *Context firing rate extraction* above), and a target word matrix **W** (*k* × 498 neurons), where each row is the population firing rate evoked by that occurrence. We computed pairwise cosine distances separately within **C** and **W**, yielding a context RDM and a target word RDM each of size *k\**(*k*−1)/2. We then calculated the Spearman rank correlation between these two distance vectors to quantify the degree of context-driven pattern separation: a positive correlation indicates that occurrences preceded by distinct contexts also elicited distinct neural responses.

#### Constructing baseline with random context

To confirm that observed coupling reflects specific semantic context rather than generic neural state fluctuations, we replaced actual preceding context firing rates with firing rates from randomly sampled words elsewhere in the recording. On each of 500 null iterations per word type, we drew k words without replacement from the full noun pool (1221 nouns × 498 neurons), assembled their firing rates into a surrogate context matrix, and recomputed the coupling between repeated words and random context geometry as the null baseline. At the group level, we tested the difference between observed and random-context coupling using a Wilcoxon signed-rank test. We repeated both analyses for context window sizes from 1 to 20 preceding words to avoid any bias from choosing a specific window

### Generalization performance across semantic dimensions (Figure 5E-H)

To test whether hippocampal population activity encodes semantic dimensions in an abstract, generalizable format, we computed generalization performance for each principal component (100 PCs in total) of the GPT-2 embedding space, borrowing the framework of cross-condition generalization performance (Bernardi et al., 2020).

#### Semantic dimension definition

GPT-2 Layer-36 embeddings for all noun words were z-scored and reduced via PCA. For each of the first 100 PCs, words were binarized into “high” (top 30th percentile of PC scores) and “low” (bottom 30th percentile) groups.

#### Generalization performance computation

For each PC, we trained a linear Support Vector Machine (SVM, MATLAB “fitcsvm”, linear kernel) to classify words as high versus low using their population firing rate vectors. On each of 20 repetitions, 70% of words from each group were randomly assigned to training and 30% to testing (minimum 3 test words per group). Generalization performance was defined as the mean classification accuracy across repetitions. The critical feature of this approach is that the classifier must generalize across different word exemplars: because training and test sets contain non-overlapping words, the SVM cannot succeed by memorizing the firing rate pattern of any specific word. Instead, above-chance classification requires that the population geometry encodes the semantic dimension in a linearly separable format that holds across arbitrary subsets of the vocabulary. This is a more stringent test than standard decoding, which can exploit stimulus-specific patterns. If a semantic dimension is encoded through a distributed, abstract representation, where the population vector systematically varies along that dimension regardless of which particular words are sampled, then a linear hyperplane learned from one subset of words will correctly classify novel words. Conversely, if the representation is entangled with word-specific features, a linear classifier trained on one subset will fail to generalize to held-out exemplars. Generalization performance thus directly tests whether semantic dimensions are represented in the low-dimensional, abstractly formatted geometry predicted by recent theoretical frameworks for flexible generalization (Bernardi et al., 2020).

#### Shuffle baseline

To establish a null distribution, we repeated the full 20-repetition procedure 50 times with randomly permuted group assignments: on each shuffle, the same number of words were assigned to “high” and “low” groups irrespective of their true PC scores. The p-value for each PC was computed as the proportion of shuffle values greater than or equal to the real generalization performance. PCs with p < 0.05 were considered to have significantly generalizable representations.

#### Shattering dimensionality

To quantify the overall representational capacity of the population, we computed shattering dimensionality (Bernardi et al., 2020), defined as the proportion of tested semantic dimensions for which generalization performance significantly exceeded the shuffle baseline (p < 0.05). A shattering dimensionality of 1 would indicate that all 100 semantic dimensions are abstractly and linearly decodable; the observed value provides an estimate of how many independent semantic dimensions are represented in a generalizable format in the population code.

Generalization performance significantly above the shuffle baseline indicates that the hippocampal population represents semantics in a high-dimensional and structured format that generalizes across different word exemplars, a hallmark of abstract, linearly decodable coding geometry.

#### Replication dataset: naturalistic dialogue listening

To test whether the coding properties observed during story listening generalize to a different speech genre, we analyzed hippocampal recordings from 6 participants who listened to a 46-min multi-speaker conversational video (an episode of The Try Guys podcast). The recording, spike sorting and unit-inclusion criteria were identical to the main dataset (see above), yielding 217 well-isolated hippocampal neurons. A ground-truth word-level transcript with onset and offset times was obtained by forced alignment of a manually corrected transcript, giving 10,217 words, 1226 nouns and 486 unique nouns.

## Acknowledgements

We thank Joshua Adkinson, Justin Fine, Victoria Gates, Suzanne Kemmer, George Kokalas, and Steven Piantadosi for their assistance.

## Supplementary Information

### Supplementary Table

**Supplementary Table 1.**
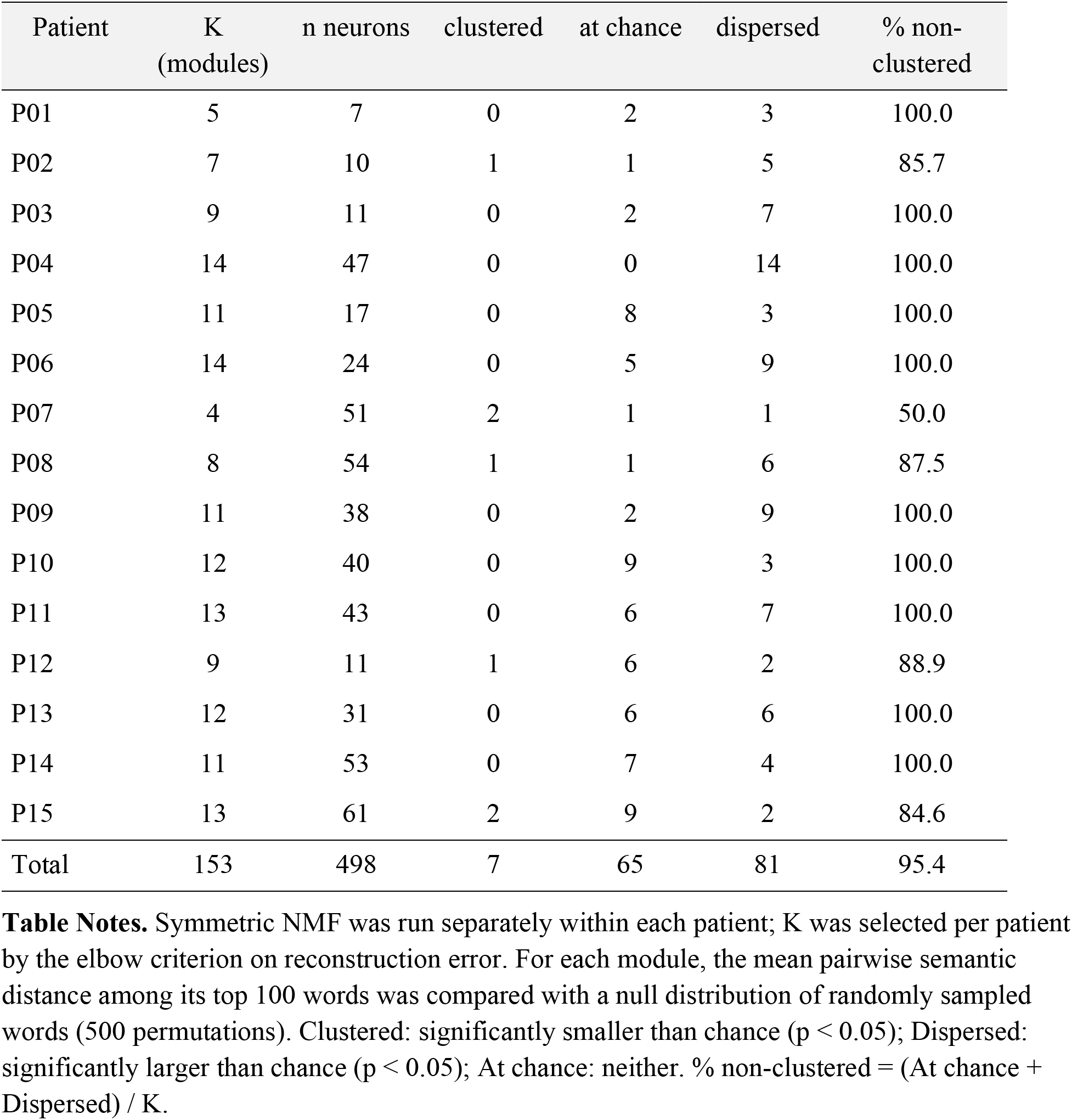
Per-patient symmetric NMF modules and their semantic dispersion.

| Patient | K (modules) | n neurons | clustered | at chance | dispersed | % non-clustered |
|---|---|---|---|---|---|---|
| P01 | 5 | 7 | 0 | 2 | 3 | 100.0 |
| P02 | 7 | 10 | 1 | 1 | 5 | 85.7 |
| P03 | 9 | 11 | 0 | 2 | 7 | 100.0 |
| P04 | 14 | 47 | 0 | 0 | 14 | 100.0 |
| P05 | 11 | 17 | 0 | 8 | 3 | 100.0 |
| P06 | 14 | 24 | 0 | 5 | 9 | 100.0 |
| P07 | 4 | 51 | 2 | 1 | 1 | 50.0 |
| P08 | 8 | 54 | 1 | 1 | 6 | 87.5 |
| P09 | 11 | 38 | 0 | 2 | 9 | 100.0 |
| P10 | 12 | 40 | 0 | 9 | 3 | 100.0 |
| P11 | 13 | 43 | 0 | 6 | 7 | 100.0 |
| P12 | 9 | 11 | 1 | 6 | 2 | 88.9 |
| P13 | 12 | 31 | 0 | 6 | 6 | 100.0 |
| P14 | 11 | 53 | 0 | 7 | 4 | 100.0 |
| P15 | 13 | 61 | 2 | 9 | 2 | 84.6 |
| Total | 153 | 498 | 7 | 65 | 81 | 95.4 |

**Table Notes.** Symmetric NMF was run separately within each patient; K was selected per patient by the elbow criterion on reconstruction error. For each module, the mean pairwise semantic distance among its top 100 words was compared with a null distribution of randomly sampled words (500 permutations). Clustered: significantly smaller than chance (p < 0.05); Dispersed: significantly larger than chance (p < 0.05); At chance: neither. % non-clustered = (At chance + Dispersed) / K.

**Supplementary Table 2.** Per-patient symmetric NMF modules and their semantic dispersion for verbs and adjectives.

| Patient | n neurons | K (modules) | Clustered | At chance | Dispersed | % non-clustered |
|---|---|---|---|---|---|---|
| *Verbs* |  |  |  |  |  |  |
| P01 | 7 | 4 | 1 | 3 | 0 | 75.0 |
| P02 | 10 | 8 | 0 | 2 | 6 | 100.0 |
| P03 | 11 | 6 | 0 | 3 | 3 | 100.0 |
| P04 | 47 | 13 | 3 | 2 | 8 | 76.9 |
| P05 | 17 | 11 | 5 | 5 | 1 | 54.5 |
| P06 | 24 | 12 | 0 | 9 | 3 | 100.0 |
| P07 | 51 | 11 | 6 | 5 | 0 | 45.5 |
| P08 | 54 | 14 | 7 | 4 | 3 | 50.0 |
| P09 | 38 | 11 | 1 | 10 | 0 | 90.9 |
| P10 | 40 | 3 | 0 | 0 | 3 | 100.0 |
| P11 | 43 | 13 | 0 | 8 | 5 | 100.0 |
| P12 | 11 | 5 | 1 | 3 | 1 | 80.0 |
| P13 | 31 | 14 | 3 | 10 | 1 | 78.6 |
| P14 | 53 | 10 | 1 | 7 | 2 | 90.0 |
| P15 | 61 | 14 | 0 | 11 | 3 | 100.0 |
| **Total** | **498** | **149** | **28** | **82** | **39** | **81.2** |
| *Adjectives* |  |  |  |  |  |  |
| P01 | 7 | 5 | 0 | 4 | 1 | 100.0 |
| P02 | 10 | 7 | 0 | 4 | 3 | 100.0 |
| P03 | 11 | 9 | 0 | 1 | 8 | 100.0 |
| P04 | 47 | 10 | 0 | 3 | 7 | 100.0 |
| P05 | 17 | 12 | 0 | 11 | 1 | 100.0 |
| P06 | 24 | 12 | 0 | 11 | 1 | 100.0 |
| P07 | 50 | 14 | 0 | 13 | 1 | 100.0 |
| P08 | 54 | 10 | 0 | 9 | 1 | 100.0 |
| P09 | 38 | 14 | 0 | 9 | 5 | 100.0 |
| P10 | 40 | 12 | 7 | 5 | 0 | 41.7 |
| P11 | 43 | 14 | 6 | 8 | 0 | 57.1 |
| P12 | 11 | 9 | 1 | 4 | 4 | 88.9 |
| P13 | 31 | 8 | 0 | 3 | 5 | 100.0 |
| P14 | 53 | 14 | 3 | 11 | 0 | 78.6 |
| P15 | 61 | 14 | 1 | 13 | 0 | 92.9 |
| **Total** | **497** | **164** | **18** | **109** | **37** | **89.0** |

**Table Notes.** Same procedure as Table S1, applied separately to verbs and adjectives. Symmetric NMF was run within each patient; K was selected per patient by the elbow criterion on reconstruction error. For each module, the mean pairwise semantic distance among its top 100 words was compared with a null distribution of randomly sampled words (500 permutations). Clustered: significantly smaller than chance (p < 0.05); Dispersed: significantly larger than chance (p < 0.05); At chance: neither. % non-clustered = (At chance + Dispersed) / K.

**Supplementary Table 3.** Validation analyses for polysemanticity and overdispersion (n = 498 neurons, 100 random occurrence splits).

| **K** | **Replication of top-K clusters** |  | **Overdispersion of replicated clusters** |  |  |  |
|---|---|---|---|---|---|---|
|  | **Percentile in B** | **p** | **Distance (top)** | **Distance (rand.)** | **d** | **p** |
| 5 | 52.8 | 5.8 × 10<sup>-9</sup> | 0.565 | 0.515 | 0.41 | 2.7 × 10<sup>-18</sup> |
| 7 | 53.0 | 1.5 × 10<sup>-13</sup> | 0.556 | 0.515 | 0.35 | 2.9 × 10<sup>-14</sup> |
| 10 | 53.3 | 3.9 × 10<sup>-21</sup> | 0.536 | 0.515 | 0.20 | 1.6 × 10<sup>-5</sup> |

Table Notes. Overdispersion: mean pairwise cosine distance between replicated top clusters (percentile > 50 in half B) vs. size-matched sets of randomly drawn clusters; paired t-test; d = Cohen’s d. All K values were evaluated within identical data partitions; statistics were averaged across the 100 splits before population-level testing.
Related to Figure 3.

### Supplementary Figures

**Figure S1.**
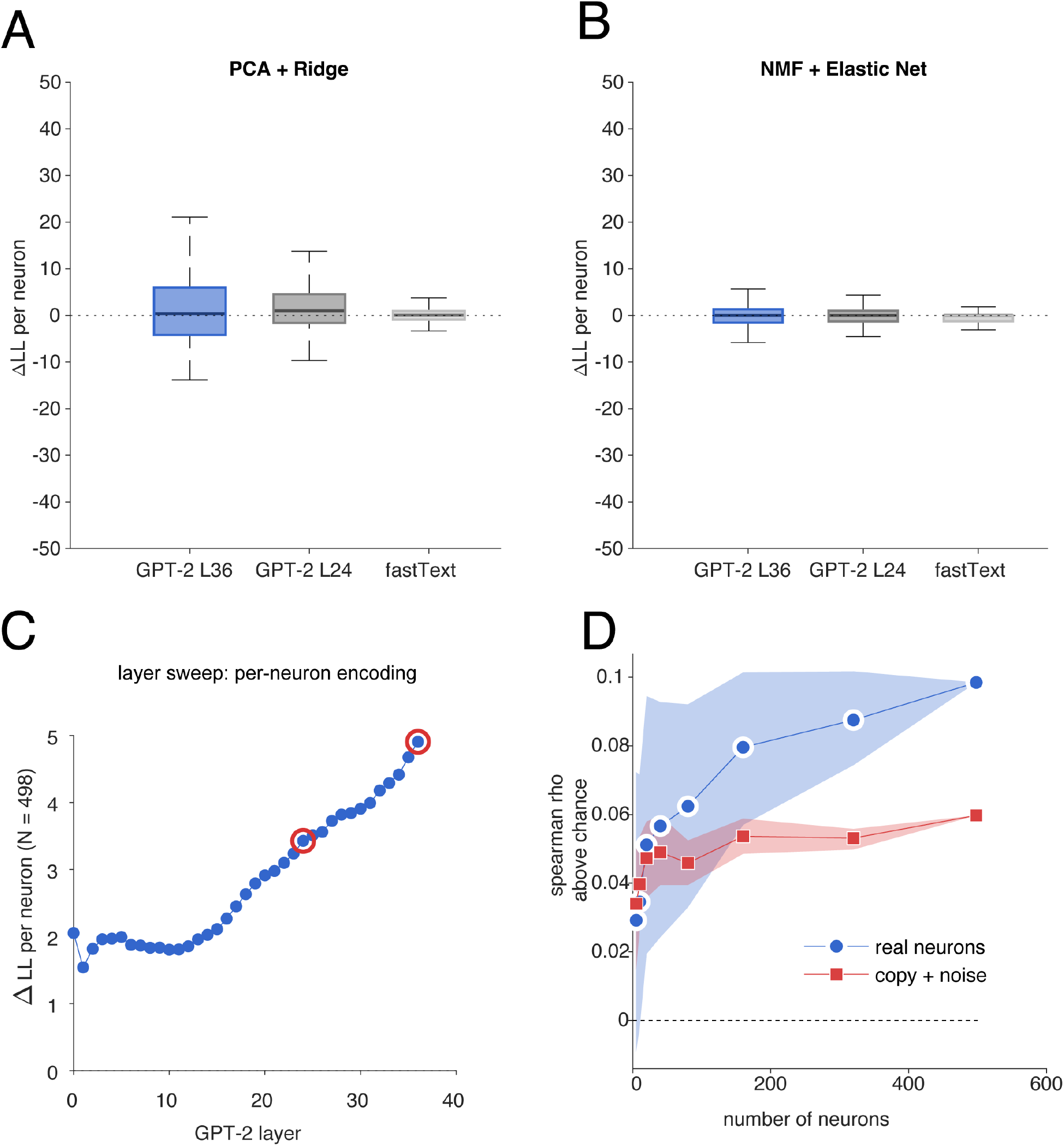
Control analyses for encoding pipeline and RSA scaling. (A, B) One potential limitation is that our encoding pipeline, PCA for dimensionality reduction paired with ridge regression, could favor distributed solutions by construction: PCA produces dense, signed components that spread variance across all dimensions, and ridge regression penalizes large weights, discouraging the sparse, high-magnitude tuning expected under a concept-cell scheme. To address this, we conducted control analyses using a sparse-friendly alternative: non-negative matrix factorization (NMF; Lee and Seung, 1999) for embedding decomposition, which yields parts-based, non-negative components that naturally accommodate sparse representations, paired with elastic net regression (L1 ratio = 0.9; see **Methods**), which drives irrelevant coefficients to exactly zero. Layer 36 again provided the strongest fit, and the distributed pipeline marginally outperformed the sparse alternative (paired t test, t(497) = 1.81, p = 0.071), indicating that the encoding results are not an artifact of pipeline choice but reflect the underlying structure of the neural code. (C) We report cross-validated encoding performance (ΔLL) for all 37 GPT-2 layers (inclusive of Layer 0). As we can see, Layer 36 is the best-performing, but Layer 24, while not as good, is also not that bad. (D) RSA decimation scaling: real neurons vs. copy-plus-noise control. Spearman rank correlation (ρ above chance) between hippocampal and GPT-2 Layer 36 representational dissimilarity matrices (cosine distance) as a function of the number of neurons included in the population. For each population size N ∈ {5, 10, 20, 40, 80, 160, 320, 498}, random subsets of N neurons were drawn without replacement from the full population of 498 units, and the pairwise cosine distance matrix over nouns was computed and compared to the corresponding GPT-2 RDM via Spearman correlation. This procedure was repeated 200 times per population size. Chance level was estimated by a shuffle control (50 iterations per N) in which the mapping between words and neural firing rates was randomly permuted; plotted values reflect the mean ρ minus the shuffle mean at each N. Shaded regions indicate 95% confidence intervals (2.5th-97.5th percentiles) of the neuron-resampling distribution. The copy-plus-noise control (red) was constructed by identifying the single neuron with the highest cross-validated encoding performance (ΔLL), decomposing its activity into a signal component (ridge-predicted) and residual noise, and generating 498 synthetic neurons by adding independent Gaussian noise (variance matched to the empirical residuals) to copies of the signal. The same decimation procedure was then applied to this synthetic population (100 iterations per N). Real neurons (blue) show monotonically increasing RSA with population size, whereas the copy-plus-noise control plateaus, confirming that additional neurons carry non-redundant semantic information rather than simply averaging out shared noise. At N = 498, the confidence interval collapses to zero width because all neurons are included in every iteration, leaving no variability across resampled subsets. Related to Figure 1.

**Figure S2.**
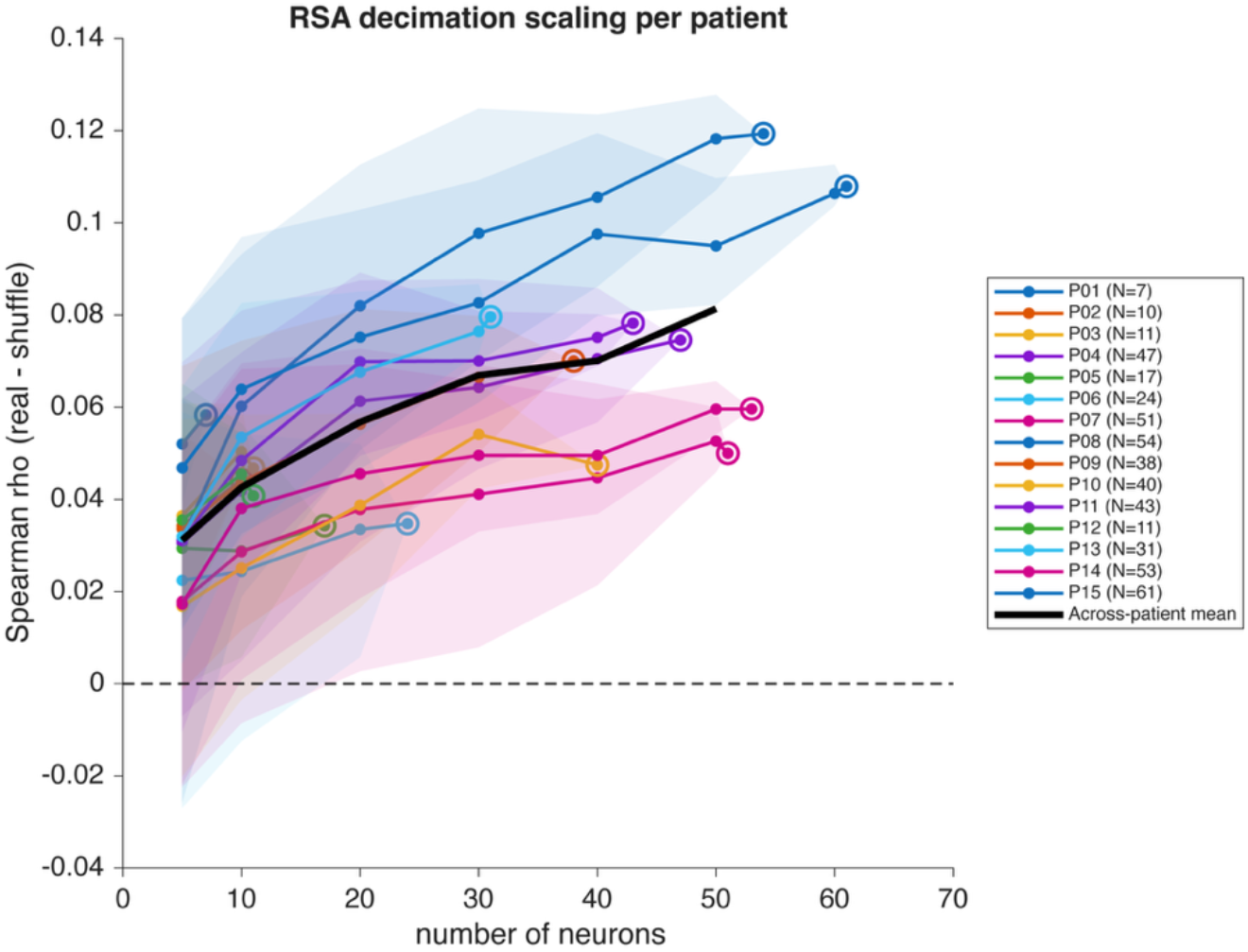
RSA decimation scaling within individual patients. Spearman correlation between the neural representational dissimilarity matrix and the GPT-2 Layer-36 embedding RDM (cosine distance; 1221 nouns), as a function of the number of hippocampal neurons sampled from a single patient. Values are shuffle-corrected (real ρ minus mean ρ after permuting word identity). For each patient, subsets of N = 5, 10, 20, 30, 40, 50, 60 neurons (truncated at the patient’s total). The patient’s full population is included as the final point (open circle). Lines show the mean and shaded bands the 2.5th-97.5th percentiles across subsets. Black line: mean across patients at each N, computed over patients with at least N neurons. Shuffle-corrected ρ was positive at every patient’s full population (range 0.034-0.119) and increased monotonically with N in all patients with three or more subset sizes (within-patient Spearman correlation between N and shuffle-corrected ρ; Wilcoxon signed-rank test of these correlations against 0, p < 0.001). Related to Figure 1.

**Figure S3.**
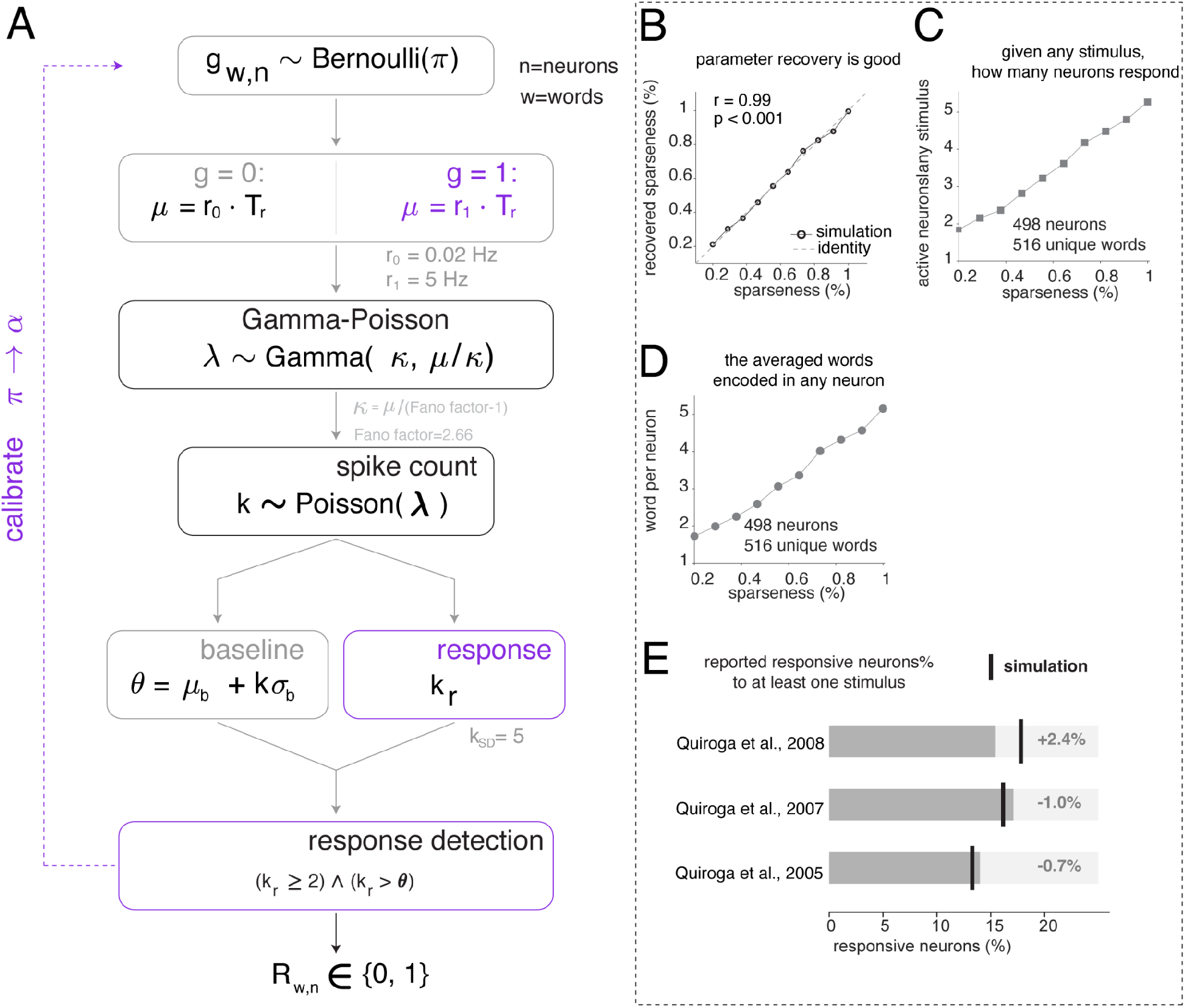
Concept cell simulation framework and validation results. (A) Concept cell simulation framework (Method). To benchmark the observed sparseness against theoretical concept-cell predictions, we developed a generative model calibrated to published concept-cell parameters and naturalistic listening constraints. For each neuron-word pair, a Bernoulli gate (with probability π, the target sparseness) determines whether the neuron is “responsive” to that word. Responsive pairs fire at an elevated rate (r_1_ = 5 Hz) and non-responsive pairs at baseline (r_0_ = 0.02 Hz), with trial-to-trial variability generated by a Gamma-Poisson process (Fano factor = 2.66). Spike counts are converted to firing rates using the word’s duration, and a response is detected if the spike count exceeds a threshold defined as baseline mean plus k standard deviations (k_SD_ = 5). This yields a binary response matrix (words × neurons) from which sparseness metrics are computed. (B) Simulation parameter recovery. To validate that the simulation accurately implements the intended sparseness level, recovered sparseness (measured from the simulated response matrix) is plotted against the input sparseness parameter (π) across a range of target values. The near-perfect correlation (r = 0.99, p < 0.001) confirms reliable parameter recovery. (C) Number of responsive neurons per stimulus. To assess how population recruitment scales with coding sparseness, the average number of neurons responding to any given stimulus is plotted as a function of target sparseness (498 neurons, 516 unique words). At the sparseness levels reported in the concept-cell literature, only 1-2 neurons respond per word, whereas real data show substantially broader recruitment. (D) Number of words encoded per neuron. To assess stimulus selectivity, the average number of words eliciting a response in any given neuron is plotted as a function of target sparseness. At concept-cell sparseness levels, each neuron responds to approximately 1 word, whereas real neurons respond to many more. (E) Comparison with published concept-cell benchmarks. To anchor the simulation to the existing literature, the percentage of responsive neurons reported in three published concept-cell studies (horizontal bars) is shown alongside the simulation’s predictions at corresponding sparseness levels. The vertical black line indicates the stimulus presentation threshold used in each study. The close agreement between published values and simulation output validates the model’s calibration to the concept-cell literature. Related to Figure 2.

**Figure S4.**
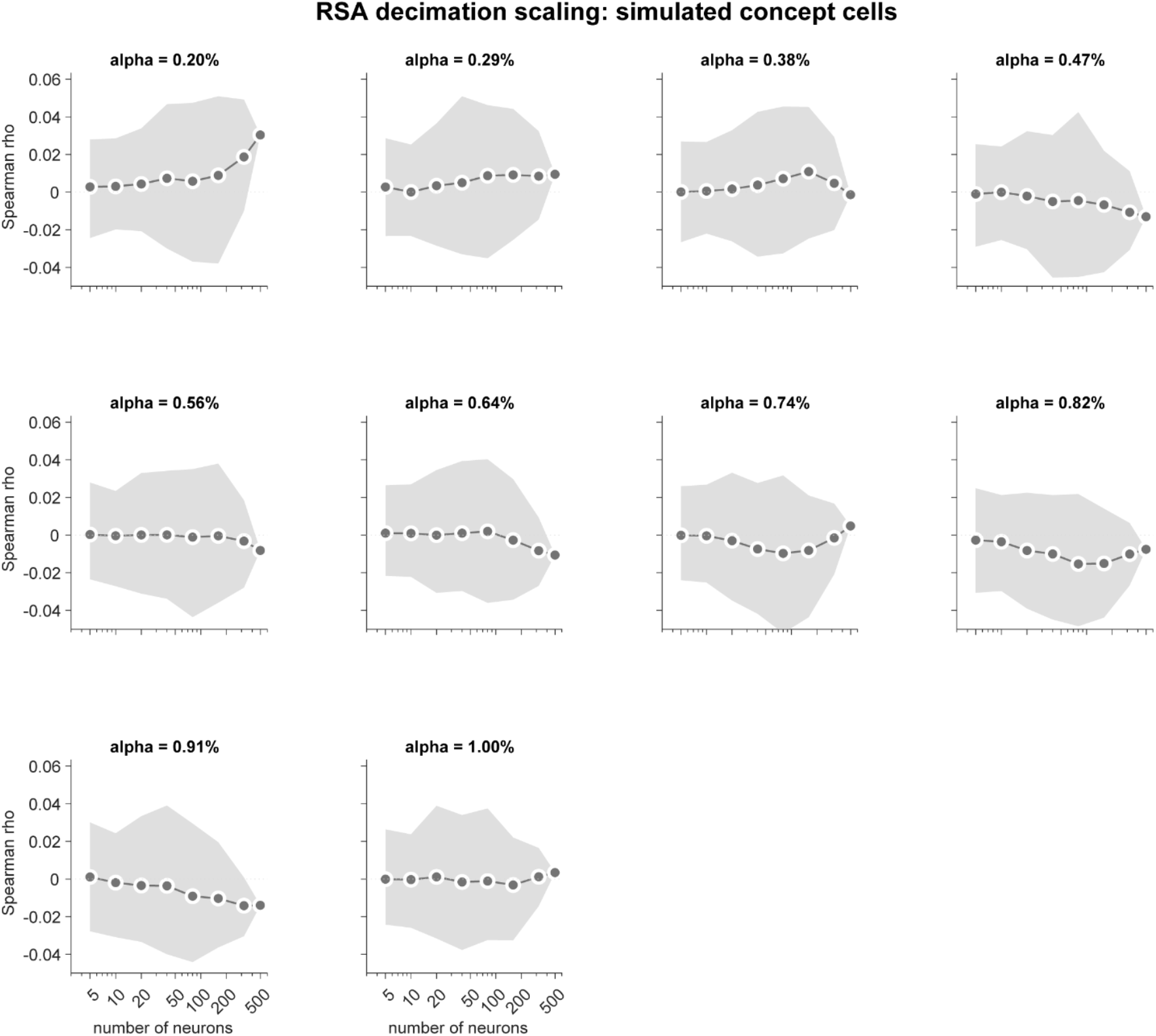
RSA decimation scaling for simulated concept cell populations. Each panel shows the Spearman rank correlation between the neural and GPT-2 Layer 36 pairwise cosine distance matrices as a function of the number of neurons included in the analysis, for simulated concept cell populations at a given sparseness level (response probability alpha (sparseness level) ranging from 0.20% to 1.00%). Grey lines and dots indicate mean correlation across 200 random subsamples; shaded regions indicate 95% confidence intervals. Across all sparseness levels and population sizes, simulated concept cells produce near-zero RSA correlations (rho range: -0.014 to 0.030 at N = 498), with no significant monotonically increasing trend as population size grows. This result indicates that sparse, concept-cell-like populations cannot reconstruct the continuous semantic geometry of the language model embedding space. Compare with the monotonically increasing RSA observed in the real hippocampal population (Figure 1H). Related to Figure 1.

**Figure S5.**
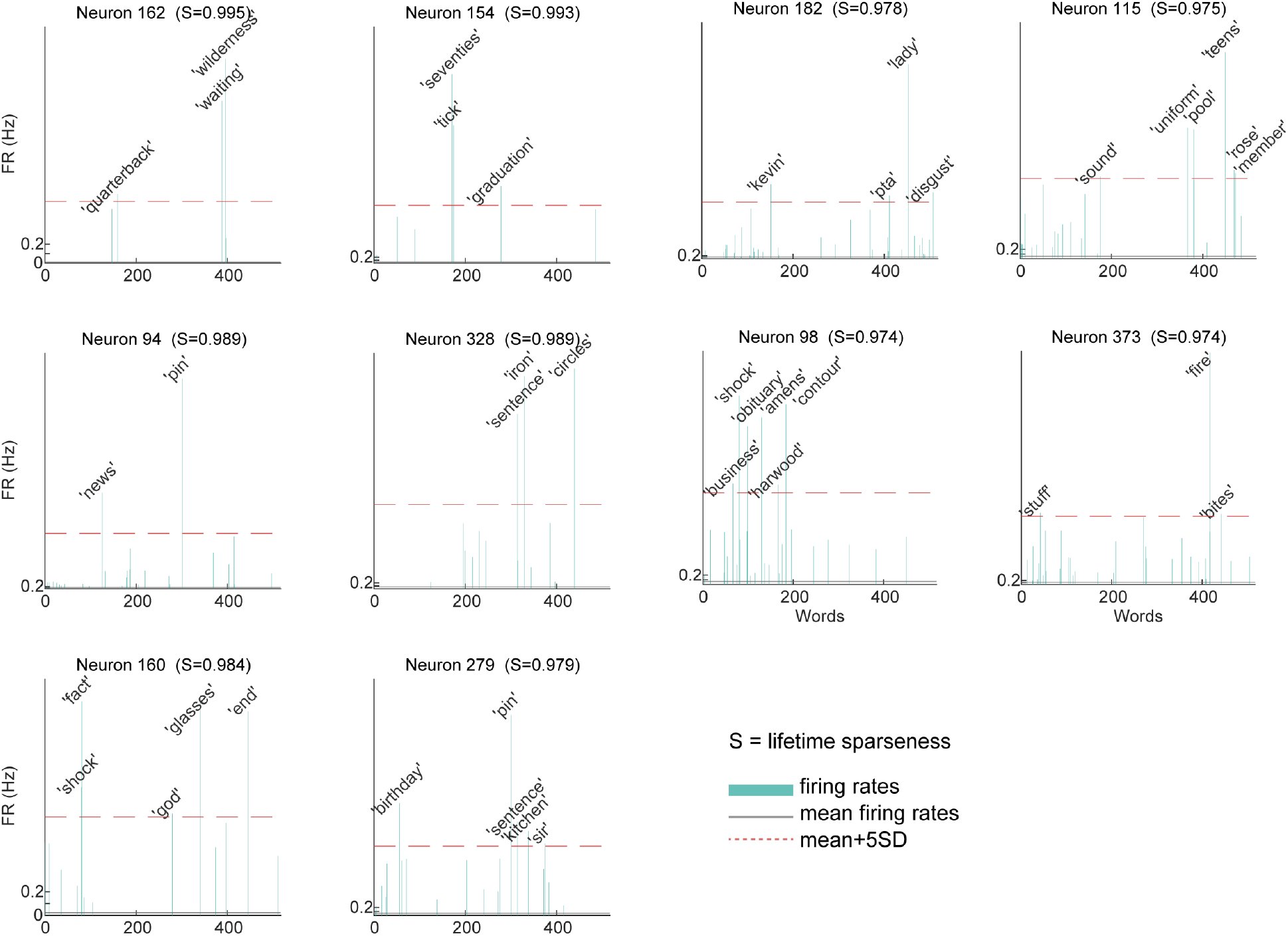
Concept-cell-like neurons identified by lifetime sparseness in language processing. Firing rate profiles across all 516 unique nouns for the 10 neurons with the highest lifetime sparseness (top 2%, S ≥ 0.974). Each panel shows one neuron’s firing rate (teal bars) to each word in its original presentation order. The solid grey line indicates the mean firing rate and the dashed red line indicates the mean + 5 SD threshold. Words exceeding this threshold are labeled. These neurons exhibit sparseness values above 0.97, responding strongly to only 2-6 words out of 516. Notably, the words driving each neuron’s responses span semantically distant categories (e.g., Neuron 162: “quarterback”, “waiting”, “wilderness”; Neuron 160: “shock”, “fact”, “god”, ‘glasses’, “end”). Related to Figure 2.

**Figure S6.**
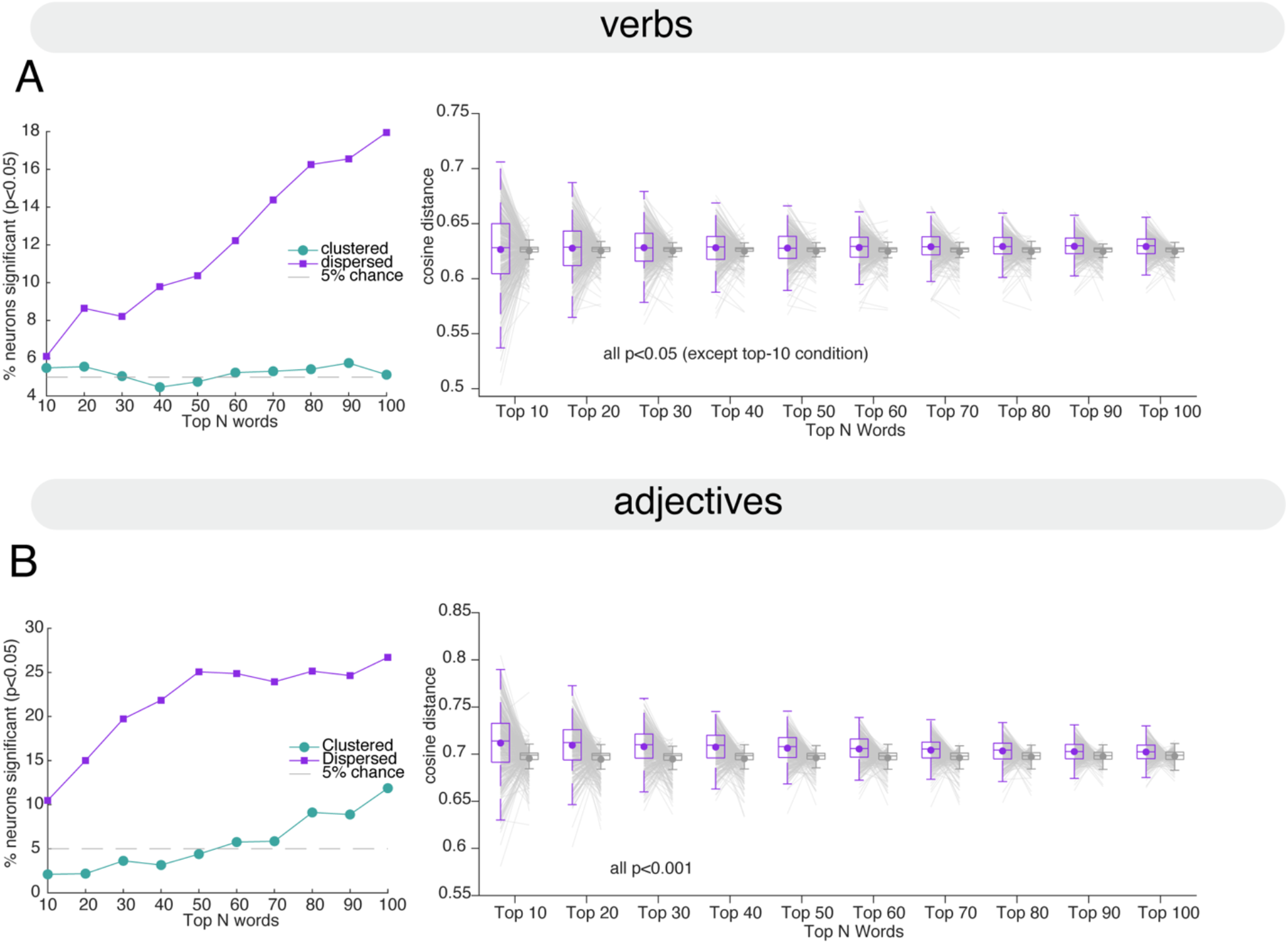
Overdispersion results held for both verbs and adjectives. (A) **Verbs**. Left panel: proportion of neurons showing significant dispersion vs. clustering across top N words. Overdispersion dominates at all N values. Right panel: mean cosine distance for top N words vs. random N words. Top-preferred words remain significantly more dispersed than chance (except the N=10, for all other conditions, paired t-tests, all t >2.183, p < 0.05). (B) **Adjectives.** Left panel: proportion of neurons showing significant dispersion vs. clustering across top N words. Overdispersion dominates at all N values. Right panel: mean cosine distance for top N words vs. random N words. Top-preferred words remain significantly more dispersed than chance (for all conditions, paired t-tests, all t > 4.830, p < 0.001). Related to Figure 3.

**Figure S7.**
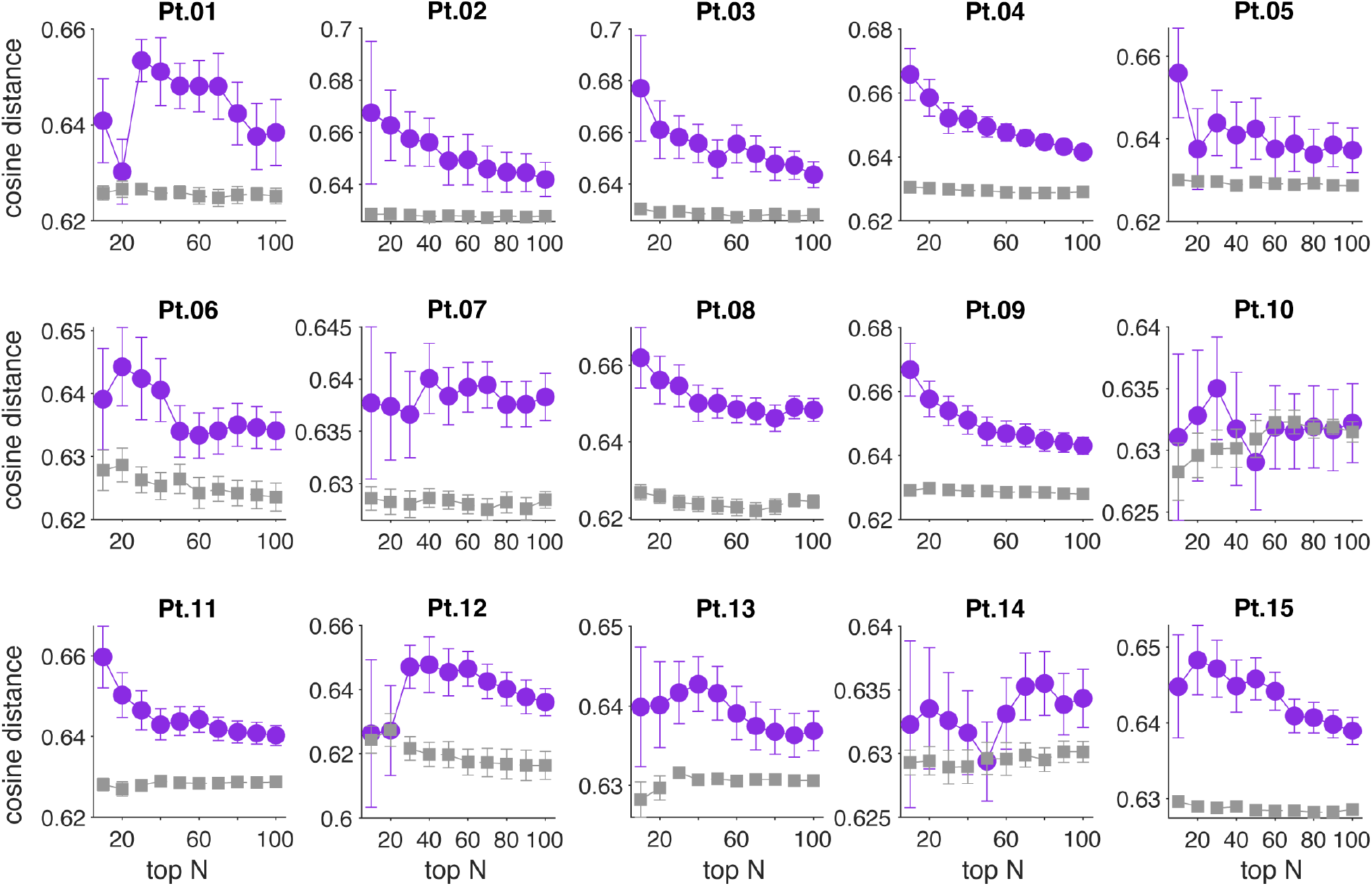
Mean cosine distance for top N words vs. random N words for individual patients. Per-patient replication of the group-level semantic dispersion analysis. For each patient, the mean pairwise cosine distance among each neuron’s top N preferred words (purple) is plotted alongside the null expectation from random words (gray, mean of 500 permutations) across top N values (N = 10-100). Error bars denote ± SEM across neurons. Results are broadly consistent with the group-level finding: in the majority of patients, top-preferred words exhibit greater semantic dispersion than random words, confirming that the preference for semantically diverse stimuli is not driven by a subset of patients but is a general property of hippocampal neurons across individuals. Related to Figure 3.

**Figure S8.**
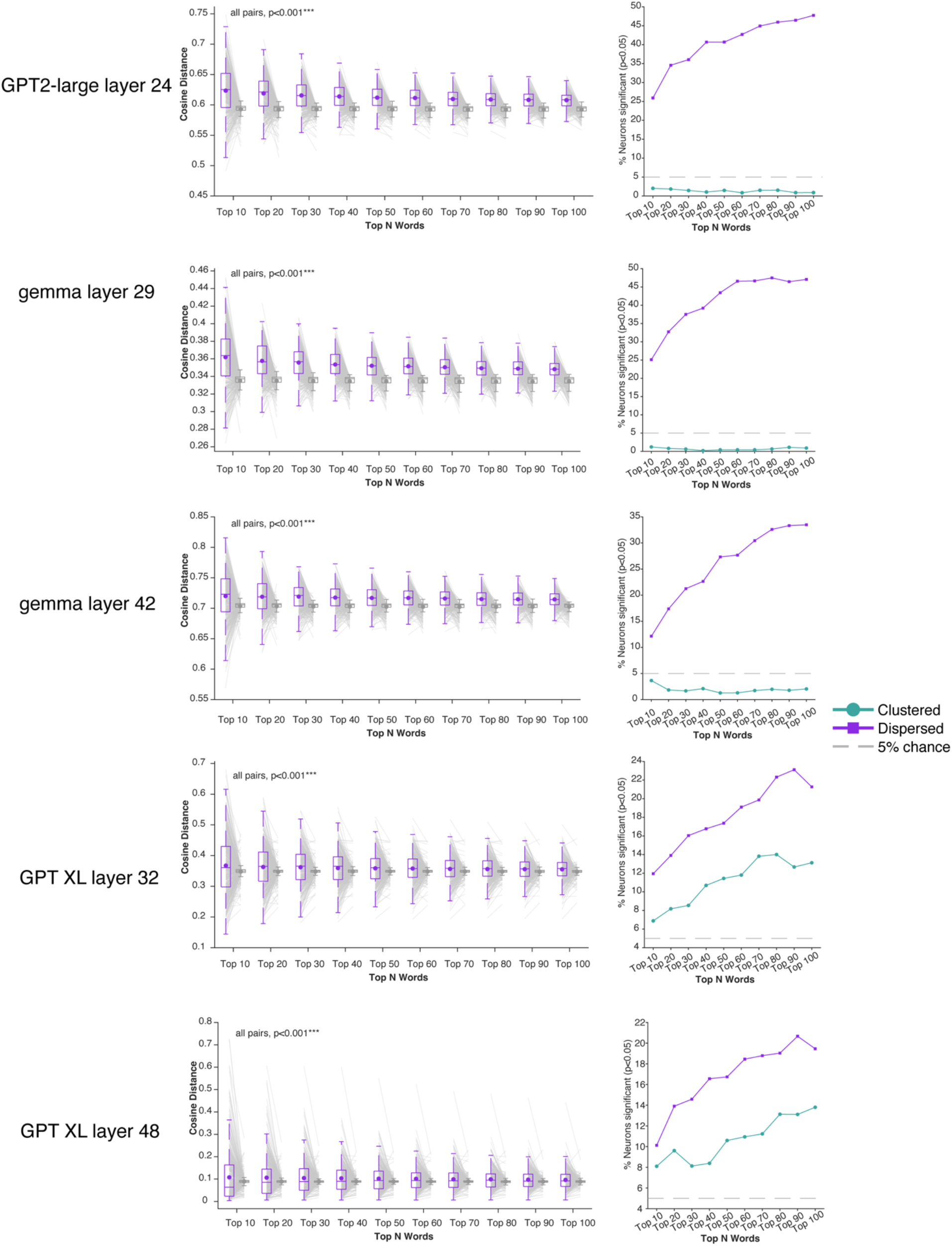
Overdispersion results held for other LLMs and other layers. Left column: Top-N words are significantly more dispersed than random words at every N (all p < 0.001). Right column: Proportion of significantly dispersed neurons exceeds both chance (5%) and clustered neurons at all N values.

**Figure S9.**
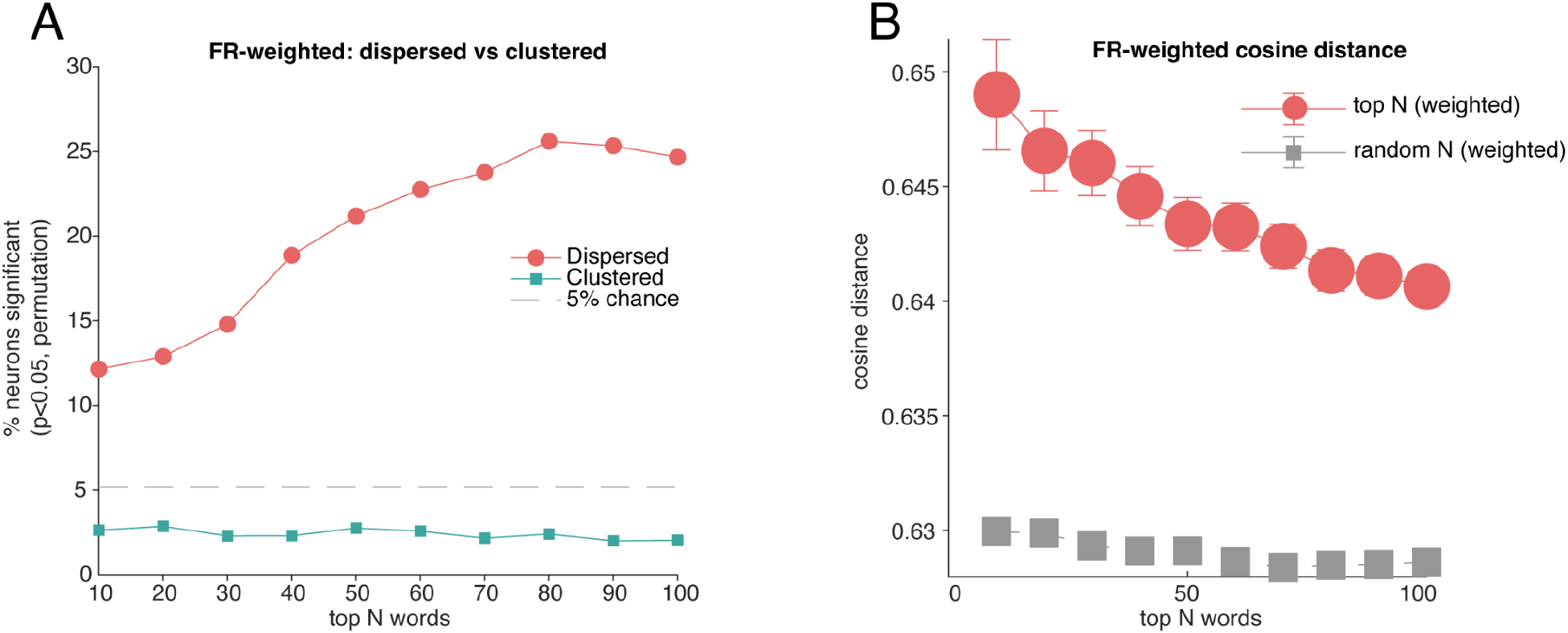
Results from the control analyses for top-N words dispersion. The weighted analysis produced results nearly identical to the unweighted analysis. (A) Under firing-rate weighting, the proportion of dispersed neurons (red line) substantially exceeds both the 5% chance level (gray line) and the proportion of clustered neurons at all values of N. (B) At every value of N from 10 to 100, the FR-weighted mean pairwise distance among each neuron’s top N words was significantly greater than the weighted null expectation (all t > 7.87, all *p* < 0.001). Related to Figure 3.

**Figure S10.**
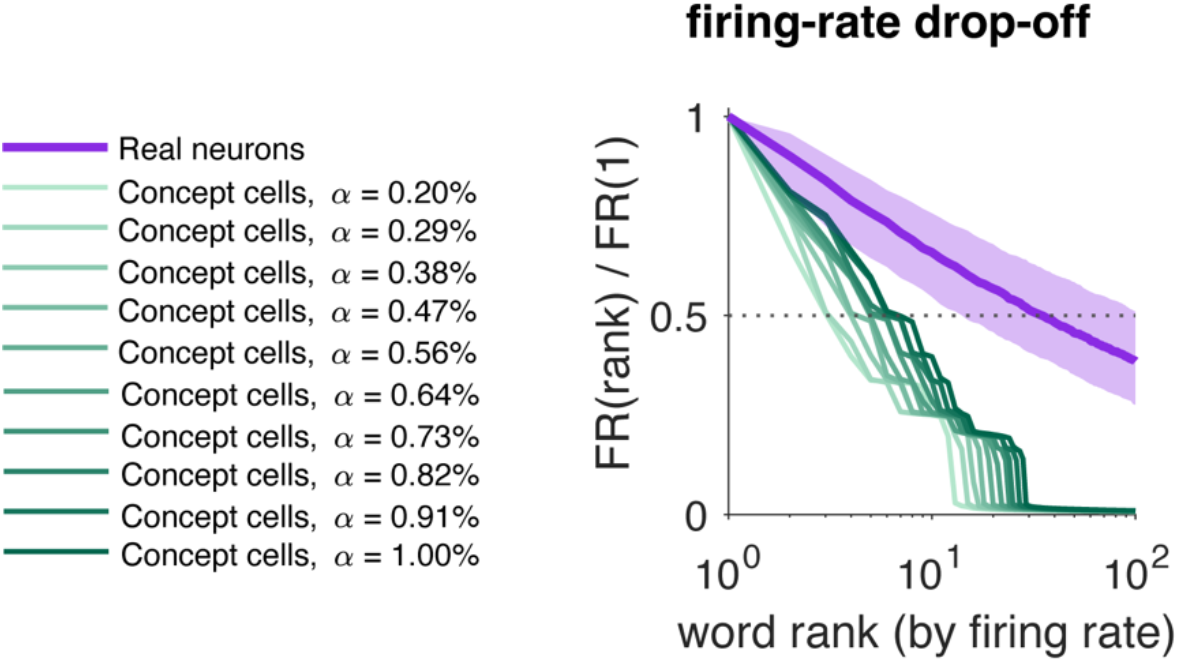
Firing-rate drop-off along the noun-preference ranking in real hippocampal neurons and simulated concept cells. Firing rate as a function of noun rank, normalized to each neuron’s rank-1 rate (median; shaded band, interquartile range for real neurons; log x-axis). Purple: real neurons (n = 498). Green: concept cells simulated with the Figure S3 framework (n = 498 per level; light to dark, α = 0.2-1.0%), each responding to a semantically coherent concept (seed noun and its nearest fastText neighbours). Real neurons decline gradually, falling below half-maximum (dotted line) at a median rank of 35; simulated concept cells fall below half-maximum by rank 4-7. Related to Figure 3.

**Figure S11.**
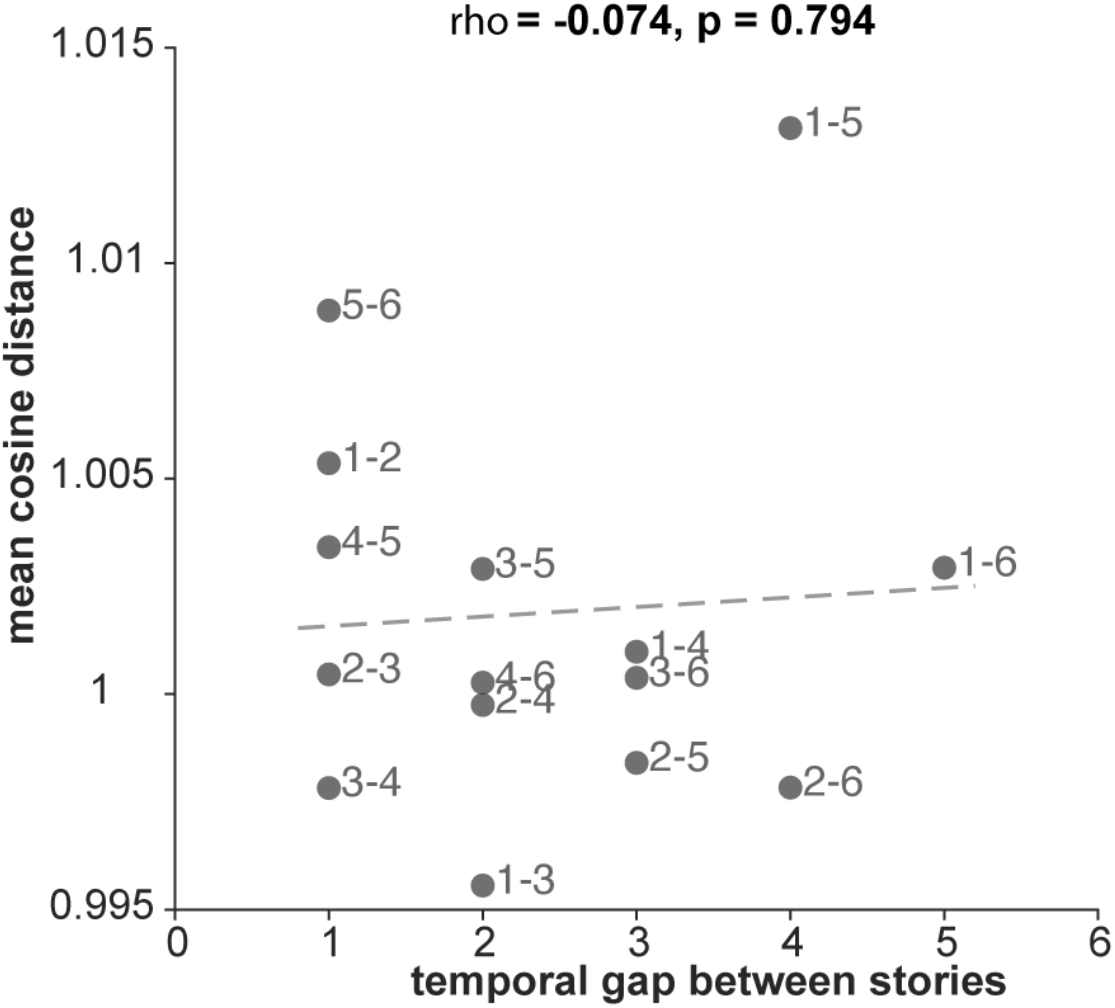
Across-story tuning distance is not explained by representational drift. Mean cosine distance between tuning vectors for each story pair (n = 15 pairs) plotted against the temporal gap between stories (ordinal distance). Tuning distance did not correlate with temporal separation (Spearman rho = -0.074, p = 0.794), indicating that the increased across-story tuning distance (Figure 4A-C) reflects context-dependent semantic retuning rather than gradual drift in neural representations over time. Related to Figure 4.

**Figure S12.**
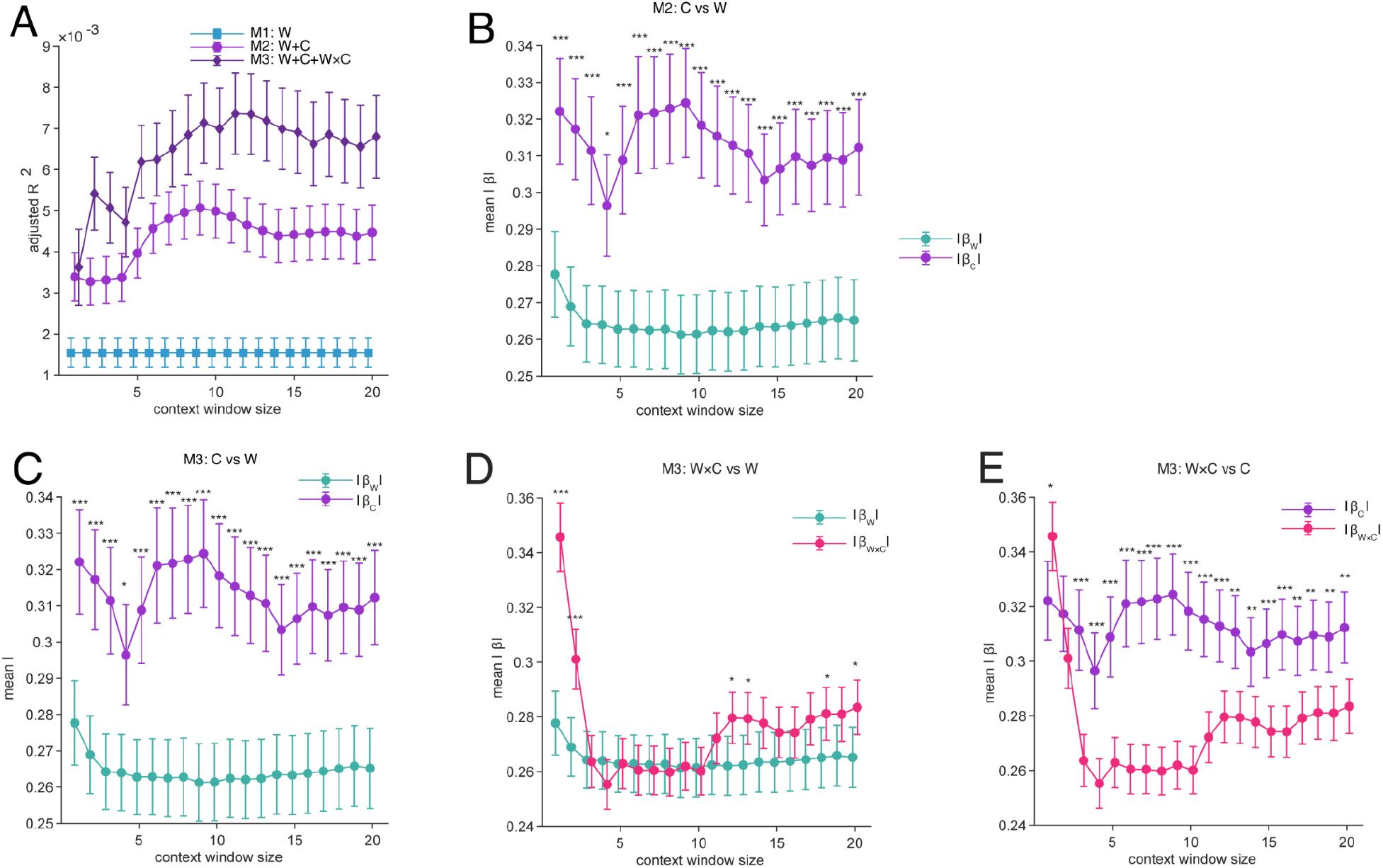
Lexical context modulation of semantic tuning across context window sizes. (A) Mean adjusted R^2^ (±SEM) for three nested regression models across 498 hippocampal neurons as a function of context window size (1-20 preceding words). M1 (blue) includes word embeddings only; M2 (light purple) adds context embeddings; M3 (dark purple) further adds word×context interaction terms. Both M2 and M3 significantly outperform M1 at all window sizes (all p < 0.001), and M3 significantly outperforms M2 at 19 of 20 window sizes (all p < 0.05, except window size=1). (B) Mean absolute standardized beta coefficients (±SEM) for word (|β_W_|, teal) and context (|β_C_|, purple) predictors in M2. Context coefficients are significantly larger than word coefficients at all window sizes (all p < 0.05). (C) Same comparison in M3. The pattern replicates: |β_C_| > |β_W_| across all window sizes (all p < 0.05). (D) Comparison of interaction (|β_W×C_|, pink) and word (|β_W_|, teal) coefficients in M3. The two are comparable in magnitude at windows 3-11 (all p > 0.05), with interaction coefficients significantly exceeding word coefficients at larger windows (window size >=19 and first 2 window sizes). (E) Comparison of interaction (|β_W×C_|, pink) and context (|β_C_|, purple) coefficients in M3. Context main-effect coefficients consistently exceed interaction coefficients at windows 3-20 (all p < 0.05). Together, these results establish a magnitude hierarchy of |β_C_| > |β_W×C_| ≥ |β_W_|, indicating that local lexical context is the dominant driver of hippocampal firing rate variance, while its nonlinear modulation of word encoding constitutes a secondary but substantial influence that exceeds the contribution of static word identity alone across the big context window sizes. * *p* < 0.05; ** *p* <0.01; *** *p*<0.001 (paired t-test, n = 498 neurons). Related to Figure 4.

**Figure S13.**
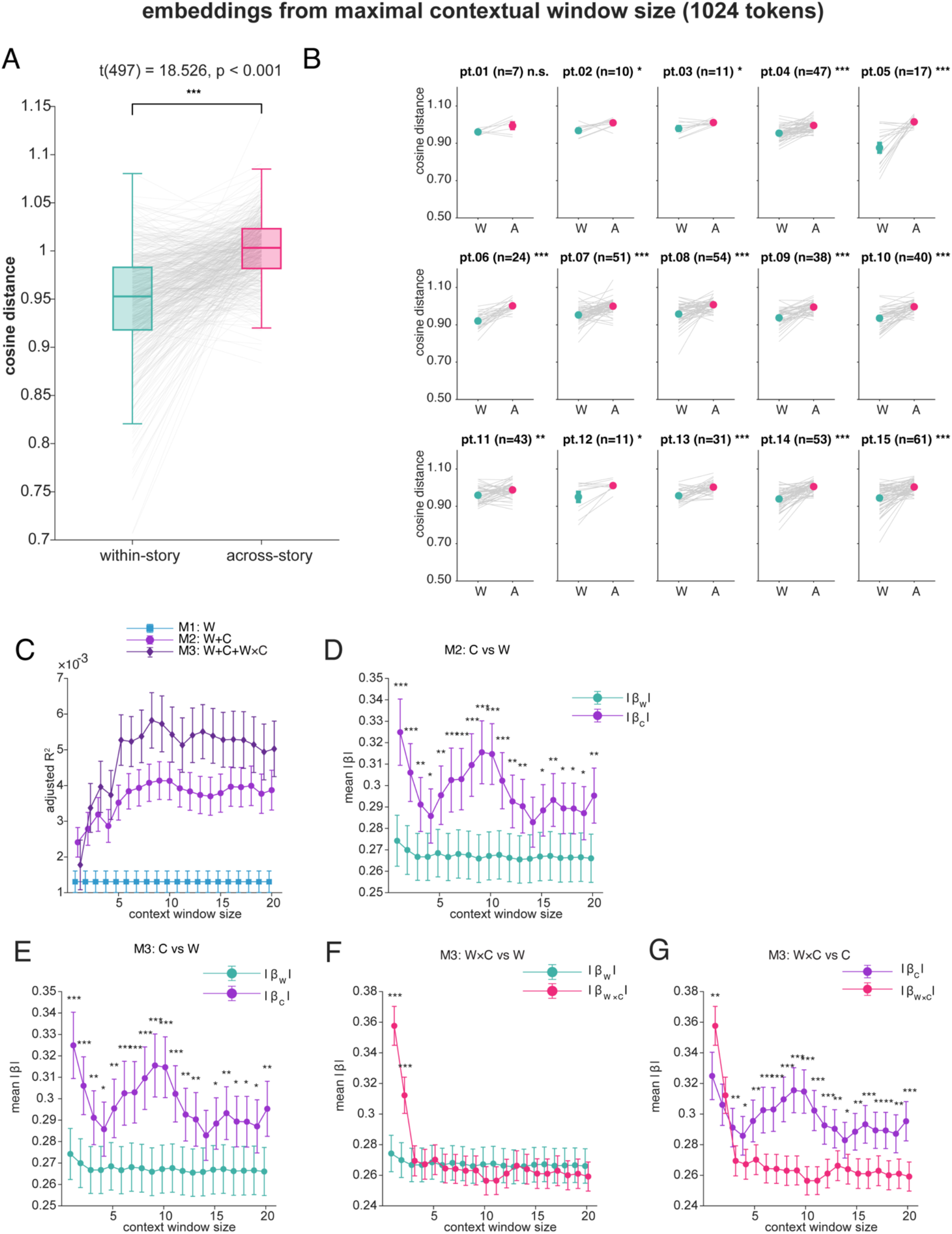
Control analyses for Figure 4A-C and Figure 4G-K. All panels repeat the analyses of Figure 4 with GPT-2 embeddings extracted using the model’s maximal context window, in which each word’s embedding incorporates the full preceding story rather than only its own sentence (Methods). (A-B) Replication of Figure 4A-C: across-story tuning distances remain significantly larger than within-story distances at the group level (A) and in the majority of patient (B). (C-G) Replication of Figure 4G-K: the nested model comparison, the ordering of coefficient magnitudes, and the interaction-neuron results are preserved across all 20 context window sizes. Related to Figure 4.

**Figure S14.**
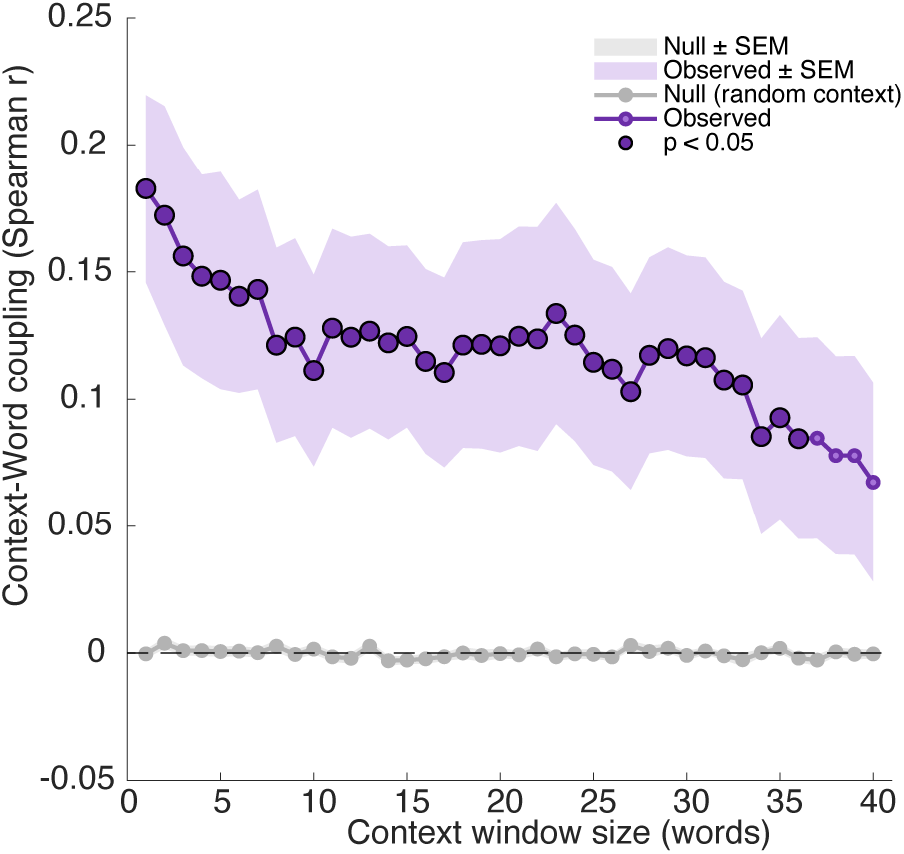
Control analysis for Figure 5D (pattern separation) To address the window-size question directly, we extended the window from 20 to 40 preceding words. Context-word geometry coupling significantly exceeded the random-context null at every window from 1 to 36 words (all p < 0.05, Wilcoxon signed-rank). Coupling was strongest for short windows (observed ρ ≈ 0.15 - 0.17 for 1-7 words), declined gradually and became non-significant beyond 36 words (p = 0.08 - 0.18 for 37-40 words). Lexical context therefore shapes the response to a repeated word over roughly the preceding 35 words, with the strongest influence from the most recent few, and the 1-20-word range reported in the manuscript falls well within this span. Related to Figure 5.

**Figure S15.**
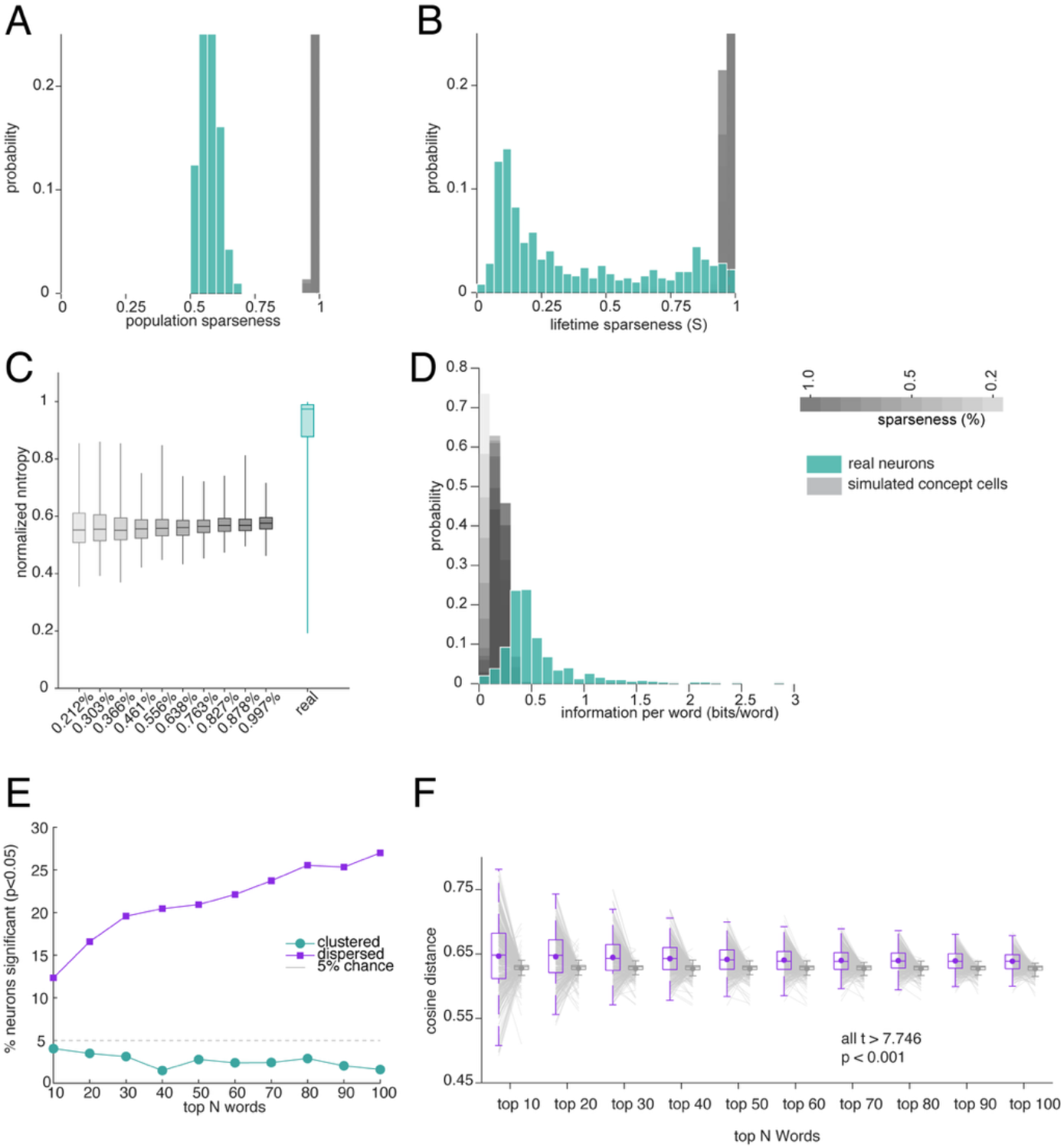
Dense coding and semantic overdispersion replicate with narrower response windows. To control for the possibility that the 40 ms post-offset extension of our response window captures early responses to the subsequent word, thereby inflating apparent polysemanticity, we repeated key analyses to support polysemanticity using a narrower window (80 ms post-onset to word offset, with no post-offset extension). (A) Population sparseness. Real neurons (teal) remain substantially denser than simulated concept cells (gray). (B) Lifetime sparseness. Real neurons remain denser than simulated concept cells. (C) Normalized entropy. Real neurons cluster at the high-entropy end, consistent with distributed coding. (D) Information per word. Real neurons carry more information per word than simulated concept cells. (E) Proportion of neurons showing significant dispersion vs. clustering across top N words. Overdispersion dominates at all N values. (F) Mean cosine distance for top N words vs. random N words. Top-preferred words remain significantly more dispersed than chance (all t > 7.746, p < 0.001). Related to Figure 2 and Figure 3.

### Supplementary Note 1. Coding-density metrics for verbs and adjectives

All four coding-density analyses reported for nouns in the main text (Figure 2) were repeated for verbs and adjectives, using the same preprocessing (lemmatization, token averaging per unique word) and the same ten simulated concept-cell populations (sparseness 0.20%-1.0%). Results closely matched those for nouns. Population sparseness was moderate (verbs: mean ± SD = 0.576 ± 0.030; adjectives: 0.581 ± 0.033; nouns: 0.581 ± 0.031), and lifetime sparseness was low (verbs: 0.364 ± 0.306; adjectives: 0.379 ± 0.306; nouns: 0.369 ± 0.305), indicating a dense code at both the population and single-neuron level. Normalized entropy was high (verbs: 0.899 ± 0.146; adjectives: 0.885 ± 0.165; nouns: 0.910 ± 0.130), and neurons carried substantial information per word (verbs: 1.707 ± 1.025 bits/word; adjectives: 1.916 ± 1.162; nouns: 0.464 ± 0.289). For every metric and every sparseness level, real neurons differed from simulated concept cells in the same direction as for nouns (Wilcoxon rank-sum tests, all p < 0.001, FDR-BH corrected; also see related results in Table S2 and Figure S6).

### Supplementary Note 2. Results of proper nouns

In our stimuli, we do have the proper nouns, including “Applebee’s”, and “Texas”, “Montana”, “Houston,” “Vietnam,” “Jesus”. We repeated the coding-density analyses of **Figure 2** restricted to the 66 unique proper nouns in the corpus. Hippocampal responses to proper nouns were indistinguishable from responses to the full noun set: lifetime sparseness, 0.365 ± 0.302 (mean±std; all nouns: 0.369 ± 0.305), population sparseness 0.588 ± 0.030 (all nouns, 0.581 ± 0.031), normalized entropy 0.868 ± 0.188 (all nouns: 0.910 ± 0.130). Information per word was higher for proper nouns (0.93 ± 0.84bits/word (all nouns 0.464 ± 0.289), but still far from the simulated concept cells (see **Figure 2**). Thus the named entities that most closely resemble classic concept-cell stimuli (people, places, brands) are not encoded more sparsely than common nouns in our data.

### Supplementary Note 3. Self-similarity of population responses to repeated and semantically related words

A sparse, stochastic, or highly variable code would produce population responses whose similarity does not depend on lexical identity or meaning. To test this, we exploited the natural repetition structure of the narratives rather than a designed repetition paradigm, which would compromise naturalness.

The 1221 nouns comprised 516 unique noun concepts. For each pair of words, we computed the Pearson correlation between the two population firing-rate vectors (498 neurons; firing rates z-scored per neuron across all words). Same-concept pairs were restricted to words separated by at least 100 words to exclude temporal autocorrelation, yielding 175 concepts with at least one valid pair (4,051 same-concept pairs). All remaining 131,208 pairs were different-concept pairs. Semantic distance between concepts was defined as the cosine distance between fastText embeddings. Semantically close pairs were defined as the bottom 2% of semantic distances (n = 2,624; e.g., anybody-anyone, kind-sort, daughter-son, brother-sister; note that this set includes contrast pairs as well as synonyms), and semantically distant pairs as a size-matched sample from the top 2%. Same- versus different-concept similarity was compared with a paired t-test across the 175 concepts (each concept’s mean same-concept correlation vs. its mean correlation with all other concepts). The relationship between neural similarity and semantic distance across different-concept pairs was assessed with a Spearman correlation, with significance from a permutation test (500 shuffles of concept identity), and with a multiple regression including temporal lag between words as a covariate.

Population responses to repetitions of the same concept were more similar than responses to different concepts (mean pairwise correlation 0.0128 vs. -0.0021; t(174) = 4.29, p = 2.9 × 10^-5^, Cohen’s d = 0.32). Among different concepts, neural similarity declined with semantic distance (Spearman ρ = -0.017, permutation p = 0.002, n = 131,208 pairs), and this relationship held after controlling for temporal lag (β_semDist = -4.3 × 10^-4^, t(131,205) = -3.64, p = 2.7 × 10^-4^). Semantically close pairs evoked more similar responses than semantically distant pairs (0.0006 vs. -0.0027; Wilcoxon rank-sum z = 3.54, p = 4.0 × 10^-4^; n = 2,624 per group). Similarity was thus ordered identical concept > semantically close > semantically distant.

This ordering is what a distributed semantic code predicts: each word occupies a reproducible location in population space (the basis of the decoding in Figure 1E-F), and nearby words occupy nearby locations (Figure 1H-M). It is incompatible with responses dominated by sparse, stochastic, or variable spiking, which would yield no dependence of similarity on identity or meaning. Individual pairwise correlations are necessarily small because each reflects a single-trial comparison in naturalistic data; the critical evidence is the systematic ordering. Note that this analysis tests signal versus noise and does not by itself distinguish concept-cell from polysemantic coding, since both predict reproducible word-specific population responses; that distinction is made at the single-neuron level (Figures 2-3). The residual variability across repetitions of the same word is itself structured by preceding lexical context (Figure 5A-D).

### Supplementary Results

To test whether our findings generalize beyond the story-listening dataset, we repeated the key analyses on recordings from six participants who listened to a 46-min dialogue (see **Methods**), yielding 217 hippocampal neurons and 1226 nouns (486 unique nouns). Simulated concept cells were matched to this dataset in neuron count, word count and word durations, at ten sparseness levels spanning 0.2%-1.0% (Methods). Real neurons used a far denser code than simulated concept cells on every measure (Wilcoxon rank-sum tests, all p < 0.001 at every sparseness level): population sparseness per word was 0.462 ± 0.051 (mean ± SD) versus 0.986-0.994 for simulations; lifetime sparseness per neuron was 0.343 ± 0.276 versus 0.972-0.981; normalized entropy was 0.916 ± 0.134 versus 0.555-0.571, consistent with distributed coding; and real neurons nevertheless carried more information per word (0.613 ± 0.274 bits/word versus 0.079-0.221), confirming that their broad tuning is informative rather than noisy. Semantic overdispersion also replicated: for every top-N value (N = 10-100), significantly more neurons showed dispersion than clustering among their top-preferred words, and the mean cosine distance among top-N words exceeded that of random N-word sets at all N (all t > 9.449, p < 0.001). The second dataset therefore reproduces both the dense coding profile (Figure 2) and the overdispersed semantic tuning (Figure 3).

